# Distinct Cellular Phenotypes of Language and Executive Decline in Amyotrophic Lateral Sclerosis

**DOI:** 10.1101/2025.02.26.640433

**Authors:** Joana Petrescu, Cláudio Gouveia Roque, Christopher A Jackson, Aidan Daly, Kristy Kang, Obadele Casel, Matthew Leung, Luke Reilly, Jacqueline Eschbach, Karina McDade, Jenna M. Gregory, Richard Bonneau, Colin Smith, Hemali Phatnani

## Abstract

Cognitive manifestations, including impairment in language and executive functions, are seen in amyotrophic lateral sclerosis (ALS), but the mechanisms that underlie these deficits remain unclear. To address this, we mapped prefrontal cortex regions from ALS patients by integrating spatial and single-nucleus transcriptomics in a cognitively stratified patient cohort. We uncover that cognitive impairment in ALS is associated with distinct patterns of neuronal dysfunction and glial-vascular dysregulation that vary by region and cognitive subtype. Executive dysfunction is linked to reduced mitochondrial and synaptic activity in neurons localized to the deeper layers of the dorsolateral prefrontal cortex, whereas language-related deficits track with a more diffuse, pan-regional response involving both glial and vascular abnormalities. Our analyses also identify signatures in the prefrontal cortex that span both motor and cognitive phenotypes, including a multicellular gliosis response. The findings reveal that the clinical heterogeneity of ALS is driven by phenotype-specific molecular and cellular interactions in motor and non-motor regions of the brain.

## INTRODUCTION

Amyotrophic lateral sclerosis (ALS) is a fatal neurodegenerative disorder characterized by loss of motor neurons leading to progressive paralysis^1^. Half of all patients exhibit some degree of extra-motor involvement, and the clinical presentation exists on a spectrum from ALS cases with normal cognition and behavior to ALS with frontotemporal dementia (ALS-FTD)^2,3^. A mid-spectrum range, which encompasses about 35% of ALS patients, is defined by mild cognitive and behavioral manifestations, including deficits in language, a decline in executive function, and apathy^2,3^. Brain imaging and postmortem analyses have shown that these cognitive and behavioral profiles are predictive of underlying dysfunction in specific areas of the brain, highlighting potential regional susceptibilities to ALS pathophysiology^2,4–8^. It remains unclear, however, whether the clinical phenotypes observed in ALS stem from a shared root mechanism or emerge from brain region-specific interactions of cellular, molecular, neuropathological, and genetic factors.

Addressing this question requires an integrative, transdisciplinary approach that bridges clinical phenotyping and neuropathology with cell-level molecular profiling. In practice, however, studying the clinical heterogeneity of ALS has been challenging, partly due to the lack of detailed cognitive assessment as part of routine ALS care^2^. Additionally, while technological advances have begun to provide a multicellular understanding of ALS^9–13^, disease-related interactions and dependencies between cells remain incompletely understood. Here, we apply spatial transcriptomics and single-nucleus RNA-seq to a carefully phenotyped ALS patient cohort to interrogate how differences in cell types and intercellular relationships relate to cognitive impairment. We performed these analyses in two non-motor regions of the prefrontal cortex implicated in cognitive impairment in ALS: executive function-associated Brodmann area (BA) 46 in the dorsolateral prefrontal cortex^5,14–16^, and BA44/BA45, commonly known as Broca’s area, linked to speech production and language^5,16–18^.

We identified molecular signatures shared across the ALS spectrum as well as responses that diverge between cognitive subtypes. Fluency and language deficits were associated with widespread glial, neuronal, and vascular alterations. Executive impairment, by contrast, was linked to specific neuronal subpopulations localized to the deeper layers of BA46. Overall, our patient stratification approach combined with spatially resolved, transcriptome-wide profiling revealed detailed relationships between clinical phenotype, cellular dysfunction, and pathology. We also demonstrate that cognitive subtypes in ALS reflect changes in distinct pathways, providing evidence for the molecular and cellular complexity of the disease.

## RESULTS

### Study Overview and Cohort Demographics

Our study focuses on a unique cohort of ALS patients who underwent neuropsychological testing using the Edinburgh Cognitive and Behavioral ALS Screen (ECAS) to assess for cognitive impairment. The ECAS is a multidomain clinical assessment tool used to quantitatively evaluate cognitive domains commonly affected in ALS patients, including executive function, language, and verbal fluency^2,3,19^. The ECAS generates an aggregate score that is indicative of cognitive impairment as well as scores for individual cognitive domains that capture subtleties in cognitive presentation^19^. Cognitive deficits in ALS are associated with localized cerebral and subcortical dysfunction, and the profile of cognitive change in each patient is predictive of underlying regional abnormalities^2,4–8^. We leveraged this structure-function relationship to investigate molecular and cellular signatures linked to domain-specific deficits in ALS. For patients with executive dysfunction, we sampled from the dorsolateral prefrontal cortex (DLPFC), specifically Brodmann Area (BA) 46, which supports decision-making and other high-level executive processes^5,14–16^. For those predominantly exhibiting language impairment, we prioritized BA44/45 in the inferior frontal gyrus of the left hemisphere, commonly known as Broca’s area, given its strong links to language production^17^. Verbal fluency, which involves both BA44/45 and BA46 networks, was investigated in connection to both anatomical regions in patients exhibiting impairments in this domain^5,7,16^. Building on this sampling strategy, we combined capture-based spatial transcriptomics (ST) and single-nucleus RNA sequencing (snRNA-seq) to obtain an unbiased and comprehensive view of ALS-related changes in non-motor areas of the brain (Figure 1A).

**Figure 1:**
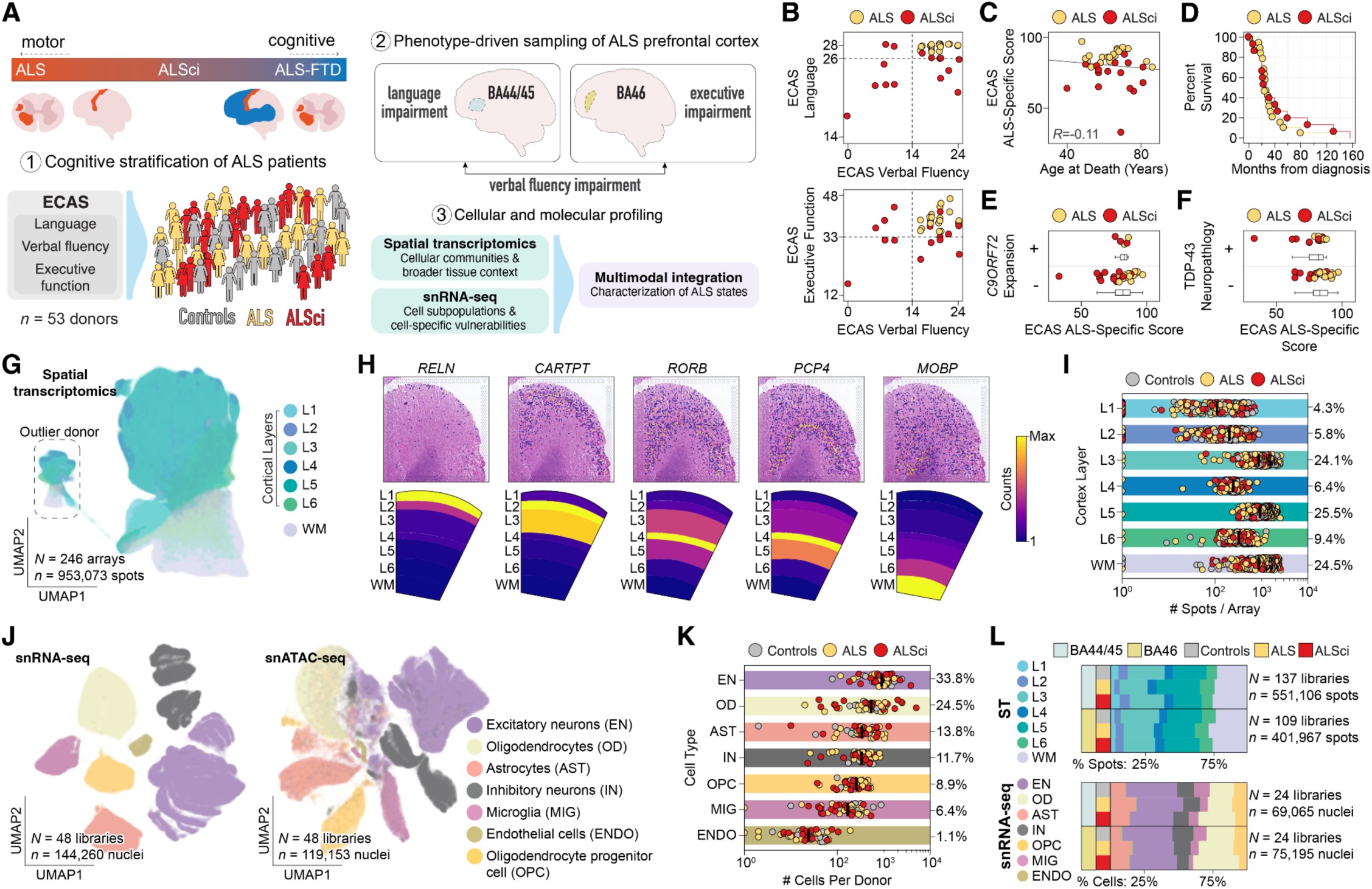
Study overview and cohort demographics. **A)** Summary of research strategy. We assembled an age- and sex-matched cohort of ALS donors (*n* = 36), along with sudden-death non-neurological controls (*n* = 17). ALS cases were cognitively stratified using the Edinburgh Cognitive and Behavioral ALS Screen (ECAS); donors with abnormal scores were classified as cognitively impaired (ALSci). Spatial transcriptomics and simultaneous capture of single-nucleus transcriptomes (snRNA-seq) and chromatin accessibility landscapes (snATAC-seq) from the same nuclei were performed on two prefrontal cortex regions: dorsolateral prefrontal cortex (BA46) and inferior frontal gyrus (BA44/45; Broca’s area). **B)** ECAS subtest scores (Executive Function, Verbal Fluency, and Language) distribution for each ALS donor, with dotted lines indicating the cutoff scores for impairment. **C)** Relationship between cognitive status and age at death across ALS donors. **D)** Kaplan-Meier plot showing disease duration (time from diagnosis to death) across ALS donors, stratified by cognitive status. **E)** Cohort stratification by *C9ORF72* genotype and cognitive phenotype. Five donors in the cohort carry an expanded *C9ORF72* allele. The ECAS ALS-specific score provides an aggregate measure of executive, fluency and language functions. **F)** Cohort stratification by presence or absence of TDP-43 neuropathology and cognitive phenotype. **G)** Uniform manifold approximation and projection (UMAP) of spatial transcriptomics data. Each data point represents a Visium array spot, color-coded by cortical layer annotation: layer 1 (L1) through layer 6 (L6) and white matter (WM). Seven arrays from a single donor, spanning both BA44/45 and BA46, separate from the rest of the dataset and were excluded from downstream analyses. These outlier data points are highlighted with a dashed box. **H)** Representative spot plots showing *RELN*, *CARTPT*, *RORB*, *PCP4*, and *MOBP* transcript expression overlaid on corresponding H&E-stained tissue section. Maximum expression is normalized per array for each transcript. The average spatial distribution of each marker gene is also displayed, with color indicating min-max scaled expression levels of each transcript across all Visium arrays. **I)** Number of spots annotated per cortical layer across all Visium arrays, color-coded by donor phenotype. The proportion of spots assigned to each layer is shown as a percentage, and the median spot count per layer is indicated with a black bar. **J)** UMAP projection of gene expression data from 48 snRNA-seq libraries (BA44/45, *n* = 24; BA46, *n* = 24), with cells colored by broad cell type. Corresponding UMAP projection of chromatin accessibility profiles from matched snATAC-seq libraries (BA44/45, *n* = 24; BA46, *n* = 24). **K)** Number of nuclei annotated per broad cell type across all snRNA-seq libraries, color-coded by donor phenotype. The proportion of cells assigned to each cell type is shown as a percentage, and the total number of nuclei per library reflects quality control and doublet removal. **L)** Distribution of annotated Visium spots across cortical layers (top) and broad cell types in snRNA-seq libraries (bottom), broken down by donor phenotype.

All ALS individuals in our study (*n* = 36) were participants in the CARE-MND initiative^20^, a Scotland-wide patient register unmatched in its depth, detail, and systematization of clinical information. This patient cohort was complemented by a cohort of age-matched sudden death donors that serve as non-neurological control subjects (*n* = 17)^21^. Postmortem brain tissue from ALS patients and control individuals was sourced from a single biorepository—the Edinburgh Brain Bank—to ensure procedural reproducibility across samples, including consistent preservation methods and standardized neuropathological assessments. Based on ECAS test scores, each donor was stratified into one of three analysis groups: ALS patients with cognition in the normal range (hereafter referred to as ‘ALS’), representing a pure motor form of the disease; ALS patients with cognitive impairment in one or more ECAS clinical domains (ALSci); and a third group consisting of neurotypical controls. We note that ALSci donors have mild cognitive impairment and do not meet the diagnostic criteria for ALS-FTD but rather fall within a mid-spectrum range on the ALS-FTD spectrum^2,3^.

We provide clinical phenotyping data and disease metrics for this unique cohort in Figures 1B-F and Supplemental Figure S1. Figure 1B details ECAS subtest scores for each ALS patient. Consistent with published literature^19,22^, language was the most frequently affected domain in ALSci donors, but the final cohort included patients with executive and fluency dysfunction (Figure 1B). We observed no correlation (*R* = -0.11) between donor age at death or disease duration with cognitive status (Figure 1C,D), and the number of donors was evenly distributed across clinical groups (Supplemental Figure S1A). The survival duration from disease onset was comparable between ALS and ALSci patients, although the ALSci group included a single long-term survivor (Figure 1D). We also sequenced donor genomes to determine *C9ORF72* repeat expansion status and performed semiquantitative TDP-43 assessments using immunohistochemistry. These analyses showed that neither *C9ORF72* genotype nor TDP-43 pathology predicted cognitive status in our cohort (Figures 1E,F and Supplemental Figures S1E,F).

### A Multimodal Cellular Atlas of the ALS Prefrontal Cortex

Spatially resolved transcriptome mapping of the prefrontal cortex was performed using the 10X Genomics Visium platform, which captures polyadenylated RNAs from intact tissue sections and localizes each RNA molecule to a specific 55-µm barcoded spot across a 6.5 x 6.5 mm capture area—sufficient to span all six cortical layers as well as adjacent white matter (Supplemental Figure S2A-F). We processed, sequenced, and analyzed 246 Visium arrays (BA44/45: *n* = 137, BA46: *n* = 109) from 45 individual donors. Each brain region was represented by at least two adjacent tissue sections, with most having higher technical replication (median of four arrays per region per donor). We obtained, in total, 953,073 discrete capture spots that passed quality control. For BA44/45, we detected an average of 3,447 unique RNA molecules aligned to 1,572 genes per spatial feature; similarly, we measured 3,679 unique RNA molecules aligned to 1,621 genes per spot in BA46 tissue (Supplemental Figure S2A-C). Each array spot was manually assigned to cortical layers L1-L6 and white matter based on histological characteristics extractable from hematoxylin and eosin (H&E)-stained brain tissue such as neuronal morphology and density. Individual array spots were preprocessed and embedded into a UMAP plot for visualization, showing separation of L1, white matter, and L2-L6 spots (Figure 1G). The quality of layer annotations was assessed using previously validated layer-specific markers; for example, L1 marker *RELN* was detected almost exclusively in this layer, while *MOBP*, a structural component of the myelin sheath, was strongly enriched in white matter (Figure 1H). Spot distribution across each layer of the prefrontal cortex was consistent between control and ALS donors (Figure 1I), suggesting no gross structural changes linked to ALS clinical phenotypes. These analyses also revealed a notable outlier, who clustered separately from the rest of the cohort in the UMAP plot and exhibited abnormally high expression of interferon and antiviral response genes (Supplemental Figures S2G,H). Given this atypical gene expression profile, this individual (an ALS patient without cognitive impairment) was excluded from all downstream analyses.

In parallel, we used 10X Genomics’ single-nucleus Multiome technology to obtain paired snRNA-seq and snATAC-seq profiles from nuclei isolated from 48 brain samples, evenly distributed across BA44/45 and BA46 regions (BA44/45: *n* = 24, BA46: *n* = 24). To aid in cell type annotation and ensure consistency with published molecular classification schemes, we also assembled a collection of 2.6 million prefrontal cortex snRNA-seq profiles from publicly available sources^9,12,23–29^ on which we trained a machine learning-based classifier to predict cell type annotations and remove low-quality or doublet cells (Supplemental Figure S3). After preprocessing and annotation (Supplemental Figure S4), we assigned 144,260 high-quality nuclei to seven broad CNS cell types: excitatory neurons (EN), inhibitory neurons (IN), oligodendrocytes (OD), oligodendrocyte progenitor cells (OPC), astrocytes (AST), microglia (MIG), and endothelial cells (ENDO) (Figure 1J). Neuronal and neuroglial populations were evenly represented, with excitatory neurons (33.8%), oligodendrocytes (24.5%), and astrocytes (13.8%) comprising the three most abundant cell populations (Figure 1K). All broad cell types were consistently detected across individuals and conditions (Figures 1K-L), and did not show substantial differences between ALS and control donors.

The integrative strategy we describe below leverages the complementary strengths of each modality—snRNA-seq for high-resolution cell type characterization and spatial transcriptomics (ST) for structural context across intact brain tissue. Both the results of snRNA-seq and cell-level deconvolution of Visium data with cell2location^30^ reveal broadly consistent cell compositions across clinical groups, though deconvolution of the spatial data suggested a marginal decrease in inhibitory neuron subtypes in the prefrontal cortex of ALS patients (Supplemental Figure S5). While orthogonal immunohistochemistry studies in donor tissue sections did not corroborate this decrease *in situ* (Supplemental Figure S6), analysis of snRNA-seq profiles revealed a general reduction in *PVALB*, *SST*, *VIP*, and *LAMP5* expression within each corresponding inhibitory neuron subtype (Supplemental Figure S5E,F), suggesting that cell2location detected changes in neuronal marker gene expression rather than cellular abundance. Finally, we note that a detailed exploration of the snATAC-seq data modality is beyond the scope of the current study and will be presented in a forthcoming publication.

### Glial Activation is a Frequent Feature of the ALS Prefrontal Cortex

We began by searching for signatures commonly seen across ALS patients, independent of cognitive status, as identifying such shared responses is critical for disentangling mechanisms specifically associated with cognitive impairment from broader disease processes. Among all cell types, endothelial cells globally showed the strongest transcriptional changes, with differential expression analysis between control and ALS donors revealing enrichment in gene ontology terms related to immune response, apoptosis, and angiogenesis (Figure 2A). However, endothelial cells represented only 1.1% of captured nuclei in the snRNA-seq dataset (Figure 1K), limiting our ability to resolve vasculature-associated subtypes or draw more detailed inferences using this modality. The undersampling of vascular cells we describe is consistent with other reports and reflects a well-recognized limitation of current protocols^29,31^.

**Figure 2:**
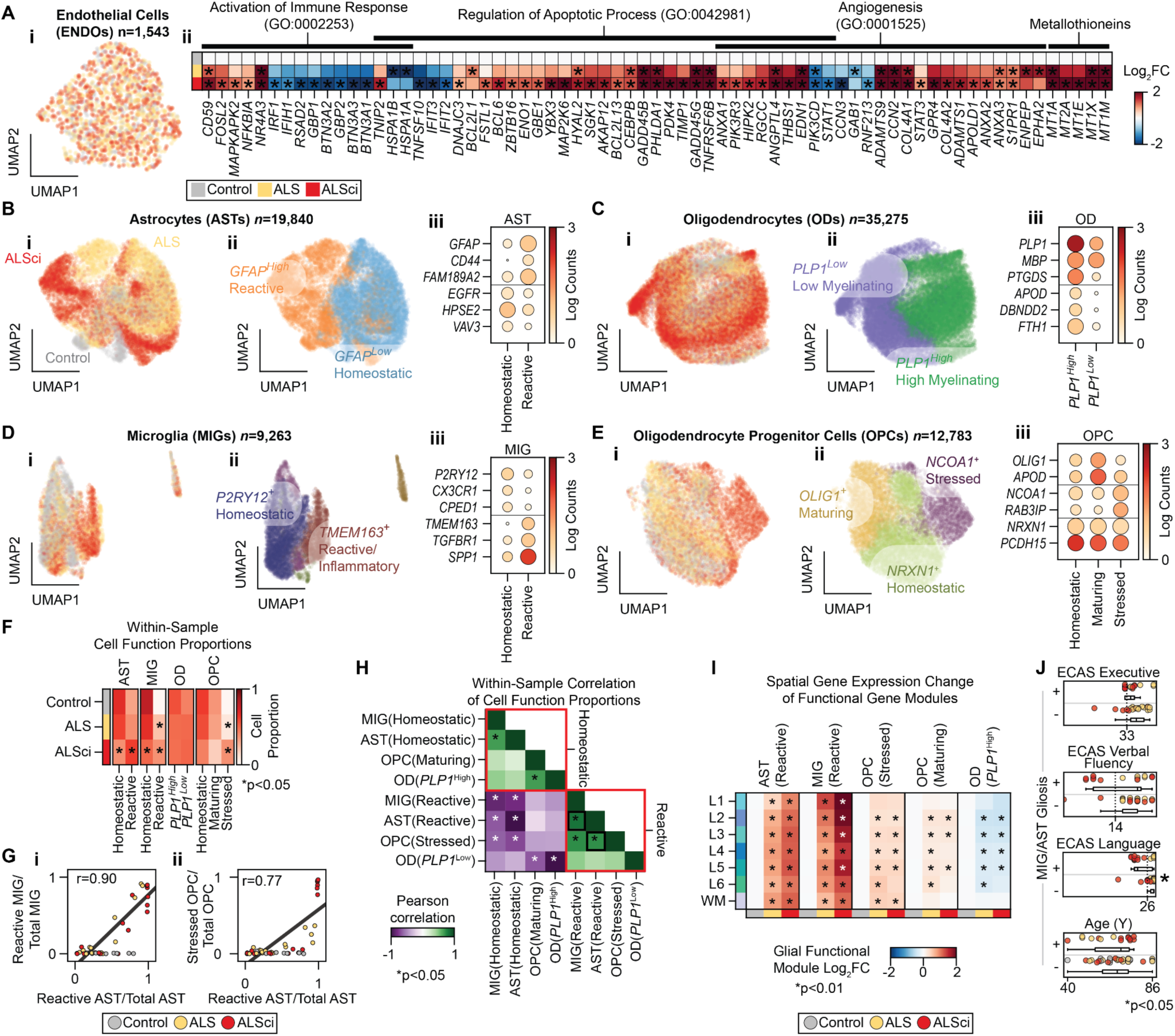
ALS donors with and without cognitive impairment have transcriptomic signatures of gliosis. **A)** Endothelial cells from snRNA-seq dataset projected into a UMAP plot and colored by donor phenotype (i) and heatmap showing representative pseudobulked snRNA-seq gene expression differences between control donors and ALS or ALSci donors, grouped into four selected ontology categories (ii). Asterisks indicate significant difference from Log_2_ fold change of 0 (DESeq2; multiple hypothesis test corrected Wald test; *p* < 0.05). **B)** Astrocytes from snRNA-seq projected into a UMAP plot colored by (i) donor phenotype, (ii) transcriptome-inferred functional states (homeostatic [*GFAP^Low^*] or reactive [*GFAP^High^*]), and (iii) specific functional marker genes. Marker gene dot plot is colored by mean gene expression and dot size indicates the proportion of cells with non-zero counts, from 0% to 100%. **C)** Classification of oligodendrocytes into low-myelinating (*PLP1*^Low^) and high-myelinating (*PLP1*^High^) functional states following the same plotting scheme as in B. **D)** Classification of microglia into homeostatic (*P2RY12*^+^), reacting/inflammatory (*TMEM163*^+^) functional states following the same plotting scheme as in B. Phagocytic, stress-response, and non-microglial immune cells are also plotted in (ii) without labels. **E)** Classification of oligodendrocyte progenitor cells into homeostatic (*NRXN1^+^*), maturing (*OLIG1^+^*), or stressed (*NCOA1^+^*) functional states following the same plotting scheme as in B. **F)** Averaged proportions of glial cell functional states across controls, ALS, and ALSci donors. Asterisks denote significant differences compared to controls (Welch’s *t*-test, *p* < 0.05). **G)** Proportion of activated-to-total microglia (i) and stressed-to-total OPCs (ii) plotted against proportion of reactive-to-total astrocytes. Dots represent individual donor samples and each dot is colored by donor phenotype (controls, grey; ALS, yellow; ALSci, red). Plots are also annotated with least squares regression line and Pearson correlation coefficient (*R*). **H)** Pearson correlation matrix of glial functional states proportions. Black boxes highlight correlations shown in G. Asterisks denote correlation significantly different from 0 (Wald test, *p* < 0.05, false discovery rate-corrected). **I)** Spatial distribution of glial functional module scores across Visium arrays, plotted as log_2_-transformed fold-change relative to controls. Asterisks denote significant differences relative to control donors (Welch’s *t*-test against, *p* < 0.01, false discovery rate-corrected). **J)** ECAS domain scores (executive, verbal fluency, and language) and donor age, grouped by presence (+) and absence (-) of transcriptional signatures of gliosis (defined as >80% reactive astrocytes or >20% reactive microglia). ECAS scores range from 0 to 48 (executive), 24 (fluency), and 28 (language). Vertical dashed lines denote impairment thresholds. Dots are colored by donor phenotype, and asterisks indicate significant group differences (Welch’s *t*-test, *p* < 0.05).

Additional preliminary observations turned our attention to glial cells, including the presence of broadly similar transcriptional alterations in both ALS and ALSci donors without significant changes in overall glial cell abundance compared to controls (Supplemental Figure S7). These findings suggested that the pathophysiology of ALS in non-motor regions involves transitions between functional states (e.g., the conversion of homeostatic microglia into a reactive state) rather than changes in the proliferation of glial cells, prompting a more detailed examination of glial subpopulations. Overall, glial cells were well represented in our snRNA-seq dataset (Figures 1J-L), and based on Leiden clustering, key marker gene expression, and cross-referencing with previous analyses, we annotated multiple functional states for each major glial cell type. For astrocytes, we distinguished homeostatic and reactive states based on characteristic transcriptional profiles and used these to classify nuclei into reactive or homeostatic astrocytes (Figure 2B and Supplemental Figure S8A). Oligodendrocytes were subclassified into high- and low-myelinating states based on expression of myelin-associated genes (Figure 2C and Supplemental Figure S8B). Microglia were divided into homeostatic and reactive/inflammatory states, with smaller subsets exhibiting phagocytic or heat shock-associated signatures (Figure 2D and Supplemental Figure S8C). In addition, a small population of non-microglial immune cells was also present among nuclei broadly classified as microglia and was annotated separately as ‘other immune’ (Supplemental Figure S8C). For OPCs, we classified nuclei into homeostatic and *OLIG1*^+^ maturing/differentiating states, along with a *NCOA1^+^* subpopulation with a distinctive transcriptional profile which we have named ‘stressed’ (Figure 2E and Supplemental Figure S8D).

Building on these annotations, we then performed compositional analyses to determine whether specific glial functional states differed between disease and control donors. We observed a significant increase in reactive microglia and stressed OPCs in both ALS and ALSci donors, while the proportion of reactive astrocytes was only significantly higher in ALSci donors (Figure 2F). Notably, these reactive cell states—reactive astrocytes, reactive microglia, stressed OPCs—were highly correlated within individual samples (e.g., a donor with more homeostatic astrocytes also tended to have more homeostatic microglia; Figure 2G and Supplemental Figure S8F), suggesting they are part of a coordinated gliotic response. In addition to this strong association among reactive populations, we also found that the abundance of certain homeostatic glial subpopulations correlated positively with one another (Figure 2H). This was particularly notable between homeostatic astrocytes and microglia, as well as between OPCs and high-myelinating oligodendrocytes. By contrast, homeostatic microglia and astrocytes did not correlate with maturing OPCs or myelinating oligodendrocytes (Figure 2H).

We next examined the spatial distribution of this shared gliotic response across intact tissue sections using our ST dataset. We first identified transcripts that are both specific to individual glial cell types and differentially expressed between functional states of that glial cell type (e.g., transcripts upregulated in reactive versus homeostatic astrocytes) (Supplemental Figure S8G). We then used these expression signatures to score each array spot, allowing us to quantify shifts in glial cell states between control and ALS donors. This analysis revealed that reactive astrocytes, reactive microglia, and stressed OPCs were significantly increased across layers L1-L6 and white matter in both ALS and ALSci donors (Figure 2I), indicating a diffuse, pan-layer response. In contrast, the transcriptional signature of myelinating oligodendrocytes declined in L2-L6 and the signature of maturing OPCs increased in L2-L6, while both were unchanged in white matter (Figure 2I). Together, the findings are consistent with the idea that glial responses in ALS are not only coordinated but also spatially organized.

While we detected gliosis signatures in both ALS and ALSci donors, they were not present in every patient (Figure 2G). These signatures were, however, generally more pronounced in ALSci (Figures 2F, 2G and 2I), raising the question of whether specific cognitive profiles are associated with this glial response. To test this, and to account for donor heterogeneity, we categorized each sample in the snRNA-seq dataset as either positive or negative for reactive microglia or reactive astrocytes (AST/MIG gliosis). ECAS domain scores were largely similar between both donor groups, with the exception of language domain, which was significantly lower in donors with reactive gliosis (Figure 2J). Our analyses also revealed that gliosis was not associated with donor age in our cohort (Figure 2J). These results suggest that while gliosis is a feature of ALS, it is unlikely to be the primary driver of cognitive dysfunction. Collectively, the findings suggest that cognitive decline in ALS is likely driven by additional mechanisms beyond the gliotic responses observed here.

### Spatial Expression Signatures Distinguish ALS Cognitive Subtypes

At the molecular level, the central nervous system is characterized by spatially correlated transcriptomic patterns that reflect cellular interactions^32,33^. Disruptions in these spatial expression programs have been linked to neurological conditions^34,35^, including in the ALS spinal cord^13^. To investigate ALS-related changes in non-motor areas of the brain in an unbiased manner, we employed Splotch^13,36,37^, a Bayesian hierarchical model designed for spatial transcriptomic data (Figure 3A). Splotch models gene expression at each spatial spot as a zero-inflated Poisson distribution, with spot-level **λ** (transcript abundance) informed by spot- and donor-level metadata (cortical layer and ALS diagnosis), spatial autocorrelation with neighboring spots, and spot-specific noise (Supplemental Figure S9A-C). After Bayesian inference, we quantified both the magnitude (log_2_ fold-change) and significance (Bayes factor) of gene expression changes between control and ALS/ALSci donors within each cortical layer through analysis of posterior distributions over model parameters (Supplemental Figure S9D-E; see Methods). Globally, the Poisson **λ** parameters from the fitted Splotch model—which represent denoised estimates of transcript expression at every array spot across the dataset—exhibited similar spatial correlation patterns in both BA44/45 and BA46, consistent with the shared laminar architecture and cell type composition of these prefrontal cortex regions (Supplemental Figure S10A).

**Figure 3:**
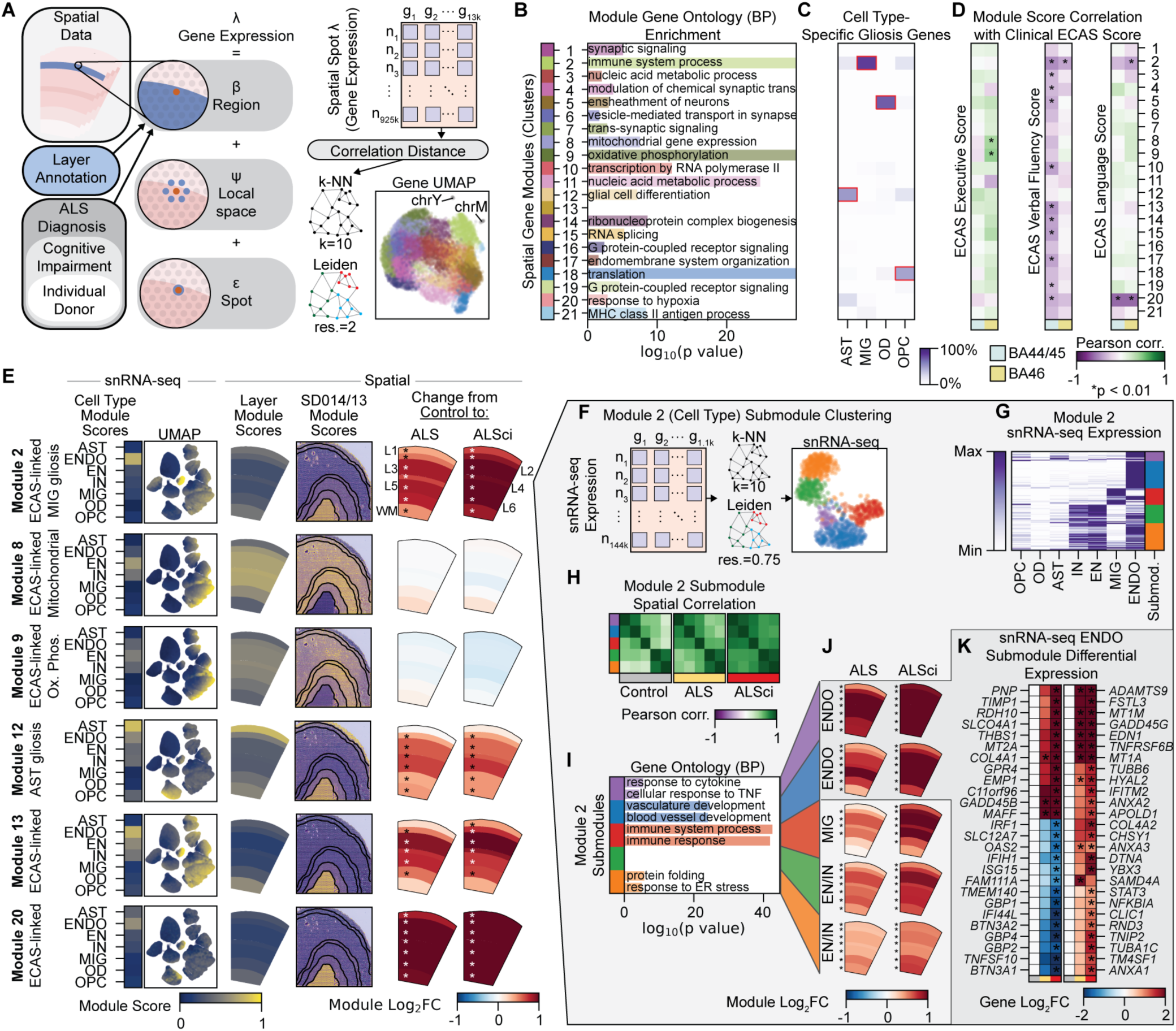
Spatial Expression Signatures Distinguish ALS Cognitive Subtypes. **A)** Schematic of the Splotch model fit to each gene captured in the spatial transcriptomic data set. Transcript expression (λ) at each spatial spot on each array is modeled as a combination of ALS disease state, cognitive impairment profile, and donor ID (β), a spatial autoregressive component (φ), and a spot noise (ε) component. Distance between genes is calculated as the correlation distance between modeled spot expression (λ), genes are embedded into a k-Nearest Neighbors (k-NN) graph, and clustered with the Leiden community clustering algorithm to identify spatially correlated gene modules. Gene-gene k-NN is embedded into a UMAP plot where each dot is a gene, colored by spatial gene module. **B)** Most significant biological process (BP) gene ontology terms associated with each spatial gene module. Bar length reflects -log_10_(*p*-value) of enrichment. Blank line indicates no GO:BP enriched terms for that spatial module. **C)** Heatmap showing spatial modules containing transcripts specific for reactive astrocytes and microglia, low-myelinating oligodendrocytes, and stressed OPCs. Color denotes the proportion of transcripts specific for the glial cell function that are assigned to each spatial module. Modules with >50% of function-specific transcripts are highlighted in red. **D)** Pearson correlation between donor-aggregated gene module and ECAS cognitive scores. Asterisks indicate correlations significantly different from zero (*p* < 0.01) by Benjaminini-Yekutieli corrected Wald test. **E)** Summary of selected spatial gene modules. From left to right, the average score of the spatial gene module from snRNA-seq expression data is plotted for each snRNA-seq cell type, and the snRNA-seq UMAP is plotted where each dot is an individual cell colored by the module score. Average score of the spatial gene module from spatial expression data is plotted for each cortex layer, and a representative example array is shown with module scores overlaid onto each spot. Differential module expression is plotted from spatial expression data, comparing control donors to ALS and ALSci donors. Asterisks indicate significant differences for a cortex layer, tested by Benjaminini-Yekutieli controlled Welch’s *t*-test (*p* < 0.01). **F)** Schematic illustrating the assignment of genes from spatial module M2 into submodules by correlation within snRNA-seq data. Module M2 gene-gene k-NN is embedded into a UMAP plot where each dot is a gene, colored by submodule. **G)** Average expression of each gene in module M2 across cell types in snRNA-seq. Expression values are min-max scaled per gene for visualization, and submodule identity is plotted on the right. **H)** Pearson correlation between module M2 submodule scores, stratified by clinical phenotype. **I)** Most significantly enriched gene ontology terms (biological process) associated with each module M2 submodule. Bar length reflects -log_10_(*p*-value). Blank line indicates no GO:BP enriched terms for that spatial submodule. **J)** M2 submodule expression in ALS and ALSci donors as log_2_ fold change relative to controls, plotted by cortical layer. Asterisks indicate layer-specific significant differences, tested by Benjaminini-Yekutieli controlled Welch’s *t*-test (*p* < 0.01). **K)** Heatmap of pseudobulked snRNA-seq gene expression changes, showing representative differentially expressed endothelial genes from endothelial M2 submodules as Log_2_ fold change over control donors. Asterisks indicate significant difference from Log_2_ fold change of 0 (DESeq2; multiple hypothesis test corrected Wald test; *p* < 0.05).

Beyond identification of individual genes that exhibit spatial differential expression, we sought to identify groups of spatially co-expressed genes. To this end, we embedded each transcript into a k-Nearest Neighbors (k-NN) graph, using Pearson correlation distance, and clustered them with the Leiden graph clustering algorithm into co-expressed spatial modules (Figure 3A, Supplemental Figure S8B). This approach yielded 23 spatial co-expression modules, each representing a set of genes co-expressed in overlapping spatial patterns across the cortex, with the exception of two capturing chromosome-specific gene expression (from the Y-chromosome [chrY] and mitochondria [chrM], respectively), which were excluded from subsequent analyses (Figure 3A, Supplemental Figure S10C).

We functionally characterized the remaining 21 spatial modules based on gene ontology (Figure 3B), finding two modules linked to immune response (M2 and M21), one module linked to myelination (M5), two modules linked to mitochondria (M8 and M9), and several linked to synapse or receptor signaling (M1, M4, M6, M7, M16 and M19). Also of note, M2, M12, and M20 were enriched for transcripts associated with reactive astrocytes and microglia (Figure 3C), consistent with the gliotic response we have characterized above (Figure 2). These analyses suggest that the gene co-expression modules we identified are structured by both spatial and functional relationships. To further explore these associations, we calculated module scores, which represent the aggregate expression of all transcripts comprising each module for each Visium spot. Because our dataset includes cognitive domain-specific scores and samples from two associated cortical regions, we were able to correlate spatial co-expression module scores from BA44/45 and BA46 with specific cognitive impairments and localize their regional specificity (Figure 3D). Notably, we observed distinct spatial correlations that correspond to executive, language, and verbal fluency functions (Figure 3D). Impairment in executive function correlated with downregulation of M8 and M9 in BA46, two modules associated with mitochondrial function, particularly oxidative metabolism (Figure 3B,D). This contrasted with impairment in verbal fluency, which correlated predominantly with upregulation of several spatial modules in BA44/45 (Figure 3D). Deficits in language function correlated with an upregulation of gliosis-linked modules M2 and M20 in both BA44/45 and BA46 (Figure 3C,D).

To explore the association between spatial modules and cellular processes, we similarly calculated module scores within our snRNA-seq data and determined the strongest cell type associations for each module (Figure 3E, Supplemental Figure S10B). We also evaluated module scores across cortical layers and sought to identify disease-related changes by comparing control to ALS and ALSci donors (Figure 3E, Supplemental Figure S10D). Because Visium spots often sample multiple cells, our spatial co-expression modules can be associated with multiple distinct cell types. We therefore further separated them into submodules to obtain greater cell type specificity. For each module, module transcripts were embedded into a k-NN graph using snRNA-seq transcript correlation distance as a metric and reclustered into submodules (Figure 3F, Supplemental Figure S10D). We selected spatial module M2 for further exploration as it was upregulated in ALS and ALSci donors and associated with both language and verbal fluency dysfunction (Figure 3D-E). After separating M2 into submodules (Figure 3F), we found transcriptional signatures related to endothelial, microglial, and neuronal nuclei. Notably, we observed a progressive increase in spatial cross-correlation between M2 submodules from control to ALS to ALSci donors, suggesting that cognitive impairment is linked to the disruption and an intermixing of cellular processes that are typically spatially segregated (Figure 3H, Supplemental Figure S11). Gene ontology (GO) analyses further revealed that specific M2 submodules were enriched for microglial immune responses, endothelial vascular development, and neuronal protein folding functions (Figure 3I). Overall, these submodules were upregulated in ALS donors, with the effect generally more pronounced in ALSci individuals (Figures 3J,K). These spatially coordinated vascular, neuronal, and glial changes implicate disrupted intercellular interactions as key contributors to language and fluency decline in ALS.

### Executive Impairment is Linked to BA46 Deep-layer Neurons

While many spatial gene modules (e.g., M2) showed links to verbal fluency and language deficits in ALS patients, a smaller and distinct set correlated with executive dysfunction, suggesting that there are different mechanisms associated with impairment in this cognitive domain. To further dissect the molecular basis of executive deficits in ALS, we focused on M8 and M9—the only spatial modules whose expression significantly correlated with ECAS executive test scores (Figure 3D). These two modules, which are enriched for mitochondria-associated genes and linked to both excitatory and inhibitory neurons (Figure 3B, Supplemental Figure S10D), were downregulated in ALS patients with executive impairment, specifically in BA46 (Figure 3D). When we stratified spatial module correlations by cortical layer, this response was found to be localized to the deeper layers (L5 and L6) of the gray matter (Figure 4A), suggesting that executive deficits in ALS are tied to dysfunction in spatially distinct neuronal subpopulations. These findings prompted a more detailed investigation of the cellular and molecular underpinnings of executive dysfunction in ALS.

**Figure 4:**
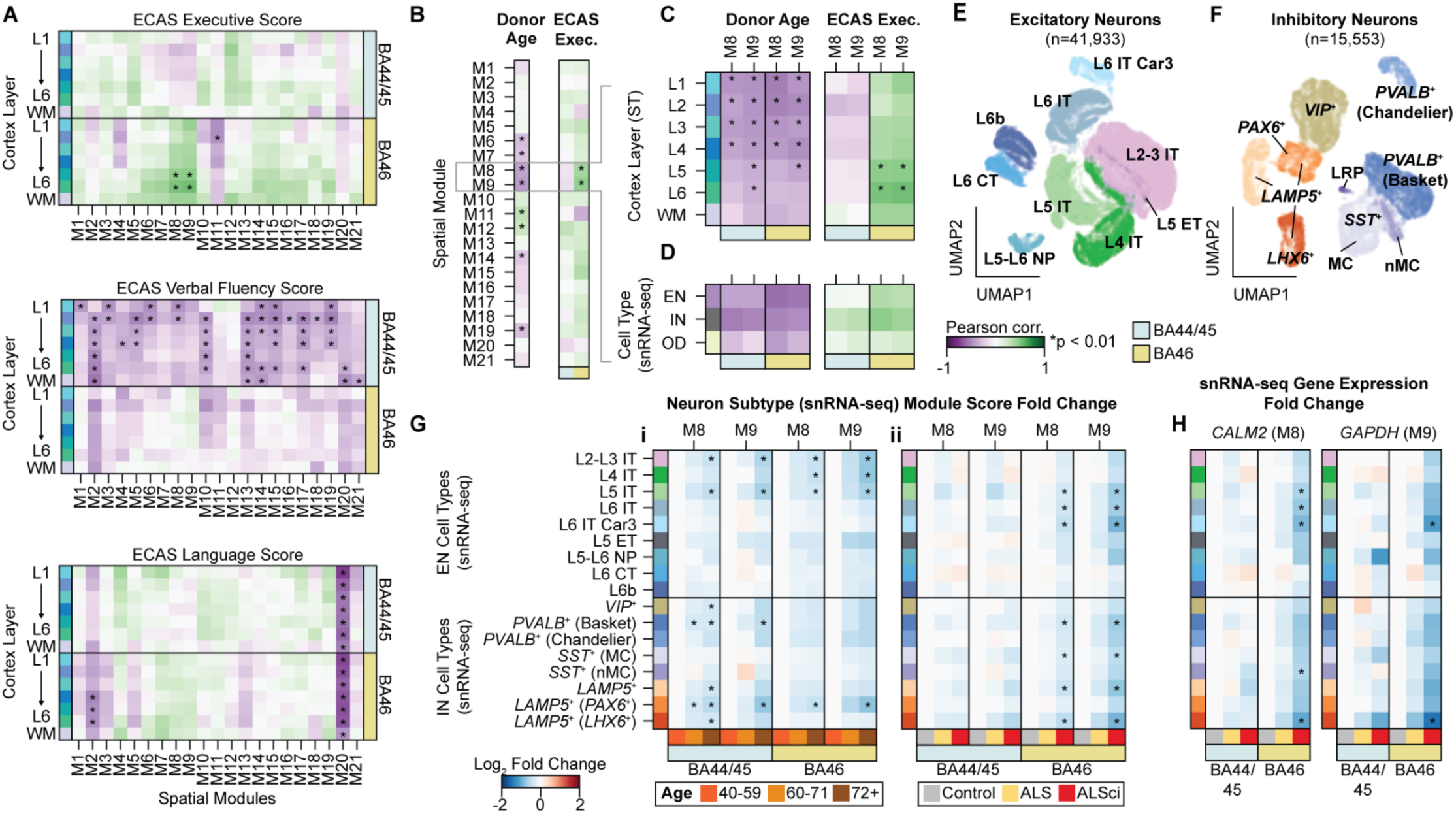
Executive Impairment is Linked to BA46 Deep-layer Neurons. **A)** Pearson correlation between ST co-expression module scores and ECAS domain scores (executive, verbal fluency, and language), stratified by brain region and cortex layer. Asterisks indicate correlations significantly different from zero (*p* < 0.01) by Benjaminini-Yekutieli corrected Wald test. **B)** Pearson correlation between ST co-expression module scores and donor age, compared to correlation between spatial module and ECAS executive score. Asterisks indicate correlations significantly different from zero (*p* < 0.01) by Benjaminini-Yekutieli corrected Wald test. **C)** Pearson correlation between spatial module M8 and M9 scores and donor age, compared to correlation between these module scores and ECAS executive score, stratified by brain region and cortical layer. Asterisks indicate correlations significantly different from zero (*p* < 0.01) by Benjaminini-Yekutieli corrected Wald test. **D)** Pearson correlation between module M8 and M9 scores (here calculated from snRNA-seq data) and donor age, compared to the correlation between these module scores and ECAS executive score, stratified by brain region and broad cell type. Correlations significantly different from zero (*p* < 0.05) by Benjamini-Hochberg corrected Wald test would be indicated with asterisks (none are significant). **E)** UMAP plot of excitatory neurons, colored by neuronal subtype. Excitatory subtypes include L2/L3 IT, L4 IT, L5 IT, L5 ET, L6 IT, L6 IT Car3, L5/L6 NP, L6 CT, and L6b neurons. **F)** UMAP plot of inhibitory neurons, colored by neuronal subtype. Inhibitory subtypes include *VIP*^+^, *LAMP5*^+^, *LAMP5*^+^ (*LHX6*^+^), *LAMP5*^+^ (*PAX6*^+^), *PVALB*^+^ Chandelier, *PVALB*^+^ Basket, *SST*^+^ Martinotti cells (MC), *SST*^+^ non-Martinotti cells (nMC), and *SST*^+^ Long-range Projection (LRP) neurons. **G)** Log_2_ fold change of module M8 and M9 scores calculated from snRNA-seq data, stratified by brain region and neuronal subtypes. (**i**) Comparison of donors from older age groups (60–71 years and 72+ years) to a younger group (40–59 years). Age categories were defined to stratify donors into three equal-sized groups. (**ii**) Comparison of ALS and ALSci donors to control donors. Asterisks indicate fold changes significantly different from zero (*p* < 0.05) by Benjamini-Hochberg corrected Welch’s *t*-test. **H)** Log_2_ fold change of *CALM2* (the highest expressed module M8 transcript) and *GAPDH* (the highest expressed module M9 transcript) from snRNA-seq dataset, stratified by brain region and neuronal subtype, and comparing ALS and ALSci donors to controls. Asterisks indicate fold changes significantly different from zero (*p* < 0.05) by Benjamini-Hochberg corrected Welch’s *t*-test.

Given that executive functions are commonly affected by aging^38,39^, we first examined the influence of this clinical variable on modules M8 and M9 to distinguish age- and disease-driven executive impairment. Interestingly, donor age did correlate with several spatial modules, including executive dysfunction-linked M8 and M9 (Figure 4B). Specifically, both M8 and M9 showed decreased expression in older donors, as well as in those with lower ECAS executive scores (Figure 4B). However, the spatial location of these age- and executive-linked changes in the ST data was not the same (Figure 4C). M8 and M9 exhibited significant anti-correlation with age in layers L1-L5 of both BA44/45 and BA46 samples (Figure 4C). In contrast, the same modules showed significant correlation with ECAS score in layers L5-L6 of BA46 samples alone. This highlights the importance of integrating spatial profiling in transcriptomic analyses, as perturbations to the same molecular processes (e.g., modules M8/M9) in differing spatial patterns may help explain differing associations with related clinical variables such as age-driven and disease-driven cognitive decline.

While we have investigated correlations between spatial modules and broadly annotated excitatory and inhibitory neurons, we next sought to determine if specific *sub*populations of neurons were being affected. From our snRNA-seq dataset, we separately clustered excitatory and inhibitory neurons, and assigned each cluster a neuronal subtype based on the expression of key marker genes (Supplemental Figure S12)^40^. We identified nine well-characterized EN subtypes, including L2/3, L4, L5, L6, and L6 Car3 intratelencephalic (IT) neurons, L5 extratelencephalic (ET) neurons, L5/L6 near-projecting (NP) neurons, L6 corticothalamic (CT) projection neurons, and L6b neurons (Figure 4E). For INs, we identified nine well-characterized subtypes, including *VIP*^+^, basket and chandelier *PVALB*^+^, three *LAMP5*^+^ subtypes, and three *SST*^+^ interneuron subtypes (Figure 4F). We annotated *SST*^+^ clusters as Martinotti (MC), non-Martinotti (nMC), and long-range projection (LRP) interneurons based on expression of comparable mouse marker transcripts (Figure 4F)^41^. Globally, the relative proportion of excitatory and inhibitory neuronal subtypes was not significantly different between control, ALS, and ALSci donors (Supplemental Figures S12C,E).

After classifying individual nuclei in our snRNA-seq dataset, we compared spatial module scores between three age groups, and between control and ALS donors for each neuronal subtype (Figure 4G). In line with our previous layer-level analysis (Figure 4C), M8 and M9 showed decreased expression in older donors in L2-L3, L4, and L5 IT ENs, and in L1 *LAMP5*^+^/*PAX6*^+^ INs, and did so comparably in both BA44/45 and BA46 samples (Figure 4Gi). Consistent with the executive impairment-associated downregulation of modules M8 and M9 in the deeper layers of BA46, we found that L5 IT, L6 IT, and L6 IT Car3 ENs exhibited significant downregulation of M8 and M9 genes (Figure 4Gii). Likewise, L5-L6 *LAMP5*^+^/*LHX6*^+^ and L5 *PVALB*^+^ basket inhibitory neurons also showed decreased M8 and M9 gene expression in ALSci donors (Figure 4Gii). In contrast, M8- and M9-linked transcripts were not significantly affected in other neuron subtypes located to L5 or L6, including L5/L6 NP, L6b, and L6 CT ENs, nor in other EN and IN subtypes (Figure 4G). As a specific example, we examined the expression of calmodulin (*CALM2*) and *GAPDH*, the two highest-expressed genes in M8 and M9. Both genes were found to be downregulated in L5 and L6 IT ENs and *LAMP5*^+^/*LHX6*^+^ INs (Figure 4H). This highlights agreement between our ST and snRNA-seq data, and posits specific cellular drivers for observed changes in layer-specific gene expression.

We conclude that while transcriptional alterations linked to increased age and executive dysfunction are similar, they are spatially distinct. Age-related changes are associated with neurons in shallower layers (L1-L5), whereas ALSci-linked changes are associated with neurons in deeper layers (L5-L6) of the cortex. Critically, executive dysfunction-linked transcriptional shifts are strongly associated with the BA46 region, unlike age-linked changes, which appear similar across both regions of the PFC. We note that, apart from age, no other confounding donor variables (e.g., ALS disease duration) was found to correlate with ALS or ALSci-linked spatial gene expression modules in either the ST data or in snRNA-seq-defined neuronal subtypes (Supplemental Figure S13).

### Executive Dysfunction is Associated with Mitochondrial and Synaptic Alterations

These findings implicate selective deep-layer neuronal subtype dysfunction in the pathophysiology of executive impairment in ALS. To address the nature of this dysfunction, we examined in detail gene expression changes in deep-layer neuronal subtypes linked to executive impairment (Figure 5A; see also Supplemental Figure S14). Across all five EN and IN subtypes, differentially expressed transcripts in BA46 of ALSci donors were enriched for three top gene ontology (GO) terms: presynapse (GO:0098793), postsynapse (GO:0098794), and mitochondrial membrane (GO:0031966). Within the synaptic category, pathways related to the synaptic vesicle cycle (GO:0099504) and synaptic organization (GO:0050808) were particularly prominent (Figure 5A). For BA46, but not BA44, we observed an ALSci-specific decrease in presynaptic gene expression, particularly vacuolar type H^+^-ATPase (V-ATPase) genes involved in synaptic vesicle acidification, alongside an increase in transcripts encoding for postsynaptic glutamate receptor subunits, such as *GRIK1*, *GRIA1*, and *GRM1* (Figure 5A). We also detected a strong downregulation of oxidative phosphorylation pathways and alterations in PINK1-Parkin-mediated mitophagy (Figure 5A). Interestingly, we found that oxidative phosphorylation and synaptic-related transcripts, including glutamatergic and GABAergic receptors, correlated within individual neuronal subtypes across donor tissue samples (Figure 5B), indicating that these changes co-occur in the same tissue and were not artifacts of separate donors having non-overlapping oxidative phosphorylation and synaptic responses.

**Figure 5:**
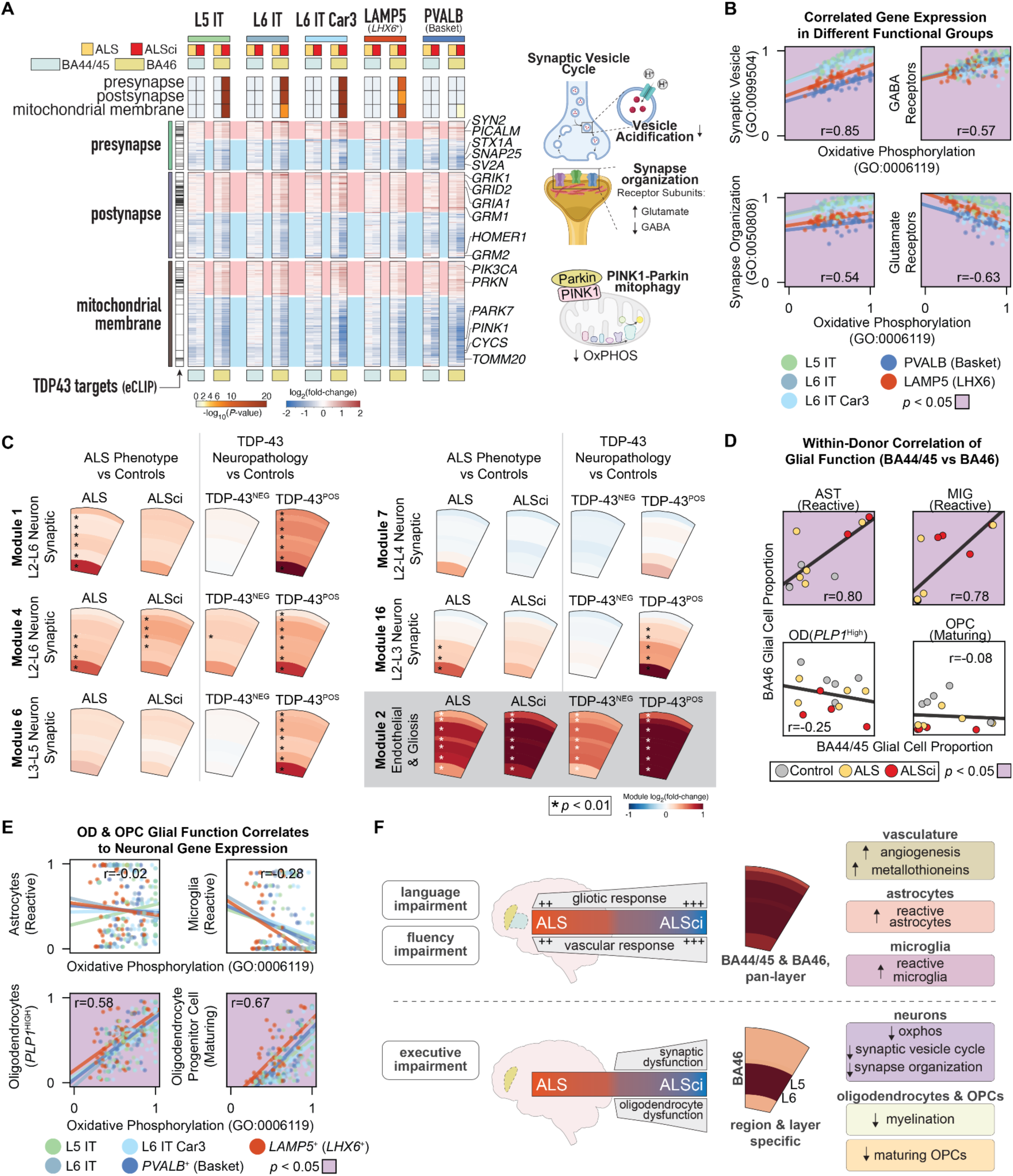
Executive Dysfunction is Associated with Mitochondrial and Synaptic Alterations. **A)** snRNA-seq differentially expressed genes (adjusted *p* < 0.05 from DeSeq2 on pseudobulk) in neuronal subtypes linked to executive impairment. Gene ontology (GO) enrichment of differentially expressed genes is plotted as -log_10_(*p*-value) for the top three GO functional terms, where white boxes are non-significant. Gene expression for transcripts that are both differentially expressed and annotated within these GO terms is plotted as log_2_ fold change (ALS vs. control and ALSci vs. control). Transcripts identified as TDP-43 binding by eCLIP (Enhanced Crosslinking Immunoprecipitation) are marked with black bars on the left side of the plot. **B)** Genes from selected GO terms (oxidative phosphorylation, synaptic vesicle, synaptic organization, glutamatergic receptors, and GABAergic receptors) were aggregated into ontology expression scores from snRNA-seq data using the same method employed to derive module scores. Mean ontology scores from neuronal cells for each donor sample are plotted. Scatter plots show oxidative phosphorylation scores on the x-axis and another ontology score on the y-axis, with best-fit linear regression line plotted for each neuronal subtype. Pearson correlation coefficients (*r*), averaged across the five neuronal subtypes, are also reported. Plots with correlations significantly different from zero (Wald test, *p* < 0.05) are shaded in lilac. **C)** Differential module expression from spatial expression data, stratifying ALS donors by cognitive or TDP-43 pathology profile. Asterisks indicate cortical layers with significant differences from control, tested by Benjaminini-Yekutieli corrected Welch’s *t*-test (*p* < 0.01). **D)** Cross-region, donor-level comparison of the proportion of glial cells annotated with specific functional subtypes between BA44/45 and BA46 (*n* = 14 matched samples). Reactive astrocytes and microglia plots with correlations significantly different from zero (Wald test, *p* < 0.05) are shaded in lilac. **E)** The proportion of reactive microglia, reactive astrocytes, high-myelinating oligodendrocytes, and maturing OPCs in each donor sample is plotted against oxidative phosphorylation ontology expression scores, calculated as in Figure 5B. Plots with correlations significantly different from zero (Wald test, *p* < 0.05) are shaded in lilac. **F)** Summary schematic highlighting region- and cortex layer-specific molecular differences between language impairment, verbal fluency impairment, and executive impairment.

TDP-43 aggregation is a pathological hallmark of ALS and has been implicated in disease-driving gene expression changes^42^, including prominently at the level of the synapse^43^. TDP-43 also accumulates in neuronal mitochondria and its dysfunction is linked to deficits in this organelle^44^, raising the possibility that TDP-43 pathology contributes to the transcriptional alterations we observed in these neuronal subtypes. To investigate this, we intersected differentially expressed genes from our dataset with TDP-43-associated targets, such as mRNAs identified by eCLIP-seq^45^, known protein-protein interactors^46^, and genes affected by TDP-43 nuclear depletion in postmortem FTD-ALS brains^47^ (Figure 5A, Supplemental Figure S15A). This analysis revealed a significant representation of eCLIP-defined TDP-43 target genes in our BA46 pool of ALSci-associated differentially expressed genes. Notably, TDP-43 eCLIP targets were predominantly upregulated (Figure 5A, Supplemental Figures S15A-C), and GO enrichment analysis pointed to the postsynapse as the most significantly affected cellular compartment (Supplemental Figures S15D,E). In contrast, protein-protein interaction partners and genes affected by nuclear TDP-43 depletion in FTD-ALS brains were not significantly enriched in our dataset (Supplemental Figure S15B).

### TDP-43 Pathology and Cognitive Impairment Have Partially Distinct Molecular Signatures

These observations raise the question of the extent to which TDP-43 pathology contributes to the transcriptional responses observed in the ALS prefrontal cortex. In our initial analyses, several neuronal spatial modules associated with neuronal function were upregulated in both ALS and ALSci donors (Supplemental Figure 10D), which we initially interpreted as reflecting gene expression changes shared across all ALS patients, irrespective of cognitive phenotype. However, when we re-stratified ALS donors by TDP-43 pathology—grouping them into those with and without neuropathological evidence of TDP-43 aggregation (TDP-43^POS^ and TDP-43^NEG^, respectively)—rather than by cognitive status, we found that many of the gene expression changes that occurred in both ALS and ALSci donors were, in fact, linked to the presence of TDP-43 pathology (Figure 5C and Supplemental Figure S16A). Modules enriched for neuronal gene expression, particularly those linked to synaptic function (e.g., modules M6), showed minimal change in TDP-43^NEG^ donors in relation to controls but were significantly upregulated in TDP-43^POS^ individuals (Figure 5C and Supplemental Figure S16A). This contrast was not apparent when stratifying by cognitive phenotype, suggesting that, at least in part, synaptic transcriptional alterations in ALS are mechanistically tied to TDP-43 aggregation (Figure 5C and Supplemental Figure S16A), consistent with the strong overlap between eCLIP-defined TDP-43 targets with synapse-related genes (Figure 5A and Supplemental Figures S15C,D). Spatial modules not enriched for synaptic genes, such as the gliosis-associated M2 module, were significantly upregulated in both TDP-43^NEG^ and TDP-43^POS^ donors (Figure 5C and Supplemental Figure S16A). In line with this, the proportion of reactive glia was also similar between TDP-43^NEG^ and TDP-43^POS^ cases (Supplemental Figure S16B), indicating that this gliotic response is likely driven by alternative mechanisms (Supplemental Figure S16B). Together, these data highlight a nuanced relationship between TDP-43 dysfunction and cognitive deficits: pathology and clinical phenotype do not fully overlap at the molecular level but instead reveal distinct features of ALS pathophysiology. This is in line with previous findings that TDP-43 aggregation alone correlates poorly with cognitive symptom manifestation^48^.

### Neuron-glia correlates of cognitive impairment

Our analyses so far uncovered both shared and region-specific patterns of neuronal and glial dysregulation, which, when considered together, may provide a better understanding of the cellular mechanisms underlying cognitive impairment in ALS. To this end, we examined within-donor correlations across BA44/45 and BA46 to assess whether these patterns reflected coordinated or region-specific changes. These analyses revealed a shared pattern of gliosis across BA44/45 and BA46 primarily driven by astrocytes and microglia (Figure 5D and Supplemental Figure S17A). In other words, when this gliosis signature was present in BA44/45, it was also consistently observed in BA46 (Figure 5D), suggesting that certain glial activation patterns may be broadly expressed within the frontal cortex in ALS, potentially reflecting a common gliotic response to ALS irrespective of cognitive phenotype. While astrocytic and microglial responses unfolded similarly between BA44/45 and BA46, myelinating oligodendrocytes (*PLP1*^High^) and maturing OPCs displayed region-specific behaviors, as revealed by the weak correlation in their proportions across these regions (Figure 5D). Interestingly, the abundance of both glial subtypes tracked closely with neuronal transcriptional dysregulation, particularly in relation to synapse- and mitochondria-associated genes (Figure 5E, Supplemental Figures S17B-D). Overall, this oligodendrocyte/maturing OPC/neuron correlation emerged as one the strongest within-sample associations, and was largely independent of neuronal subtype (Supplemental Figure S17B,D). While the basis for this regional heterogeneity remains unclear, these results uncover a complex landscape of neuron-glia interactions that correlate with cognitive impairment in ALS (Figure 5F).

## Discussion

ALS is traditionally framed as a disease of motor neurons^1^. Together with other recent reports^9,11,12^, our study provides a broader understanding of this neurological disorder beyond its motor manifestations and contributes to a more unified understanding of this devastating disease. By combining detailed patient stratification with transcriptome-wide profiling of two prefrontal cortex regions, we identify spatially defined signatures associated with cognitive changes and trace molecular patterns linked to language, verbal fluency, and executive impairment. These changes occur in the absence of overt neuronal degeneration, suggesting that mild cognitive impairment in ALS precedes neuronal loss in the prefrontal cortex. Instead, our findings support a model of the disease in which abnormal interactions between neurons, glia, and the vasculature, rather than the simple degeneration of vulnerable neurons, are mechanistically involved in the decline of cognitive function. These interactions differ between clinical phenotypes, indicating a heterogeneous disease etiology driven by distinct mechanisms across affected regions rather than a uniform pathological process.

We identified two distinct glial signatures in the ALS prefrontal cortex. One is a classic gliosis response^49^, shared between BA44/45 and BA46, that involves a coordinated increase in reactive microglia, reactive astrocytes, and stressed OPCs across all cortical layers and white matter. While upregulated in ALS donors, this gliotic response is largely not correlated to cognitive impairment. The other is more regionally restricted, being characterized by a decrease in high-myelinating oligodendrocytes alongside an increase in maturing OPCs, and lacks correspondence between the two regions. While the glial alterations we detected were generally more pronounced in ALSci, they were broadly shared across the ALS spectrum, suggesting that additional mechanisms contribute to cognitive decline in ALS. Indeed, the widespread gliotic signature we characterized, marked by increased reactive microglia and astrocytes and observed across both prefrontal cortex regions, could reflect hypoxic stress due to respiratory failure, commonly seen at terminal stages of ALS^50^. In contrast, the regional specificity of the oligodendrocyte/OPCs response—and its transcriptional coupling to neuronal gene programs in BA46—points to a possible link to the emergence of executive deficits.

Deficits in executive function appear to stem from specific neuronal subtypes localized to the deeper layers of BA46. At the molecular level, these neurons show robust alterations in synaptic processes—both pre- and postsynaptic—as well as in mitochondrial function. In contrast, impairments in language and verbal fluency are associated with more diffuse transcriptional patterns, largely involving glial and vascular processes. Specifically, language deficits correlate with molecular signatures shared across both sampled brain regions, whereas verbal fluency deficits are linked specifically to BA44/45 but involve broad transcriptional responses spanning many of the spatial co-expression modules we identified. These programs may reflect the complexity of the brain’s language network, which is distributed across interconnected frontal and temporal regions^18^. It is plausible, therefore, that the diffuse signals we observed stem from this anatomical complexity—a possibility that future studies could test by expanding sampling to additional brain regions involved in language processing.

TDP-43 proteinopathy is a pathognomonic feature of the broader ALS clinical spectrum^42^. This RNA-binding protein is ubiquitously expressed in the brain and typically localizes to the nucleus. However, in disease states, TDP-43 is redistributed to the cytoplasm where it forms insoluble aggregates. The resulting loss of nuclear TDP-43 is linked to perturbations in gene expression, which are hypothesized to contribute to ALS pathogenesis^42^. Our analyses revealed that the relationship between TDP-43 neuropathology and clinical outcomes is nuanced. As previously reported^48^, TDP-43 aggregation was not predictive of cognitive deficits in our cohort; however, some molecular responses, particularly those linked to neuronal and synaptic function, were tightly associated with TDP-43 pathology. This suggests that TDP-43 dysfunction is not in itself sufficient to drive the development of cognitive impairment in ALS, which is dependent on additional disease mechanisms such as the coordinated responses we observed involving neurons, glia, and vasculature. In other words, the cognitive dimension of ALS cannot be understood as a linear outcome of TDP-43 pathology. Instead, cognitive phenotypes in ALS appear to have a multicellular etiology, with regional specificities, that interact and are likely modulated by TDP-43 dysfunction.

In summary, we have uncovered complex relationships between clinical phenotype and cellular dysfunction in ALS. Our findings highlight cognitive domain-specific transcriptional signatures that are not only anatomically distinct but also exhibit layer-specific localization within the prefrontal cortex. This suggests that different cognitive impairments in ALS arise from disruptions in unique neuronal and glial populations, underscoring the need for targeted therapeutic strategies.

## METHODS

### Tissue sourcing and donor information

Postmortem tissue specimens were obtained from the Medical Research Council (MRC) Edinburgh Brain Bank (Research Ethics Committee approval: 21/ES/0087), following review by the bank’s internal ethics committee, the Academic and Clinical Central Office for Research and Development Medical Research Ethics Committee, and the Biomedical Research Alliance of New York. Tissue specimens originated from patients recruited through the Scottish Motor Neurone Disease Register (a prospective, population-based ALS study in Scotland) who had consented to the collection and use of their health-related information for research purposes. Clinical metadata associated with each donor, including ECAS test scores, were extracted from the Clinical Audit Research Evaluation (CARE)-MND database. Tissue from neurotypical control cases – defined in general terms as individuals without neurological or psychiatric conditions and no substantial neuropathology postmortem – was provided by the Edinburgh Sudden Death Brain Bank ^21^. All applicable federal, state, and local regulations were followed when transferring materials to the New York Genome Center.

The diagnosis of patients fulfilled the revised El Escorial criteria for clinically definite or probable ALS. Comorbid neurological disorders, including frontotemporal dementia, or psychiatric history were exclusion criteria for the selection of study participants. We note also that dementia was not documented in the clinical files of any recruited patients. Clinical metadata is provided in Supplemental Figure 1 and Supplemental Table 1.

### Cognitive profiling

Cognitive performance was assessed in life using the Edinburgh Cognitive and Behavioural ALS Screen (ECAS) following standard guidelines (https://ecas.psy.ed.ac.uk). Briefly, the ECAS incorporates 16 subtests assessing a range of cognitive domains, including executive functions (reverse digit span, alternation, inhibitory sentence completion, and social cognition), verbal fluency (free and fixed), language (naming, comprehension, and spelling), memory (immediate recall, delayed percentage retention, and delayed recognition), and visuospatial functions (dot counting, cube counting, and number location), complemented by a separate semi-structured interview with a relative or carer designed to detect behavioral changes^19,51^. Performance across all tasks generates the ECAS Total Score, which adds up to an aggregated maximum of 136 points. The ECAS Total Score is composed of two fractions, the ALS-Specific and ALS Non-Specific Scores. Subtest evaluations of executive functions, language, and verbal fluency are combined to generate the ALS-Specific Score, which can reach a maximum of 100 points in cognitively healthy individuals. Memory and visuospatial functions tasks contribute to the ALS Non-Specific Score. Abnormal scores are defined as ≤77 for the ALS-Specific Score, ≤24 for the ALS Non-Specific Score (out of a maximum of 36 points), and ≤105 for the ECAS Total Score. Further subdomain-specific thresholds include ≤33 out of 48 for executive function, ≤14 out of24 for fluency, ≤26 out of 28 for language, ≤13 out of 24 for memory, and ≤10 out of 12 for visuospatial functions.

### Neuropathology Assessments

Brain regions of interest, including BA44/45 and BA46, were fixed in 10% neutral-buffered formalin for a minimum of 24 hours to ensure tissue integrity and morphology. Post-fixation, tissue processing involved a dehydration step through an ascending ethanol series (70%–100%), followed by clearing with three rounds of xylene treatment. Specimens were then embedded in molten paraffin wax over three consecutive 5-hour stages and, after solidifying at room temperature, sectioned onto Superfrost Plus slides on a Leica microtome at 4 μm thickness with overnight drying at 40°C. All staining procedures were performed using a Leica Bond RX autostainer instrument. Aggregated TDP-43 was detected using an antibody against phosphorylated TDP-43 (2B Scientific, #CAC-TIP-PTD-MO1, 1:2000; Leica Bond RX: protocol F ER1 — 30 mins) , combined with 3,3′-diamino-benzidine (DAB) visualization. TDP-43 pathology was staged using a semiquantitative four-tiered scoring system, where a score of 0 indicates the absence of pathology, and scores 1 through 3 represent progressively increasing levels of TDP-43 pathology burden (mild, moderate, and severe, respectively). Scores were defined as follows: 1 (mild) for up to 5 affected cells, 2 (moderate) for 6–15 affected cells, and 3 (severe) for more than 15 affected cells, all within at least a single 20× field of view per tissue section. This scoring scheme was applied independently to both neurons and glia using haematoxylin counterstaining for morphology-based cell classification ^5^. Tau: Thermo Fisher, #MN1020, 1:1000; Leica Bond RX: protocol F, no pretreatment. Beta-amyloid: BioLegend, #800712, 1:16000; Leica Bond RX: protocol F wet start, formic acid pretreatment (5 mins). Alpha-synuclein: Leica, #NCL-L-ASYN, 1:20; Leica Bond RX: protocol F ER2 (30 mins).

### Nuclei isolation from frozen post-mortem brain tissue and library preparation

Nuclei isolation protocol was adapted from previously described methods ^52,53^. All procedures were carried out on ice unless otherwise stated. Frozen post-mortem brain tissue was retrieved from storage at -80 °C and equilibrated on dry ice for 20–30 minutes before being transferred to a cryostat at -18 °C. Tissue samples were left for 15–20 minutes inside the cryostat before sectioning. We collected 20-50 mg of tissue per preparation, taking care to avoid excessive sampling of white matter. Tissue cuts were placed in a pre-chilled Dounce homogenizer containing 700 µL of ice-cold homogenization buffer (320 mM sucrose, 5 mM CaCl_2_ [Millipore Sigma; #21115;], 3 mM magnesium acetate [Millipore Sigma; #63052], 10 mM Tris HCl [pH 7.8; prepared from Thermo Fisher’s #AM9850G and #AM9855G], 0.1 mM EDTA [pH 8.0; Thermo Fisher, #AM9260G], 0.1% IGEPAL CA-630 [Millipore Sigma; #I8896], 1 mM DTT [Millipore Sigma; #646563], and 0.4 U/µl Protector RNase inhibitor [ Millipore Sigma; #03335402001]). Homogenization was performed with 10 strokes of the loose pestle followed by 10 strokes of the tight pestle. After a 5-minute incubation on ice, the homogenized tissue was passed through a 70-µm Flowmi Cell Strainer (Bel-Art; #H13680-0070) and combined with an equal volume of a solution made of 50% OptiPrep (#07820, STEMCELL Technologies), 5 mM CaCl_2_, 3 mM magnesium acetate, 10 mM Tris HCl (pH 7.8), 0.1 mM EDTA (pH 8.0), and 1 mM DTT. Using a 15-mL conical tube, this mixture was then layered onto a density gradient made up of 750 µL of a 30% OptiPrep solution supplemented with 134 mM sucrose, 5 mM CaCl_2_, 3 mM magnesium acetate, 10 mM Tris HCl (pH 7.8), 0.1 mM EDTA (pH 8.0), 1 mM DTT, 0.04% IGEPAL CA-630, and 0.17 U µl^−1^ RNase inhibitor) over 300 µL of 40% OptiPrep solution supplemented with 96 mM sucrose, 5 mM CaCl_2_, 3 mM magnesium acetate, 10 mM Tris HCl (pH 7.8), 0.1 mM EDTA (pH 8.0), 1 mM DTT, 0.03% IGEPAL CA-630, and 0.12 U µl−1 RNase inhibitor. Nuclei were separated by centrifugation at 3,000 × *g* for 20 minutes at 4 °C using a pre-chilled swinging-bucket rotor with the brake off. Approximately 150 µL of nuclei were collected from the 30%/40% gradient interphase and subsequently washed in PBS containing 1% BSA and 1 U µl^−1^ RNase inhibitor. Following a 5-minute incubation on ice, the nuclei were centrifuged at 450 × *g* for 3 minutes at 4 °C, and the pellet resuspended in 15 µL of diluted 10X Genomics Nuclei Buffer. Nuclei counting was performed using a Countess II FL instrument (Thermo Fisher, #AMQAF1000) by staining with SYTOX Green (Thermo Fisher; #S7020). Additional quality control measures included Trypan Blue staining (#EBT-001, NanoEntek) to evaluate adequate permeabilization (cutoff: >95% of ‘dead cells’) and the assessment of nuclear morphology by light microscopy before proceeding with transposition.

Matched snRNA-seq and snATAC-seq libraries were prepared using the Chromium Next GEM Single Cell Multiome ATAC + Gene Expression Kit (10X Genomics, #PN-1000283) according to the protocol provided by 10X Genomics. Nuclei (16,100 per reaction; targeted recovery: 10,000 nuclei) from a single donor were used for each library preparation. Partitioning of nuclei was performed on a Chromium X instrument (10X Genomics, #1000331). Quality control of cDNAs and libraries was conducted using Agilent High Sensitivity DNA Kits and a BioAnalyzer instrument. Final library quantification prior to pooling utilized KAPA Biosystems’ Library Quantification Kit for Illumina Platforms (KK4824). Libraries were sequenced on a NovaSeq X system using a 2 × 100 bp configuration run on a 10B flow cell. BA44/45 and BA46 libraries (*n* = 48) were prepared and sequenced in independent batches.

### Prefrontal Cortex Atlas and Classifier

Published 10X Genomics snRNA-seq sequencing data was acquired from NCBI GEO or Synapse repositories: GSE157827 ^24^, GSE167494 ^25^, GSE174332 ^9^, GSE174367 ^26^, GSE207334 ^27^, GSE219280 ^12^, PRJNA434002 ^28^, syn45351388 ^23^, and syn51123517 ^29^ (Supplemental Table 2).

Sequencing data was aligned to a standard reference genome (hg38; GENCODE v32/Ensembl98) and counted (cellranger v7.1.0). Individual cells were filtered so that the number of transcripts (UMIs) was between 500 and 50000, and the maximum fraction of mitochondrially-encoded transcripts was 0.2. Prefrontal cortex atlas was restricted to donor samples from patients between 50 and 90 years of age at death taken from the prefrontal cortex. The resulting dataset consists of 2,603,756 cells from 407 donors.

Each cell was scaled to a total of 10,000 counts, the count data was transformed by log(x + 1), and each gene was scaled by dividing by the interquartile range (RobustScaler). Genes which would have a variance greater than 1 after scaling were further scaled so that their variance was equal to 1. Cells were embedded into a k-nearest neighbors (k-NN) graph from 50 principal components and 25 neighbors, then clustered into 114 separate clusters using the leiden algorithm (resolution set to 2.5). Labels for cell type and cell subtype were hand annotated for each cluster based on known marker genes (Supplemental Table 3).

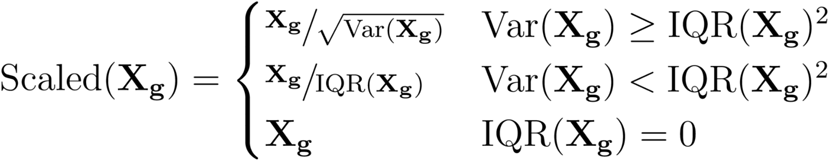

This standardized, transformed, scaled data was used as input to a neural network (NN) autoencoder/classifier that predicts cell type, subtype, and denoised transcript expression data. The NN encoder module takes 36601 transcript counts as input, and is fully connected to a 1000 node hidden layer, a 250 node hidden layer, then a 100 node hidden layer, each of which has a hyperbolic tangent (tanh) activation function. The 100 node hidden layer is connected to a 50 node (h_width_) hidden layer with a tanh activation function that represents a shared latent space embedding. This hidden layer is shared between three decoder modules. The transcript expression predictor connects the 50 node shared latent space embedding to a 50 and then 250 node hidden layers with a tanh activation function, then a 36601 node output layer with a rectified linear (ReLU) activation function. The cell type classifier connects the 50 node shared latent space embedding to a 50 node hidden layer with a tanh activation function, and connects that hidden layer to a 9 node output softmax layer that predicts cell type label. The cell subtype classifier connects the 50 node shared latent space embedding to a 50 node hidden layer with a tanh activation function, and connects that hidden layer to a 14 node output softmax layer that predicts cell subtype label.

The NN model was constructed in pyTorch and trained with the Adam optimizer. Model hyperparameters h_width_, weight decay, and learning rate were tuned by training on randomly sampled 465,000 cell training sets for 200 epochs and scoring on 155,000 cell validation sets. Transcript expression was trained to minimize mean squared error, and cell type and subtype classifiers were trained to minimize cross-entropy. Full training was performed using 1.95m cells for 1500 epochs, and scored on 650k cells held out. Held out test cells were not used during cross-validation. Models were evaluated based on coefficient of determination (R^2^) for transcript expression, and on F1 score for cell type and subtype classifiers.

### 10X Genomics Multiome Data Processing

Sequenced 10X Genomics gene expression and ATAC libraries were aligned to a standard reference genome (hg38; GENCODE v32/Ensembl98) and counted (Cellranger-ARC v2.0.2; Supplemental Table 4). ATAC peaks were aligned between samples for quantification (Cellranger-ARC aggr v2.0.2). Individual cells were filtered so that the number of gene expression transcripts (UMIs) was between 500 and 50000, the maximum fraction of mitochondrially-encoded transcripts was 0.2, and the number of quantified ATAC fragments in peaks was between 250 and 100,000. A total of 165,884 cells passed this initial filtering. Cell types and subtypes were determined by the prefrontal cortex cell type classifier, and validated by comparing to published marker genes (Supplemental Table 3). Doublets and low-quality cells were removed, leaving 144,260 usable cells from 48 libraries for analysis.

To maintain the difference in count depth between cell types, individual cells were standardized for count depth by dividing all transcript counts for the cell by a size factor, to reach the median count depth for the assigned cell type. Standardized counts were log(x+1) transformed, and each gene *g* was then scaled by dividing by the interquartile range (IQR). Genes which would have a variance (Var) greater than 1 after scaling were further scaled so that their variance was equal to 1, giving log standardized counts.

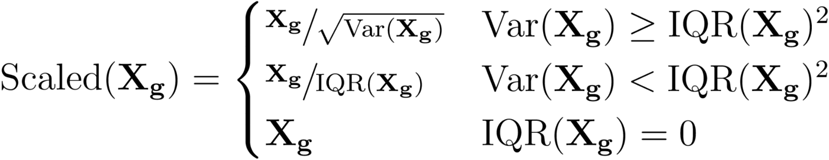

Principal Component Analysis (PCA) was performed using a truncated singular value decomposition with 25 components on the log standardized counts. Cells were embedded into a 25-neighbor k-Nearest Neighbors (k-NN) graph using these 25 components. Cells were projected into 2 dimensions with Uniform Manifold Approximation and Projection (UMAP) using the k-NN embedding (Minimum distance of 0.5; initialized from spectral graph embedding).

For ATAC analysis, the 144,260 usable cells were further filtered for cells with at least 2500 ATAC peak counts, yielding 119,153 cells with both RNA and ATAC modalities. Individual cells were standardized for ATAC count depth by dividing all peak counts for the cell by a size factor, to reach the median ATAC count depth for the assigned cell type. Standardized ATAC counts were then log(x+1) transformed to give log standardized ATAC counts. PCA, k-NN embedding, and UMAP projection were performed identically to gene expression data from log standardized ATAC counts.

### Glial Cell Functional Annotation

snRNA-seq log standardized counts for each glial cell type were separated for PCA (25 components), k-NN graph embedding (25 neighbors), and UMAP as for the full dataset. Cells from each cell type were separately clustered into a large number of groups with the leiden algorithm (resolution of 2), and then each cluster was assigned to one of a few functional groups based on relative expression of key functional marker genes (Supplemental Table 6).

The proportion of glial cells within each functional category was calculated for each snRNA-seq donor sample by taking the number of functionally annotated glial cells and dividing it by the total number of glial cells of that type (e.g. the number of homeostatic astrocytes divided by the total number of astrocytes).

### Neuronal Cell Subtype Annotation

snRNA-seq log standardized counts for excitatory (EN) and inhibitory neurons (IN) were further filtered by transcript count to neurons with a minimum of 2500 UMI counts, yielding 41,933 ENs and 15,553 INs. ENs and INs were separately processed for PCA (25 components), k-NN graph embedding (25 neighbors), and UMAP as for the full dataset. ENs and INs were then clustered into transcriptionally distinct clusters with the leiden algorithm (resolution of 3), and clusters were assigned to a neuronal subtype based on marker expression (Supplemental Table 6).

### Cell Proportions Per Sample

For each single nucleus sample (n=48), the number of cells in each functional group for a glial cell type are divided by the total number of cells of that type (e.g., 60 homeostatic astrocytes and 40 reactive astrocytes have proportions of 0.6 and 0.4). The correlation between these ratios is calculated as the within-sample correlation of glial cell function proportions. Welch’s *t*-test was used as the statistical test between proportions in different groups. Wald test was used as the statistical test for correlations. Multiple comparisons were corrected with Benjamini-Hochberg as appropriate.

### Single-cell Differential Expression Analysis

Transcript differential expression analysis was performed on pseudobulked raw integer counts with pyDeseq2 ^54,55^, standardizing using only non-zero counts (‘poscounts’) and excluding mitochondrially encoded genes and *MALAT1* from size factor calculations. Statistical testing was against a null hypothesis of 0, rejecting the null for adjusted *p*-values ≤ 0.05.

### Gene Ontology Enrichment Analysis

Gene ontology enrichment was performed with Metascape using the default complete proteome as the background of each analysis ^56^, and g:Profiler ^57^, which tests gene lists for enrichment against background a multiple hypothesis test corrected hypergeometric test. Here, gene lists are differentially expressed genes against a background of genes which are not all-zero in counts, or genes in a module against a background of genes which were included in the Splotch spatial model.

### Tissue processing, RNA Capture, and Library Preparation for Spatial Transcriptomics

Frozen tissue blocks, kept at -80°C prior to processing, were equilibrated to -16°C inside a Leica CM3050S cryostat, embedded in Tissue Plus OCT Compound (Fisher Healthcare, #4585), and sectioned in a perpendicular orientation with regard to the pial surface. Tissue sections (10 µm-thick) from quality controlled blocks (H&E staining and imaging) were collected onto prechilled Visium Spatial Gene Expression Slides (10X Genomics, #1000185) while briefly warming the back of the slide with a gloved finger to improve adherence and minimize artifacts.

We sought to capture cortical layers 1 through 6 and white matter within each spatially barcoded array. For each subject, four technical replicates, sampled from tissue cuts approximately 20 µm apart, were collected across two Visium slides. A deliberate effort was made to section control and ALS subjects onto the same Visium slide when possible, with sample processing proceeding in parallel from that point onward. Where necessary due to tissue sectioning artifacts or poor yield, additional replicates were collected and processed on a later date.

Generation of spatially resolved transcriptomes was performed in accordance with 10X Genomics guidelines (CG000239, Revisions F and G). Briefly, tissue sections were fixed in chilled methanol, stained using an abbreviated H&E protocol, and imaged on a Zeiss Axio Observer Z1 fitted with an EC Plan-Neofluar 10x/0.3 M27 objective and a Zeiss Axiocam 506 Mono camera. Tissue sections were permeabilized for 12 minutes following optimization (10X Genomics, CG000238, Revision E). The number of cDNA amplification cycles for each Visium array was calculated using qPCR, as instructed by 10X Genomics.

Visium libraries were prepared using unique Illumina-compatible PCR primers with Dual Index Kit TT Set A (10X Genomics, #1000215). Size distribution profiles of the final libraries were assessed by running Agilent High Sensitivity DNA chips (#5067-4626) on a 2100 Bioanalyzer system (Agilent, #G2939BA). Libraries outside the expected range (450 ± 50 bp) or that contained adapter dimer contaminants were discarded and reprocessed. Total yields were initially measured on a Qubit 3.0 fluorometer before final qPCR quantification (KAPA Library Quantification Kit for Illumina Platforms, KAPA Biosystems, #KK4824). Per batch, libraries from 48-52 Visium arrays were pooled at a concentration of 10 nM prior to sequencing. We processed 246 arrays from 46 unique donors.

### 10X Genomics Visium Data Processing and Annotation

Each sequenced 10X Visium library was aligned to a standard reference genome (hg38; GENCODE v32/Ensembl98) and counted with spaceranger (v3.0.0; Supplemental Table 5). We initially filtered any spot with fewer than 100 UMI transcript counts. Visium spots were manually assigned to one of the six cortical layers or the white matter using 10X Genomics Loupe Browser (version 8.0). Layer annotations were based on cortical architecture and cellular morphology. The white matter was identified by the absence of neuronal cell bodies and the linear arrangement of glial nuclei along neuronal processes. Cortical layer 6 was defined as a heterogeneous neuronal layer spanning from the white matter to the beginning of cortical layer 5, characterized by its large pyramidal neurons oriented perpendicularly to the pial surface. Cortical layer 4 was identified by the presence of small, granular neurons adjacent to the large pyramidal neurons of cortical layer 5. Cortical layer 3 was labeled as the pyramidal cell layer located between granular cortical layers 2 and 4. Cortical layer 1 was distinguished by its paucity of cells and lack of large neuronal somas. Spots containing meninges contiguous to cortical layer 1 were excluded from downstream analysis. Additionally, spots overlying tissue artifacts, such as freeze fractures, folds, or tears, were excluded. After annotation, we have 953,073 individual spots across 246 arrays.

To maintain the difference in count depth between cell types, individual spots were standardized for count depth by dividing all transcript counts for the cell by a size factor, to reach the median count depth for the annotated cortex layer. Size factors were clipped to a minimum of 0.25 to mitigate the risk of overinflating low-count spots. Standardized spatial counts were then scaled by dividing by the interquartile range (RobustScaler). Genes which would have a variance greater than 1 after scaling were further scaled so that their variance was equal to 1, giving standardized spatial counts.

PCA was performed using a truncated singular value decomposition with 25 components on the standardized counts. Cells were embedded into a 25-neighbor k-NN graph using these 25 components. Cells were projected into 2 dimensions with UMAP using the k-NN embedding (Minimum distance of 0.5; initialized from spectral graph embedding).

### Splotch modeling of spatial gene expression across clinical groups

Raw sequencing data was filtered to remove mitochondrially encoded genes, lncRNAs, pseudogenes, and genes expressed in less than 1% of spots (this filtering was applied separately to BA44/45 and BA46 datasets). Spots with fewer than 100 UMIs across remaining genes were discarded prior to Splotch analysis. Finally, spots without a valid annotation were discarded, and any spots without at least one immediate neighbor in the Visium grid were discarded to prevent singularities in the spatial autocorrelation component of the Splotch model. This yielded expression data for 13,188 genes measured across 401,976 spots (286,581 ALS, 115,386 control) with an average depth of 3,679 UMIs in BA46, and 13,116 genes measured across 551,106 spots (347,548 ALS, 203,558 control) with an average depth of 3,447 UMIs in BA44.

Splotch analysis was conducted as described previously^13,36,37^. Briefly, Splotch employs a zero-inflated Poisson likelihood function (where the zero-inflation accounts for technical dropouts) to model the expression (λ) of each gene in each spot. This expression level is in turn modeled using a generalized linear model (GLM) with the following three components: the characteristic expression of the spot’s donor ID and ALS phenotype (β), autocorrelation with the spot’s spatial neighbors (ψ), and spot-specific variation (ε). Furthermore, the characteristic expression rate (β) is hierarchically formulated to account for the experimental design, enabling the investigation of the effect of sample covariates on differential gene expression. By performing posterior inference on the model given the observed spatial expression data, we identified clinical group level trends in spatial gene expression (β) that best explain our observations.

Compared to other computational methods for analysis of spatial transcriptomics data, Splotch i) enables quantification of expression differences between conditions and anatomical regions, ii) is designed to rigorously account for the uncertainty of low counts, and iii) analyzes multiple tissue sections simultaneously, stratifying patients based on clinical phenotype in order to quantify biological and technical variation.

### Parameter inference

Splotch has been implemented in the probabilistic programming language Stan, and is available at https://github.com/adaly/cSplotch. For all analyses, Bayesian inference was performed over the parameters using Stan’s adaptive Hamiltonian Monte-Carlo (HMC) sampler with default parameters. Four independent chains were sampled, each with 175 warm-up iterations and 175 sampling iterations (700 total), and convergence was monitored using the R-hat statistic.

### Splotch differential expression analysis

To quantify difference in expression between two conditions using Splotch, the Bayes factor between posterior distributions over characteristic expression coefficients β estimated by the model was calculated. Without loss of generality, difference in expression was quantified between conditions represented by *β*(1) and β(2) , which may differ across any combination of genes, donor covariates, or cortical layers. A random variable Δ*_β_*= *β*(1) ⎻ *β*(2) was defined, which captures the difference between *β*(1) and Δ*_β_*. If Δ*_β_* is tightly centered around zero, then the two distributions are very similar to each other, and the null hypothesis of identical expression cannot be rejected. To quantify this similarity, the posterior distribution Δ*_β_* | *D* (where *D* represents the data used to train the model) was compared to the prior distribution Δ*_β_* using the Savage-Dickey density ratio^58^ that estimates the Bayes factor between the conditions:

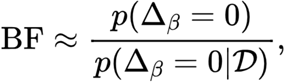

where the probability density functions were evaluated at zero. If expression is different between the two conditions, then the posterior Δ*_β_* | *D* will have very little mass at 0, and the estimated Bayes factor will be large (by convention, BF > 5 indicates substantial support). Conversely, for similar expression regimes, the posterior will place a mass equal to or greater than that of the prior at zero, and the Bayes factor will be ≤ 1.*p*(Δ*_β_* | *D*). is calculated using the posterior samples obtained in the previous section.

### Assigning Genes to Spatial Modules

Spatial modules are inferred from the mean posterior estimates (λ) of the depth-normalized expression of each gene in each spatial spot (a matrix of dimension 13,188 genes x 401,976 spots in BA46; 13,116 genes x 551,106 spots in BA44/45). Mean posterior gene expression estimates λ for each gene were clipped to the 99th percentile to remove the effect of outlier spots. Gene expression values were then scaled by dividing by the gene interquartile range, and genes which would have a variance greater than 1 after scaling were further scaled so that their variance was equal to 1. Pearson correlation coefficients were then calculated across all spots for all pairs of genes, resulting in a 13,188 x 13,188 correlation matrix in BA46 and 13,116 x 13,116 correlation matrix in BA44/45.

Correlation distance between 13,253 genes was calculated by taking the mean Pearson correlation distance (1 - pearson correlation coefficient) for a gene-gene pair from both brain regions. This correlation distance matrix was used as the distance matrix for a gene-gene k-NN graph embedding (k=10). Genes were projected into 2 dimensions with the UMAP algorithm, and clustered into 23 clusters using the leiden algorithm (resolution of 2). Each cluster is considered a spatially-correlated gene module. Clusters 21 and 22 contain all chrM and chrY-encoded transcripts, respectively.

Spatial modules and submodules are listed in Supplemental Table 7. Gene Ontology enrichment for each spatial module is listed in Supplemental Table 8. Python functions for module assignment are available in the scself (single-cell self-supervised; v0.4.8) package, installable through python’s pip package manager.

### Module Scores

For each gene, standardized transcript count (snRNA-seq) or modeled expression lambda (spatial) was standardized to a 0-1 range for each gene, where 0 is fixed to the 1st percentile, 1 is fixed to the 99th percentile, and values are linearly scaled between 0 and 1. Values below the 1st percentile are clipped to 0, and values above the 99th percentile are clipped to 1. For each module, a module score is calculated for each cell (snRNA-seq) or array spot (spatial) by taking the mean of the 0-1 scaled gene scores for every gene in the module.

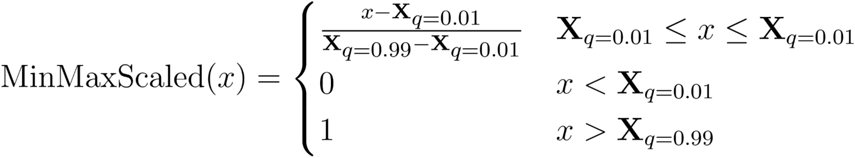

Spatial spot-level module scores are aggregated by taking the median module score for all spots in a layer (L1-L6 and WM) for each array, and rescaling the median layer-array module scores to a 0-1 range. These scaled layer-array module scores are used for statistical testing, and are further aggregated into mean module scores per ALS phenotype for plotting.

snRNA-seq module scores are calculated for each cell as above, and aggregated to the donor tissue sample by taking the mean module score for all cells of a specified type. These aggregated mean scores are rescaled to a 0-1 range and used for statistical testing, and are further aggregated into mean module scores per ALS phenotype for plotting.

Welch’s *t*-test was used for statistical tests between module scores in different groups, controlling for false discovery with Benjamini-Hochberg for snRNA-seq or Benjaminini-Yekutieli for ST data.

### Gene Ontology Expression Scores

Gene ontology expression scores were calculated as in Module Scores, using genes that are annotated with specified GO terms. GO term annotations used are listed in Supplemental Table 9.

### Assigning Genes to Cell Type Submodules

A gene-gene correlation matrix for all genes in a spatial module was constructed from standardized snRNA-seq transcript expression. This was subtracted from 1 to get a correlation distance matrix, which was used as the distance matrix for a gene-gene k-NN graph embedding (k=10). Genes were projected into 2 dimensions with the UMAP algorithm, and clustered into 2 to 5 clusters using the leiden algorithm (resolution of 0.75). Each cluster was interpreted as a gene submodule, and this process was repeated for all 23 spatial modules. Modules 21 and 22 (chrM and chrY) did not yield more than one submodule. Submodule scores were calculated for individual cells and spatial spots as above. Spatial modules and submodules are listed in Supplemental Table 6.

### Spatial Co-occurrence Analysis

Pearson correlation was calculated for spatial module and spatial submodule scores across all spatial spots. Modules and submodules are assigned to cortex layers by aggregating into layer-array module scores as above, and then taking the average module score across all layers. The module is assigned to the layer with the highest module score, and any other layer within 95% of that value. If all scores are within 10%, the module is assigned to the pan-layer label ‘ALL’.

### Spatial Cell Type Deconvolution

Per-spot estimates of cell type abundances were generated using Cell2location^30^. The prefrontal cortex atlas assembled in *Prefrontal Cortex Atlas and Classifier* was used to train a negative binomial (NB) regression model of gene expression for each of the 5 major glial cell types, for the 4 inhibitory neuron subtypes, and for the 3 excitatory neuron layer subtypes (Supplementary Table 3), which was used for all subsequent deconvolution experiments. Deconvolution with Cell2location was performed separately for each donor, treating individual ST arrays as batches and assuming an average of five cells per Visium spot with all other hyperparameters set to default values. Mean estimates of the abundance of each cell type in each spot were extracted and used for differential abundance testing.

### SST^+^ Cell Counting

SST labelling (Santa Cruz Biotechnology, #SC-74556, 1:200) was carried out on a Leica Bond RX autostainer (protocol F ER1 — epitope retrieval duration: 20 minutes) following standard IHC procedures. Raw H&E images were first analyzed to differentiate between white matter and gray matter regions. Each field of view (FOV) was classified based on its three dominant colors, excluding the lightest and darkest 20% of pixels to avoid artifacts from holes in tissue, as well as the cells and SST stainings themselves. The dominant colors were determined using k-means clustering (k = 3) and the resulting cluster centers were used as representative colors. A FOV was classified as gray matter if, in RGB space, the blue channel intensity exceeded the red channel by at least 25 units for any of the three dominant colors.

For FOVs identified as gray matter, color deconvolution was performed to separate the cellular component from the SST signal. Cell segmentation was conducted using a Cellpose machine learning model trained from scratch using human-in-the-loop training on several manually-annotated FOVs ^59^. The resultant cell masks were dilated by 50% of the cell’s radius to include surrounding cytoplasmic regions. The SST image was converted to grayscale, and pixel values below 90% of the median FOV pixel intensity were discarded to remove background noise. The remaining SST signal was thresholded based on size, retaining only regions larger than 20 pixels in area. Finally, colocalization of the thresholded SST signal and the dilated cell masks was performed, and any mask containing SST signal was classified as SST-positive.

### General procedures

Unless otherwise noted, data analysis was performed in Python, using the scanpy ^60^ ecosystem, plots were generated with matplotlib ^61^, and figures were assembled using Adobe Illustrator.

## ACKNOWLEDGEMENTS

With thanks to the CARE MND Register, hosted by the Euan Macdonald Centre for Motor Neuron Disease Research. This work was supported by awards from the National Institutes of Health: R01NS118183, R01NS116350, R01NS127186, and R01NS118570, as well as by a Target ALS grant (FD-2023-GEN-S1). The text of this manuscript was modified in part using a large language model (OpenAI; GPT-4o). This work utilized computational resources and assistance provided by the New York Genome Center. Additional computations were performed, at facilities run by the Scientific Computing Core at the Flatiron Institute, a division of the Simons Foundation. We thank Maria Hauge Pedersen and Nadia Propp for their contributions to project coordination. We are grateful to Luc Dupuis for his critical reading, discussions, and numerous suggestions.

## AUTHOR CONTRIBUTIONS

Conceptualization: HP, CS, JP, CGR, RB, JG; Methodology: JP, CGR, CAJ, AD, RB, KK, JE; Software: AD, CAJ, RB, JE; Validation: AD, CAJ, CGR, JE, JP, KK; Formal analysis: CAJ, CGR, AD, JE; Investigation: CGR, JP, KK, OC, ML, LR; Resources: HP, CS, KM; Data curation: AD, CAJ, CGR, JP, KK; Writing – original draft preparation: CGR, CAJ, AD, HP; Writing – review and editing: HP, CGR, CAJ, AD, JP, CS; Visualization: CAJ, CGR, AD, JP, JE; Supervision: HP; Project administration: HP; Funding acquisition: HP.

## DATA AVAILABILITY

Paired snRNA-seq and snATAC-seq data have been deposited in NCBI GEO (GSE290359) . Spatial transcriptomic and whole-genome sequencing data are available on the Target ALS portal.

## COMPETING INTEREST

Dr. Bonneau is a member of the Genentech and Roche leadership. The other authors declare no competing interests.

## Notes

### Competing Interest Statement

Dr. Richard Bonneau is a member of the Genentech and Roche leadership. The other authors declare no competing interests.

### Summary of Updates

New and updated analyses, in addition to manuscript text revisions.

## REFERENCES

1. Mead, R.J., Shan, N., Reiser, H.J., Marshall, F., and Shaw, P.J. (2023). Amyotrophic lateral sclerosis: a neurodegenerative disorder poised for successful therapeutic translation. Nat. Rev. Drug Discov. 22, 185–212.

2. Abrahams, S. (2023). Neuropsychological impairment in amyotrophic lateral sclerosis-frontotemporal spectrum disorder. Nat. Rev. Neurol. 19, 655–667.

3. Strong, M.J., Abrahams, S., Goldstein, L.H., Woolley, S., Mclaughlin, P., Snowden, J., Mioshi, E., Roberts-South, A., Benatar, M., HortobáGyi, T., et al. (2017). Amyotrophic lateral sclerosis - frontotemporal spectrum disorder (ALS-FTSD): Revised diagnostic criteria. Amyotroph. Lateral Scler. Frontotemporal Degener. 18, 153–174.

4. Abrahams, S., Goldstein, L.H., Kew, J.J., Brooks, D.J., Lloyd, C.M., Frith, C.D., and Leigh, P.N. (1996). Frontal lobe dysfunction in amyotrophic lateral sclerosis. A PET study. Brain 119 ( *Pt 6*), 2105–2120.

5. Gregory, J.M., McDade, K., Bak, T.H., Pal, S., Chandran, S., Smith, C., and Abrahams, S. (2020). Executive, language and fluency dysfunction are markers of localised TDP-43 cerebral pathology in non-demented ALS. J. Neurol. Neurosurg. Psychiatry 91, 149–157.

6. Kew, J.J., Goldstein, L.H., Leigh, P.N., Abrahams, S., Cosgrave, N., Passingham, R.E., Frackowiak, R.S., and Brooks, D.J. (1993). The relationship between abnormalities of cognitive function and cerebral activation in amyotrophic lateral sclerosis. A neuropsychological and positron emission tomography study. Brain 116 ( *Pt 6*), 1399–1423.

7. Abrahams, S., Goldstein, L.H., Simmons, A., Brammer, M.J., Williams, S.C.R., Giampietro, V.P., Andrew, C.M., and Leigh, P.N. (2003). Functional magnetic resonance imaging of verbal fluency and confrontation naming using compressed image acquisition to permit overt responses. Hum. Brain Mapp. 20, 29–40.

8. Abrahams, S., Goldstein, L.H., Simmons, A., Brammer, M., Williams, S.C.R., Giampietro, V., and Leigh, P.N. (2004). Word retrieval in amyotrophic lateral sclerosis: a functional magnetic resonance imaging study. Brain 127, 1507–1517.

9. Pineda, S.S., Lee, H., Ulloa-Navas, M.J., Linville, R.M., Garcia, F.J., Galani, K., Engelberg-Cook, E., Castanedes, M.C., Fitzwalter, B.E., Pregent, L.J., et al. (2024). Single-cell dissection of the human motor and prefrontal cortices in ALS and FTLD. Cell 187, 1971–1989.e16.

10. Yadav, A., Matson, K.J.E., Li, L., Hua, I., Petrescu, J., Kang, K., Alkaslasi, M.R., Lee, D.I., Hasan, S., Galuta, A., et al. (2023). A cellular taxonomy of the adult human spinal cord. Neuron 111, 328–344.e7.

11. Limone, F., Mordes, D.A., Couto, A., Joseph, B.J., Mitchell, J.M., Therrien, M., Ghosh, S.D., Meyer, D., Zhang, Y., Goldman, M., et al. (2024). Single-nucleus sequencing reveals enriched expression of genetic risk factors in extratelencephalic neurons sensitive to degeneration in ALS. Nat. Aging 4, 984–997.

12. Li, J., Jaiswal, M.K., Chien, J.-F., Kozlenkov, A., Jung, J., Zhou, P., Gardashli, M., Pregent, L.J., Engelberg-Cook, E., Dickson, D.W., et al. (2023). Divergent single cell transcriptome and epigenome alterations in ALS and FTD patients with C9orf72 mutation. Nat. Commun. 14, 5714.

13. Maniatis, S., Äijö, T., Vickovic, S., Braine, C., Kang, K., Mollbrink, A., Fagegaltier, D., Andrusivová, Ž., Saarenpää, S., Saiz-Castro, G., et al. (2019). Spatiotemporal dynamics of molecular pathology in amyotrophic lateral sclerosis. Science 364, 89–93.

14. Arnsten, A.F.T. (2015). Stress weakens prefrontal networks: molecular insults to higher cognition. Nat. Neurosci. 18, 1376–1385.

15. Gilbert, S.J., and Burgess, P.W. (2008). Executive function. Curr. Biol. 18, R110–R114.

16. Gregory, J.M., Elliott, E., McDade, K., Bak, T., Pal, S., Chandran, S., Abrahams, S., and Smith, C. (2020). Neuronal clusterin expression is associated with cognitive protection in amyotrophic lateral sclerosis. Neuropathol. Appl. Neurobiol. 46, 255–263.

17. Keller, S.S., Crow, T., Foundas, A., Amunts, K., and Roberts, N. (2009). Broca’s area: nomenclature, anatomy, typology and asymmetry. Brain Lang. 109, 29–48.

18. Fedorenko, E., Ivanova, A.A., and Regev, T.I. (2024). The language network as a natural kind within the broader landscape of the human brain. Nat. Rev. Neurosci. 25, 289–312.

19. Niven, E., Newton, J., Foley, J., Colville, S., Swingler, R., Chandran, S., Bak, T.H., and Abrahams, S. (2015). Validation of the Edinburgh Cognitive and Behavioural Amyotrophic Lateral Sclerosis Screen (ECAS): A cognitive tool for motor disorders. Amyotroph. Lateral Scler. Frontotemporal Degener. 16, 172–179.

20. Leighton, D., Newton, J., Colville, S., Bethell, A., Craig, G., Cunningham, L., Flett, M., Fraser, D., Hatrick, J., Lennox, H., et al. (2019). Clinical audit research and evaluation of motor neuron disease (CARE-MND): a national electronic platform for prospective, longitudinal monitoring of MND in Scotland. Amyotroph. Lateral Scler. Frontotemporal Degener. 20, 242–250.

21. Smith, C., and Millar, T. (2018). Brain donation procedures in the Sudden Death Brain Bank in Edinburgh. Handbook of clinical neurology 150. 10.1016/B978-0-444-63639-3.00002-5.

22. Taylor, L., Brown, R.G., Tsermentseli, S., Tsermentseli, S., Al-Chalabi, A., Shaw, C., Ellis, C., Leigh, P., and Goldstein, L. (2012). Is language impairment more common than executive dysfunction in amyotrophic lateral sclerosis? Journal of Neurology, Neurosurgery, and Psychiatry 84, 494–498.

23. Gittings, L.M., Alsop, E.B., Antone, J., Singer, M., Whitsett, T.G., Sattler, R., and Van Keuren-Jensen, K. (2023). Cryptic exon detection and transcriptomic changes revealed in single-nuclei RNA sequencing of C9ORF72 patients spanning the ALS-FTD spectrum. Acta Neuropathol. 146, 433–450.

24. Lau, S.-F., Cao, H., Fu, A.K.Y., and Ip, N.Y. (2020). Single-nucleus transcriptome analysis reveals dysregulation of angiogenic endothelial cells and neuroprotective glia in Alzheimer’s disease. Proceedings of the National Academy of Sciences 117, 25800–25809.

25. Sadick, J.S., O’Dea, M.R., Hasel, P., Dykstra, T., Faustin, A., and Liddelow, S.A. (2022). Astrocytes and oligodendrocytes undergo subtype-specific transcriptional changes in Alzheimer’s disease. Neuron 110, 1788–1805.e10.

26. Morabito, S., Miyoshi, E., Michael, N., Shahin, S., Martini, A.C., Head, E., Silva, J., Leavy, K., Perez-Rosendahl, M., and Swarup, V. (2021). Single-nucleus chromatin accessibility and transcriptomic characterization of Alzheimer’s disease. Nat. Genet. 53, 1143–1155.

27. Ma, S., Skarica, M., Li, Q., Xu, C., Risgaard, R.D., Tebbenkamp, A.T.N., Mato-Blanco, X., Kovner, R., Krsnik, Ž., de Martin, X., et al. (2022). Molecular and cellular evolution of the primate dorsolateral prefrontal cortex. Science 377, eabo7257.

28. Velmeshev, D., Schirmer, L., Jung, D., Haeussler, M., Perez, Y., Mayer, S., Bhaduri, A., Goyal, N., Rowitch, D.H., and Kriegstein, A.R. (2019). Single-cell genomics identifies cell type-specific molecular changes in autism. Science 364, 685–689.

29. Mathys, H., Peng, Z., Boix, C.A., Victor, M.B., Leary, N., Babu, S., Abdelhady, G., Jiang, X., Ng, A.P., Ghafari, K., et al. (2023). Single-cell atlas reveals correlates of high cognitive function, dementia, and resilience to Alzheimer’s disease pathology. Cell 186, 4365–4385.e27.

30. Kleshchevnikov, V., Shmatko, A., Dann, E., Aivazidis, A., King, H.W., Li, T., Elmentaite, R., Lomakin, A., Kedlian, V., Gayoso, A., et al. (2022). Cell2location maps fine-grained cell types in spatial transcriptomics. Nat. Biotechnol. 40, 661–671.

31. Yang, A.C., Vest, R.T., Kern, F., Lee, D.P., Agam, M., Maat, C.A., Losada, P.M., Chen, M.B., Schaum, N., Khoury, N., et al. (2022). A human brain vascular atlas reveals diverse mediators of Alzheimer’s risk. Nature 603, 885–892.

32. Huuki-Myers, L.A., Spangler, A., Eagles, N.J., Montgomery, K.D., Kwon, S.H., Guo, B., Grant-Peters, M., Divecha, H.R., Tippani, M., Sriworarat, C., et al. (2024). A data-driven single-cell and spatial transcriptomic map of the human prefrontal cortex. Science 384, eadh1938.

33. Maynard, K.R., Collado-Torres, L., Weber, L.M., Uytingco, C., Barry, B.K., Williams, S.R., Catallini, J.L., 2nd, Tran, M.N., Besich, Z., Tippani, M., et al. (2021). Transcriptome-scale spatial gene expression in the human dorsolateral prefrontal cortex. Nat. Neurosci. 24, 425–436.

34. Chen, S., Chang, Y., Li, L., Acosta, D., Li, Y., Guo, Q., Wang, C., Turkes, E., Morrison, C., Julian, D., et al. (2022). Spatially resolved transcriptomics reveals genes associated with the vulnerability of middle temporal gyrus in Alzheimer’s disease. Acta Neuropathol. Commun. 10, 188.

35. Song, L., Chen, W., Hou, J., Guo, M., and Yang, J. (2025). Spatially resolved mapping of cells associated with human complex traits. Nature. 10.1038/s41586-025-08757-x.

36. Äijö, T., Maniatis, S., Vickovic, S., Kang, K., Cuevas, M., Braine, C., Phatnani, H., Lundeberg, J., and Bonneau, R. (2019). Splotch: Robust estimation of aligned spatial temporal gene expression data. bioRxiv, 757096. 10.1101/757096.

37. Daly, A.C., Cambuli, F., Äijö, T., Lötstedt, B., Marjanovic, N., Kuksenko, O., Smith-Erb, M., Fernandez, S., Domovic, D., Van Wittenberghe, N., et al. (2024). Tissue and cellular spatiotemporal dynamics in colon aging. bioRxiv, 2024.04.22.590125. 10.1101/2024.04.22.590125.

38. Ferguson, H.J., Brunsdon, V.E.A., and Bradford, E.E.F. (2021). The developmental trajectories of executive function from adolescence to old age. Sci. Rep. 11, 1382.

39. Idowu, M.I., and Szameitat, A.J. (2023). Executive function abilities in cognitively healthy young and older adults-A cross-sectional study. Front. Aging Neurosci. 15, 976915.

40. Gabitto, M.I., Travaglini, K.J., Rachleff, V.M., Kaplan, E.S., Long, B., Ariza, J., Ding, Y., Mahoney, J.T., Dee, N., Goldy, J., et al. (2024). Integrated multimodal cell atlas of Alzheimer’s disease. Nat. Neurosci. 27, 2366–2383.

41. Fisher, J., Verhagen, M., Long, Z., Moissidis, M., Yan, Y., He, C., Wang, J., Micoli, E., Alastruey, C.M., Moors, R., et al. (2024). Cortical somatostatin long-range projection neurons and interneurons exhibit divergent developmental trajectories. Neuron 112, 558–573.e8.

42. Suk, T.R., and Rousseaux, M.W.C. (2020). The role of TDP-43 mislocalization in amyotrophic lateral sclerosis. Mol. Neurodegener. 15, 45.

43. Ling, and Shuo-Chien Synaptic Paths to Neurodegeneration: The Emerging Role of TDP-43 and FUS in Synaptic Functions. Preprint at Hindawi, 10.1155/2018/8413496 10.1155/2018/8413496.

44. Wang, W., Wang, L., Lu, J., Siedlak, S.L., Fujioka, H., Liang, J., Jiang, S., Ma, X., Jiang, Z., da Rocha, E.L., et al. (2016). The inhibition of TDP-43 mitochondrial localization blocks its neuronal toxicity. Nat. Med. 22, 869–878.

45. Tam, O.H., Rozhkov, N.V., Shaw, R., Kim, D., Hubbard, I., Fennessey, S., Propp, N., The NYGC ALS Consortium, Phatnani, H., Kwan, J., et al. (2019). Postmortem Cortex Samples Identify Distinct Molecular Subtypes of ALS: Retrotransposon Activation, Oxidative Stress, and Activated Glia. CellReports 29, 1164–1177.e5.

46. Helmold, B.R., Pauss, K.E., and Ozdinler, P.H. (2023). TDP-43 protein interactome informs about perturbed canonical pathways and may help develop personalized medicine approaches for patients with TDP-43 pathology. Drug Discov. Today 28, 103769.

47. Liu, E.Y., Russ, J., Cali, C.P., Phan, J.M., Amlie-Wolf, A., and Lee, E.B. (2019). Loss of nuclear TDP-43 is associated with decondensation of LINE retrotransposons. Cell Rep. 27, 1409–1421.e6.

48. Spence, H., Waldron, F.M., Saleeb, R.S., Brown, A.-L., Rifai, O.M., Gilodi, M., Read, F., Roberts, K., Milne, G., Wilkinson, D., et al. (2024). RNA aptamer reveals nuclear TDP-43 pathology is an early aggregation event that coincides with STMN-2 cryptic splicing and precedes clinical manifestation in ALS. Acta Neuropathol. 147, 50.

49. Burda, J.E., and Sofroniew, M.V. (2014). Reactive gliosis and the multicellular response to CNS damage and disease. Neuron 81, 229–248.

50. Jackson, C.E., McVey, A.L., Rudnicki, S., Dimachkie, M.M., and Barohn, R.J. (2015). Symptom management and end-of-life care in amyotrophic lateral sclerosis. Neurol. Clin. 33, 889–908.

51. Abrahams, S., Newton, J., Niven, E., Foley, J., and Bak, T.H. (2014). Screening for cognition and behaviour changes in ALS. Amyotroph. Lateral Scler. Frontotemporal Degener. 15, 9–14.

52. Blanchard, J.W., Akay, L.A., Davila-Velderrain, J., von Maydell, D., Mathys, H., Davidson, S.M., Effenberger, A., Chen, C.-Y., Maner-Smith, K., Hajjar, I., et al. (2022). APOE4 impairs myelination via cholesterol dysregulation in oligodendrocytes. Nature 611, 769–779.

53. Mathys, H., Davila-Velderrain, J., Peng, Z., Gao, F., Mohammadi, S., Young, J.Z., Menon, M., He, L., Abdurrob, F., Jiang, X., et al. (2019). Single-cell transcriptomic analysis of Alzheimer’s disease. Nature 570, 332–337.

54. Muzellec, B., Teleńczuk, M., Cabeli, V., and Andreux, M. (2023). PyDESeq2: a python package for bulk RNA-seq differential expression analysis. Bioinformatics 39, btad547.

55. Anders, S., and Huber, W. (2010). Differential expression analysis for sequence count data. Genome Biol. 11, R106.

56. Zhou, Y., Zhou, B., Pache, L., Chang, M., Khodabakhshi, A.H., Tanaseichuk, O., Benner, C., and Chanda, S.K. (2019). Metascape provides a biologist-oriented resource for the analysis of systems-level datasets. Nat. Commun. 10, 1523.

57. Kolberg, L., Raudvere, U., Kuzmin, I., Adler, P., Vilo, J., and Peterson, H. (2023). g:Profiler-interoperable web service for functional enrichment analysis and gene identifier mapping (2023 update). Nucleic Acids Res. 51, W207–W212.

58. Wagenmakers, E.-J., Lodewyckx, T., Kuriyal, H., and Grasman, R. (2010). Bayesian hypothesis testing for psychologists: a tutorial on the Savage-Dickey method. Cogn. Psychol. 60, 158–189.

59. Pachitariu, M., and Stringer, C. (2022). Cellpose 2.0: how to train your own model. Nat. Methods 19, 1634–1641.

60. Wolf, F.A., Angerer, P., and Theis, F.J. (2018). SCANPY: large-scale single-cell gene expression data analysis. Genome Biol. 19, 15.

61. Hunter, J.D. (2007). Matplotlib: A 2D Graphics Environment. Computing in Science Engineering 9, 90–95.

