## Supplemental Figures for "Distinct Cellular Phenotypes of Language and Executive Decline in Amyotrophic Lateral Sclerosis"

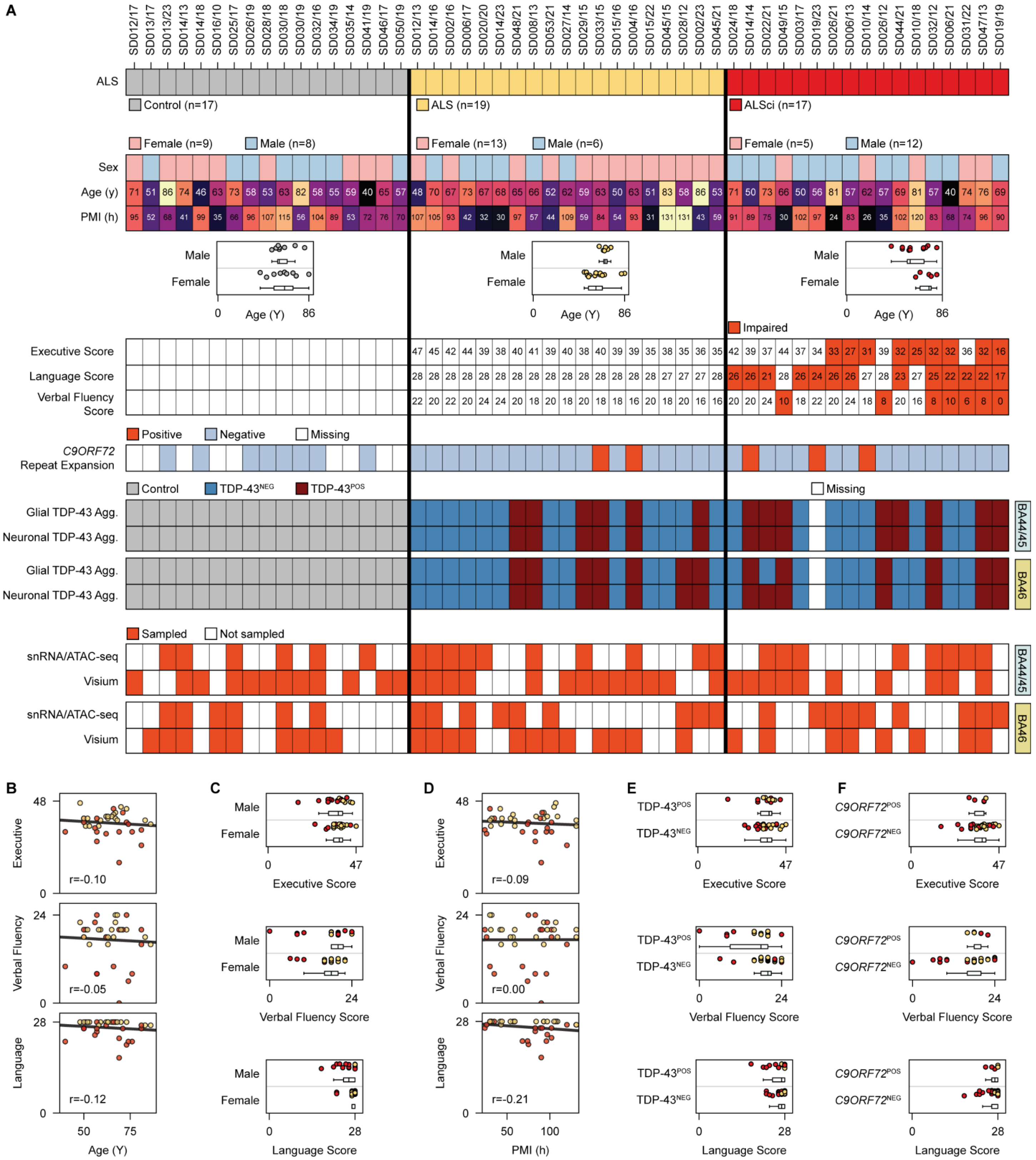
A
SD012/17 SD013/17 SD013/23 SD014/13 SD014/18 SD016/10 SD025/17 SD026/19 SD028/18 SD030/18 SD030/19 SD032/16 SD034/19 SD035/14 SD041/19 SD046/17 SD050/19 SD012/13 SD014/16 SD002/16 SD006/17 SD020/20 SD014/23 SD048/21 SD008/13 SD053/21 SD027/14 SD029/15 SD033/15 SD015/16 SD004/16 SD015/22 SD045/15 SD028/12 SD002/23 SD045/21 SD024/18 SD014/14 SD022/21 SD049/15 SD003/17 SD019/23 SD026/21 SD006/13 SD010/14 SD026/12 SD044/21 SD010/18 SD006/21 SD031/22 SD047/13 SD019/19
ALS
Control (n=17)
ALS (n=19)
ALSci (n=17)
Female (n=9)
Male (n=8)
Female (n=13)
Male (n=6)
Female (n=5)
Male (n=12)
Sex
Age (y)
PMI (h)
Male
Female
0
Age (Y)
86
Impaired
Executive Score
Language Score
Verbal Fluency Score
Positive
Negative
Missing
C9ORF72 Repeat Expansion
Control
TDP-43NEG
TDP-43POS
Missing
Glial TDP-43 Agg.
Neuronal TDP-43 Agg.
BA44/45
BA46
Sampled
Not sampled
snRNA/ATAC-seq
Visium
B
Executive
Verbal Fluency
Language
r=-0.10
r=-0.05
r=-0.12
Age (Y)
C
Executive Score
Verbal Fluency Score
Language Score
D
r=-0.09
r=0.00
r=-0.21
PMI (h)
E
TDP-43POS
TDP-43NEG
F
C9ORF72POS
C9ORF72NEG

**Supplemental Figure 1: Donor cohort metadata and individual correlation between ECAS scores and donor phenotypes.**

**A)** Each column represents a postmortem donor, labeled by donor ID at the top and annotated with the following metadata from top to bottom:1) ALS disease state; 2) donor sex; 3) donor age at death (years); 4) postmortem interval (hours); 5) ECAS executive score (scores of 33 or less are highlighted as impaired); 6) ECAS language score (scores of 26 or less are highlighted as impaired); 7) ECAS verbal fluency score (scores of 14 or less are highlighted as impaired); 8) *C9ORF72* repeat expansion status, as determined by whole-genome sequencing; 9) presence of TDP-43 aggregates in BA44/45 glial cells, assessed by immunohistochemistry; 10) presence of TDP-43 aggregates in BA44/45 neuronal cells; 11) presence of TDP-43 aggregates in BA46 glial cells; and 12) presence of TDP-43 aggregates in BA46 neuronal cells.

**B**) Donor age at death plotted against ECAS scores for executive function, verbal fluency, and language (top to bottom). The black line represents the ordinary least squares best-fit line; *r* is the Pearson correlation coefficient.

**C**) Donor sex plotted against ECAS scores for executive, verbal fluency, and language (top to bottom).

**D**) Postmortem interval (hours) plotted against ECAS scores as in panel B.

**E**) TDP-43 aggregation status plotted against ECAS scores as in panel C.

**F**) Donor *C9ORF72* repeat expansion status plotted against ECAS scores as in panel C.

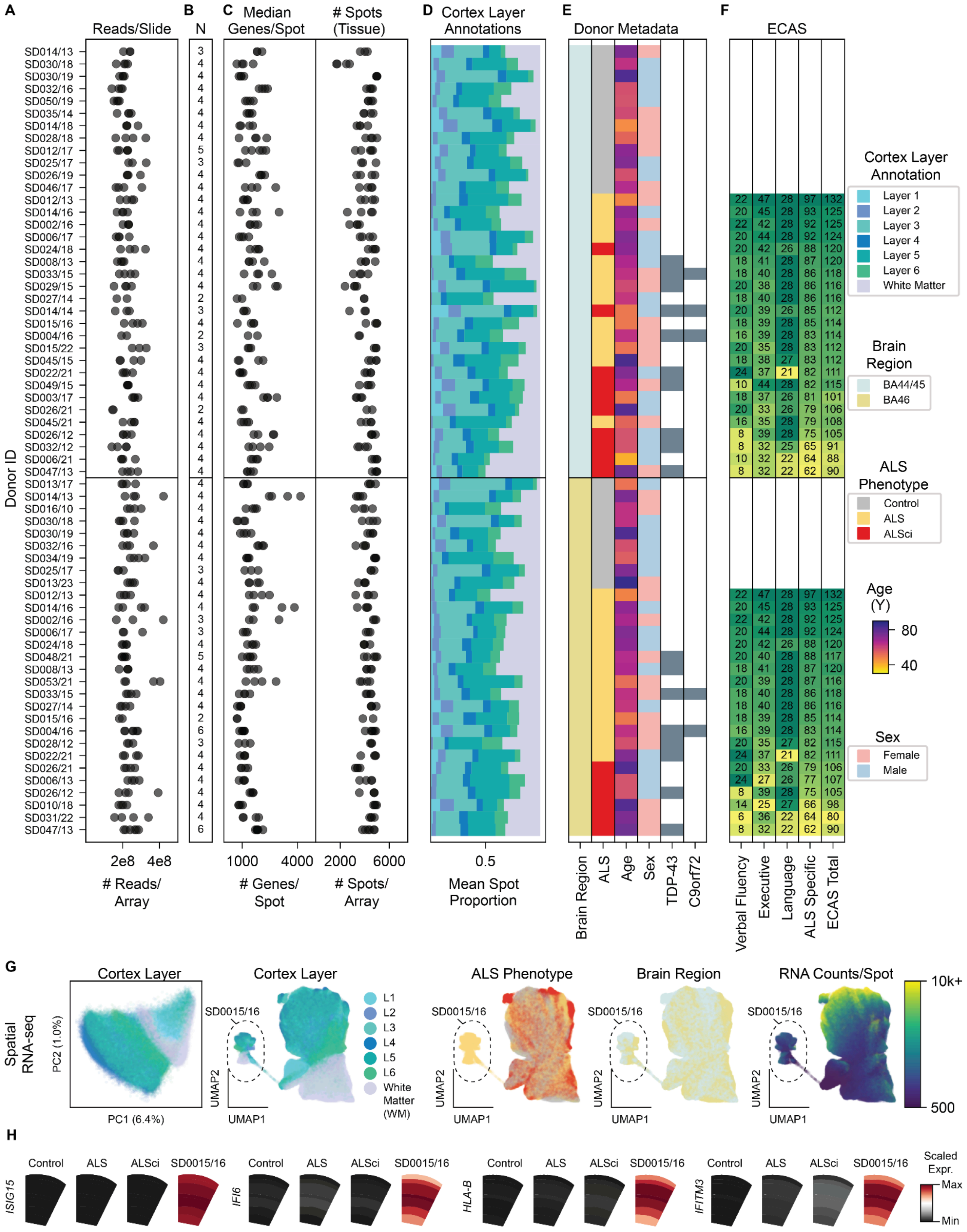
A
Reads/Slide
B
N
C
Median Genes/Spot
### Spots (Tissue)
D
Cortex Layer Annotations
E
Donor Metadata
F
ECAS
Donor ID
SD014/13
SD030/18
SD030/19
SD032/16
SD050/19
SD035/14
SD014/18
SD028/18
SD012/17
SD025/17
SD026/19
SD046/17
SD012/13
SD014/16
SD002/16
SD006/17
SD024/18
SD008/13
SD033/15
SD029/15
SD027/14
SD014/14
SD015/16
SD004/16
SD015/22
SD045/15
SD022/21
SD049/15
SD003/17
SD026/21
SD045/21
SD026/12
SD032/12
SD006/21
SD047/13
SD013/17
SD014/13
SD016/10
SD030/18
SD030/19
SD032/16
SD034/19
SD025/17
SD013/23
SD012/13
SD014/16
SD002/16
SD006/17
SD024/18
SD048/21
SD008/13
SD053/21
SD033/15
SD027/14
SD015/16
SD004/16
SD028/12
SD022/21
SD026/21
SD006/13
SD026/12
SD010/18
SD031/22
SD047/13
2e8 4e8
### Reads/ Array
1000 4000
### Genes/ Spot
2000 6000
### Spots/ Array
0.5
Mean Spot Proportion
Brain Region
ALS
Age
Sex
TDP-43
C9orf72
Verbal Fluency
Executive
Language
ALS Specific
ECAS Total
Cortex Layer Annotation
Layer 1
Layer 2
Layer 3
Layer 4
Layer 5
Layer 6
White Matter
Brain Region
BA44/45
BA46
ALS Phenotype
Control
ALS
ALSci
Age (Y)
80
40
Sex
Female
Male
G
Spatial RNA-seq
Cortex Layer
PC2 (1.0%)
PC1 (6.4%)
Cortex Layer
SD0015/16
UMAP2
UMAP1
L1
L2
L3
L4
L5
L6
White Matter (WM)
ALS Phenotype
Brain Region
RNA Counts/Spot
10k+
500
H
Control
ALS
ALSci
SD0015/16
ISG15
IFI6
HLA-B
IFITM3
Scaled Expr.
Max
Min

**Supplemental Figure S2: Quality control and layer annotation for Visium ST libraries.**

Each row is a single biological sample from a donor (SD ID denotes anonymized donor identification) and summarizes the following information:

**A**) Number of paired-end reads per Visium array.

**B**) Number of Visium arrays generated per donor.

**C**) Median number of genes per spot reported by spaceranger, and number of spots which covered by tissue in each array.

**D**) Mean proportion of spots annotated per cortical layer.

**E**) Sample metadata, including brain region (BA44/45 or BA46), donor ALS phenotype, age, sex, TDP-43 pathology status, and *C9orf72* repeat expansion genotype.

**F**) Edinburgh Cognitive and Behavioural ALS Screen (ECAS) scores for ALS donors, broken down into Verbal Fluency, Executive, and Language scores. These are summed to produce the ALS-Specific score. The total ECAS score incorporates the ALS-Specific score plus Visuospatial and Memory scores (both not shown).

**G**) PCA and UMAP plots for individual Visium array spots, colored by annotated cortical layer, ALS phenotype, brain region, and RNA content (UMI counts per spot).

**H**) Expression of interferon-induced and HLA transcripts in outlier donor (SD0015/16).

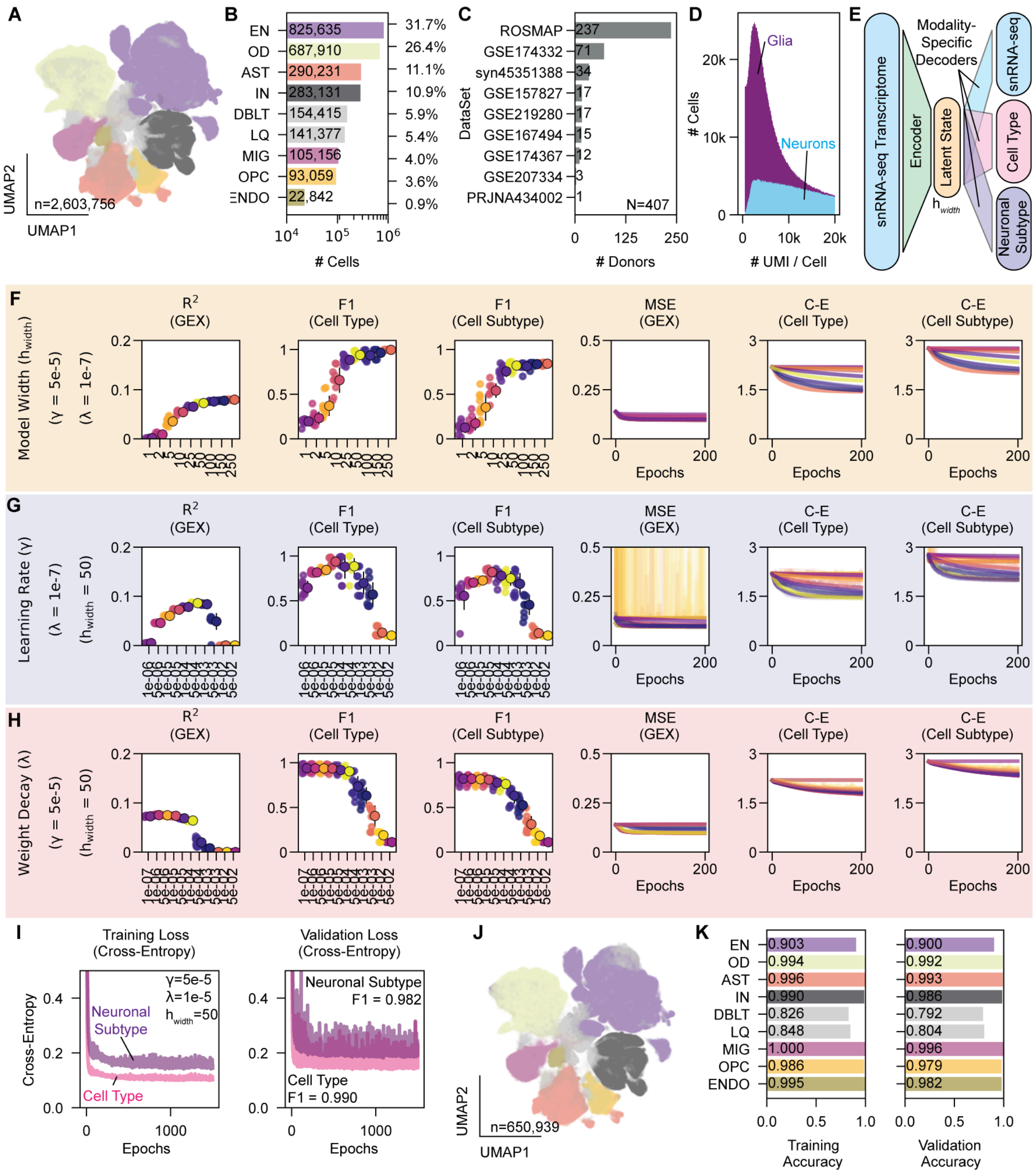
A
UMAP2
UMAP1
n=2,603,756
B
EN 825,635 31.7%
OD 687,910 26.4%
AST 290,231 11.1%
IN 283,131 10.9%
DBLT 154,415 5.9%
LQ 141,377 5.4%
MIG 105,156 4.0%
OPC 93,059 3.6%
ENDO 22,842 0.9%
### Cells
C
DataSet
ROSMAP 237
GSE174332 71
syn45351388 34
GSE157827 17
GSE219280 17
GSE167494 15
GSE174367 12
GSE207334 3
PRJNA434002 1
N=407
### Donors
D
Glia
Neurons
### Cells
### UMI / Cell
E
snRNA-seq Transcriptome
Encoder
Latent State
h_width
Modality-Specific Decoders
snRNA-seq
Cell Type
Neuronal Subtype
F
Model Width (h_width)
(γ = 5e-5)
(λ = 1e-7)
R² (GEX)
F1 (Cell Type)
F1 (Cell Subtype)
MSE (GEX)
C-E (Cell Type)
C-E (Cell Subtype)
Epochs
G
Learning Rate (γ)
(λ = 1e-7)
(h_width = 50)
H
Weight Decay (λ)
(γ = 5e-5)
(h_width = 50)
I
Training Loss (Cross-Entropy)
Validation Loss (Cross-Entropy)
Cross-Entropy
γ=5e-5
λ=1e-5
h_width=50
Neuronal Subtype
Cell Type
Neuronal Subtype F1 = 0.982
Cell Type F1 = 0.990
Epochs
J
UMAP2
UMAP1
n=650,939
K
EN 0.903 0.900
OD 0.994 0.992
AST 0.996 0.993
IN 0.990 0.986
DBLT 0.826 0.792
LQ 0.848 0.804
MIG 1.000 0.996
OPC 0.986 0.979
ENDO 0.995 0.982
Training Accuracy
Validation Accuracy

**Supplemental Figure S3: Assembly of a published prefrontal cortex snRNA-seq cohort and training of a machine learning classifier to label cell types.**

**A**) UMAP plot of 2.6 million cells from 407 donor samples, colored by annotated broad cell type.

**B**) The number of cells for the seven broad cell types, plus two technical categories (Low-Quality, LQ; and Doublet, DBLT).

**C**) The number of donors which meet the inclusion criteria from each originating data set.

**D**) Distribution of glial and neuronal cell UMI counts across the cohort.

**E**) Schematic of multimodal autoencoder, with a unified encoder and modality-specific decoders into gene expression, cell type, and cell (neuronal) subtype.

**F**) Grid search for number of nodes in the narrowest layer embedding ($h_{width}$), holding model optimizer (Adam) learning rate and weight decay constant. All models are regularized during training with a 50% dropout rate on the input layer and 25% dropout on each hidden layer. Models are evaluated using held-out cross-validation data, on recovery of transcript expression ($R^2$), on correct identification of cell type (F1 score), and on correct identification of cell subtype (F1 score). Model losses for transcript expression (Mean Squared Error, MSE) and for the cell type and subtype classifiers (Cross-entropy, C-E) are plotted against training epochs.

**G**) Grid search for model optimizer learning rate ($\gamma$), as in Supplemental Figure S3F, holding model width and weight decay constant.

**H**) Grid search for model optimizer weight decay ($\boldsymbol{\lambda}$), as in Supplemental Figure S3F, holding model width and learning rate constant.

**I**) Training and validation losses for the classifiers in the full model trained on 1.95 million cells. Validation F1 scores are annotated.

**J**) UMAP plot, as in Supplemental Figure S3A, of 650k validation cells held out of the training of the full model. Cells are colored by the predicted cell type.

**K**) Cell type model accuracy for training and validation data.

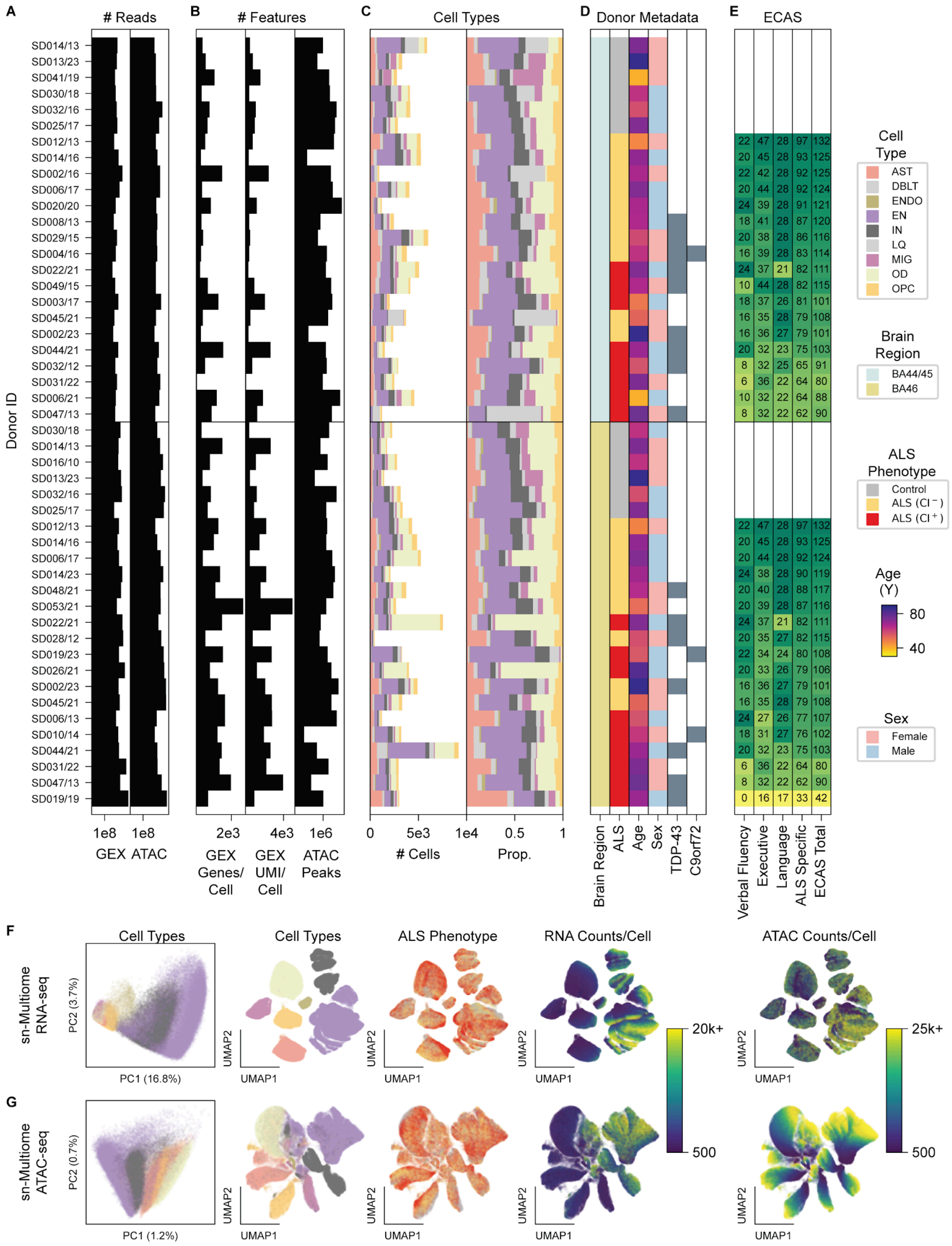
A
### Reads
B
### Features
C
Cell Types
D
Donor Metadata
E
ECAS
Donor ID
SD014/13
SD013/23
SD041/19
SD030/18
SD032/16
SD025/17
SD012/13
SD014/16
SD002/16
SD006/17
SD020/20
SD008/13
SD029/15
SD004/16
SD022/21
SD049/15
SD003/17
SD045/21
SD002/23
SD044/21
SD032/12
SD031/22
SD006/21
SD047/13
SD030/18
SD014/13
SD016/10
SD013/23
SD032/16
SD025/17
SD012/13
SD014/16
SD006/17
SD014/23
SD048/21
SD053/21
SD022/21
SD028/12
SD019/23
SD026/21
SD002/23
SD045/21
SD006/13
SD010/14
SD044/21
SD031/22
SD047/13
SD019/19
1e8
1e8
GEX ATAC
2e3
4e3
1e6
GEX Genes/ Cell
GEX UMI/ Cell
ATAC Peaks
0
5e3
1e4
0.5
1
### Cells
Prop.
Brain Region
ALS
Age
Sex
TDP-43
C9orf72
Verbal Fluency
Executive
Language
ALS Specific
ECAS Total
Cell Type
AST
DBLT
ENDO
EN
IN
LQ
MIG
OD
OPC
Brain Region
BA44/45
BA46
ALS Phenotype
Control
ALS (CI⁻)
ALS (CI⁺)
Age (Y)
80
40
Sex
Female
Male
F
sn-Multiome RNA-seq
Cell Types
PC2 (3.7%)
PC1 (16.8%)
Cell Types
ALS Phenotype
RNA Counts/Cell
ATAC Counts/Cell
UMAP2
UMAP1
20k+
500
25k+
500
G
sn-Multiome ATAC-seq
PC2 (0.7%)
PC1 (1.2%)

**Supplemental Figure S4: Quality control and cell typing for Multiome libraries.**

Each row is a single biological sample from a donor (SD ID denotes anonymized donor identification) and summarizes the following information:

**A**) Number of paired-end reads for single cell gene expression (GEX) and assay for transposase accessible chromatin (ATAC) libraries.

**B**) Median number of genes per cell in GEX and median number of UMI counts per cell in GEX, both reported after Cell Ranger ARC processing but prior to additional filtering or quality control. Also shown is the number of quantifiable ATAC peaks per library.

**C**) Broad cell types, including doublets (DBLT) and low-quality cells (LQ) called by cell type machine learning model. Cell type distributions are plotted as total cell counts (left) and as proportion of the total (right).

**D**) Sample metadata, including brain region (BA44/45 or BA46), donor ALS phenotype, age, sex, TDP-43 pathology status, and *C9orf72* repeat expansion genotype.

**E**) Edinburgh Cognitive and Behavioural ALS Screen (ECAS) scores for ALS donors, broken down into Verbal Fluency, Executive, and Language scores. These are summed to produce the ALS-Specific score. The Total ECAS score incorporates the ALS-Specific score plus Visuospatial and Memory scores (both not shown).

**F**) PCA and UMAP plots of the snRNA-seq component of Multiome data, colored by cell type, ALS phenotype, RNA UMI counts per cell, and ATAC peak fragment counts per cell.

**G**) PCA and UMAP plots of the snATAC-seq component of Multiome data, colored as in Supplemental Figure S4F.

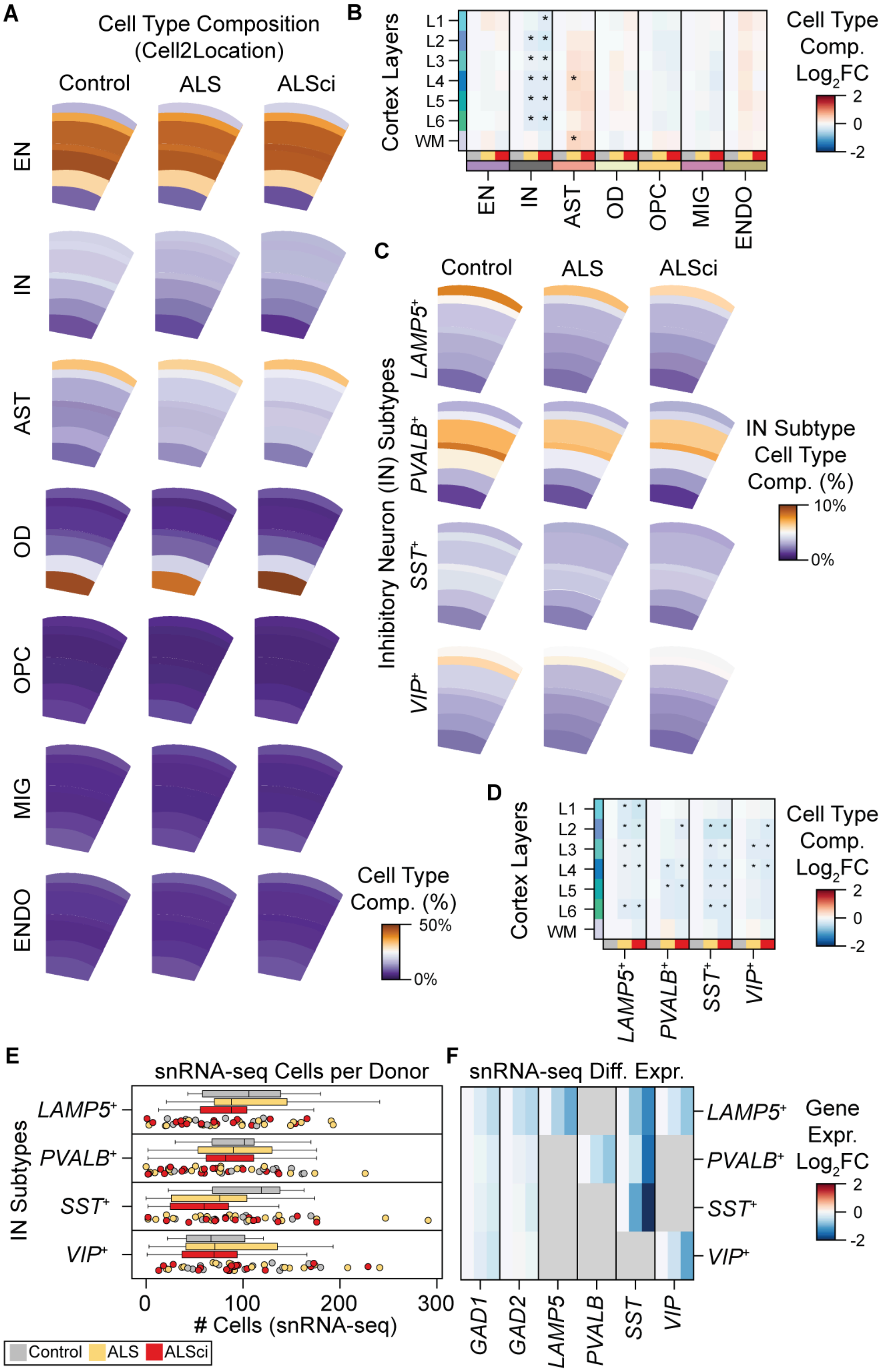

A
Cell Type Composition (Cell2Location)
Control
ALS
ALSci
EN
IN
AST
OD
OPC
MIG
ENDO
Cell Type Comp. (%)
50%
0%
B
Cortex Layers
L1
L2
L3
L4
L5
L6
WM
EN
IN
AST
OD
OPC
MIG
ENDO
Cell Type Comp. Log2FC
2
0
-2
C
Control
ALS
ALSci
Inhibitory Neuron (IN) Subtypes
LAMP5+
PVALB+
SST+
VIP+
IN Subtype Cell Type Comp. (%)
10%
0%
D
Cortex Layers
L1
L2
L3
L4
L5
L6
WM
LAMP5+
PVALB+
SST+
VIP+
Cell Type Comp. Log2FC
2
0
-2
E
snRNA-seq Cells per Donor
IN Subtypes
LAMP5+
PVALB+
SST+
VIP+
0
100
200
300
### Cells (snRNA-seq)
Control
ALS
ALSci
F
snRNA-seq Diff. Expr.
LAMP5+
PVALB+
SST+
VIP+
GAD1
GAD2
LAMP5
PVALB
SST
VIP
Gene Expr. Log2FC
2
0
-2

**Supplemental Figure S5: Cell type distribution in ST data.**

**A**) Broad cell types deconvoluted from spatial data using cell2location model built from 2.6 million annotated single nuclei. Colors represent the mean proportion of each cell type relative to the total number of cells in each cortical layer (totals may not add to exactly 100% due to rounding and averaging).

**B**) Changes in broad cell composition in ALS and ALSci donors compared to controls, colored by $\log_2$ fold change. Significant differences are marked with asterisks (Welch's *t*-test, $p < 0.01$; false discovery rate controlled at 0.01).

**C**) Inhibitory neuron (IN) subtypes deconvoluted from spatial data using cell2location. Colors represent the mean proportion of IN cells of that subtype relative to the total number of cells in the layer. Note that data are displayed on a different scale than those in Supplemental Figure S5A.

**D**) Changes in IN subtype composition in ALS and ALSci donors compared to controls, colored by $\log_2$ fold change. Significant differences are marked with asterisks (Welch's *t*-test, $p < 0.01$; false discovery rate controlled at 0.01).

**E**) Number of snRNA-seq nuclei annotated as each IN subtype per donor, colored by ALS phenotype (control, ALS, or ALSci). Individual donor measurements are displayed.

**F**) Expression of IN marker genes, shown as $\log_2$ fold change relative to the control donors. Genes with very low expression in the given IN subtype are grayed out.

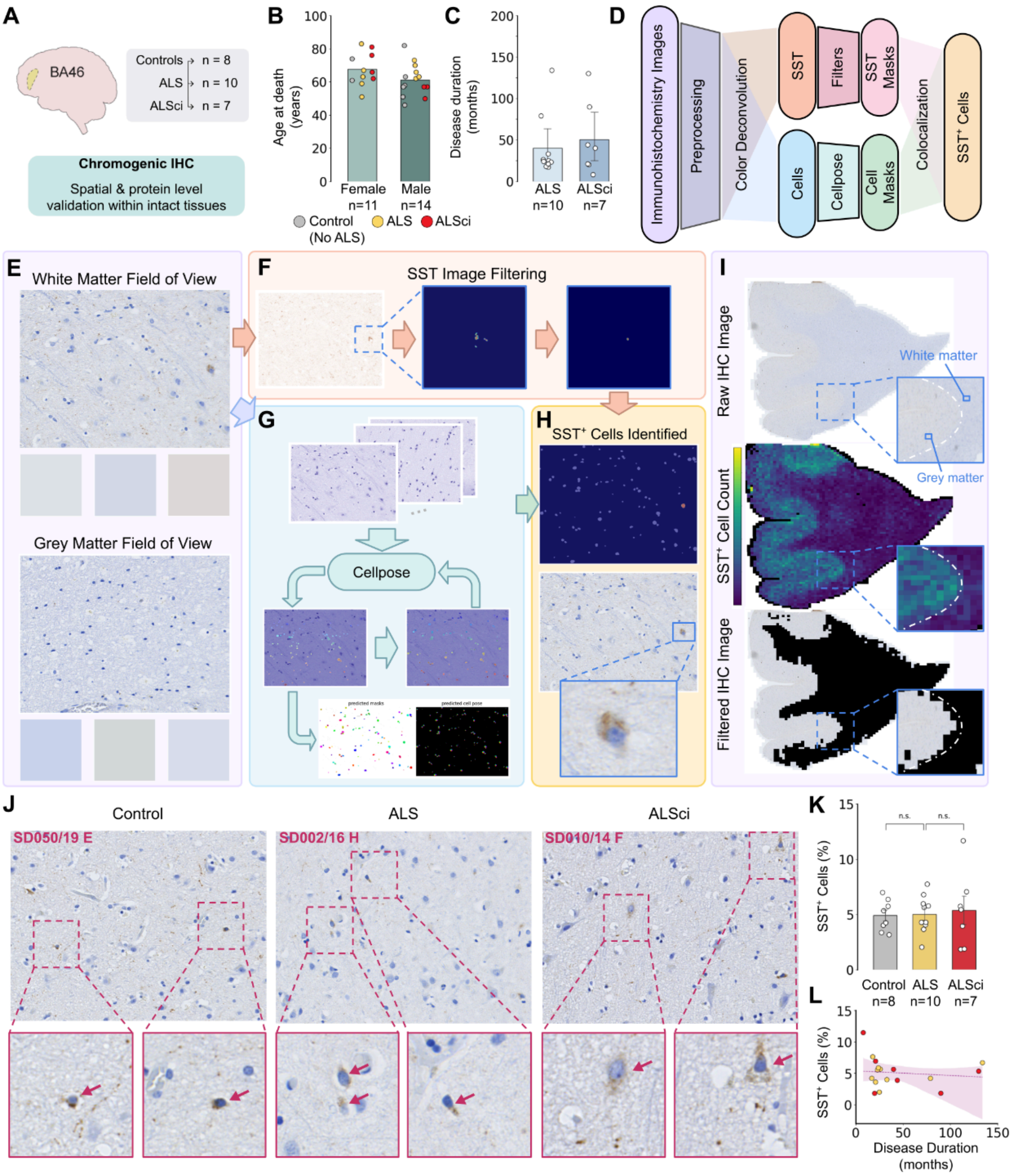
A
BA46
Controls n = 8
ALS n = 10
ALSci n = 7
Chromogenic IHC
Spatial & protein level validation within intact tissues
B
Age at death (years)
Female n=11
Male n=14
Control (No ALS)
ALS
ALSci
C
Disease duration (months)
ALS n=10
ALSci n=7
D
Immunohistochemistry Images
Preprocessing
Color Deconvolution
SST
Filters
SST Masks
Cells
Cellpose
Cell Masks
Colocalization
SST⁺ Cells
E
White Matter Field of View
Grey Matter Field of View
F
SST Image Filtering
G
Cellpose
H
SST⁺ Cells Identified
I
Raw IHC Image
White matter
Grey matter
SST⁺ Cell Count
Filtered IHC Image
J
Control
ALS
ALSci
SD050/19 E
SD002/16 H
SD010/14 F
K
SST⁺ Cells (%)
n.s.
n.s.
Control n=8
ALS n=10
ALSci n=7
L
SST⁺ Cells (%)
Disease Duration (months)

**Supplemental Figure S6: SST$^+$ cell proportions are unchanged in ALS donors.**

**A**) BA46 donor tissue blocks (*n* = 25), including ALS, ALSci, and control donors, were cryosectioned and stained with an anti-SST antibody using chromogenic immunohistochemistry (IHC) methods.

**B**) Donor age at death separated by sex and cognitive status.

**C**) The interval between diagnosis and date of death (disease duration) for ALS donors. Error bars indicate 95% confidence intervals.

**D**) Schematic of the Cellpose machine learning pipeline for cell segmentation and SST$^+$ cell classification.

**E**) Representative white and gray matter fields of view from a control donor.

**F**) Filtering of images to isolate SST$^+$ signal regions.

**G**) Human-in-the-loop fine-tuning of the Cellpose model for image segmentation, cell counting, and masking.

**H**) Colocalization of SST$^+$ signal regions with Cellpose-generated masks to identify SST$^+$ cells.

**I**) Filtering of white matter from IHC images to enable gray matter–specific quantification.

**J**) Representative fields of view from control, ALS, and ALSci donors showing SST$^+$ cells after masking, segmentation, and classification.

**K**) Percentage of SST$^+$ cells identified for each of the 25 donors. No statistically significant differences were observed (Welch's *t*-test). Error bars indicate 95% confidence intervals.

**L**) Percentage of SST$^+$ cells per donor plotted against ALS disease duration. The slope of the ordinary least squares best fit line is not significantly different from zero (Wald test).

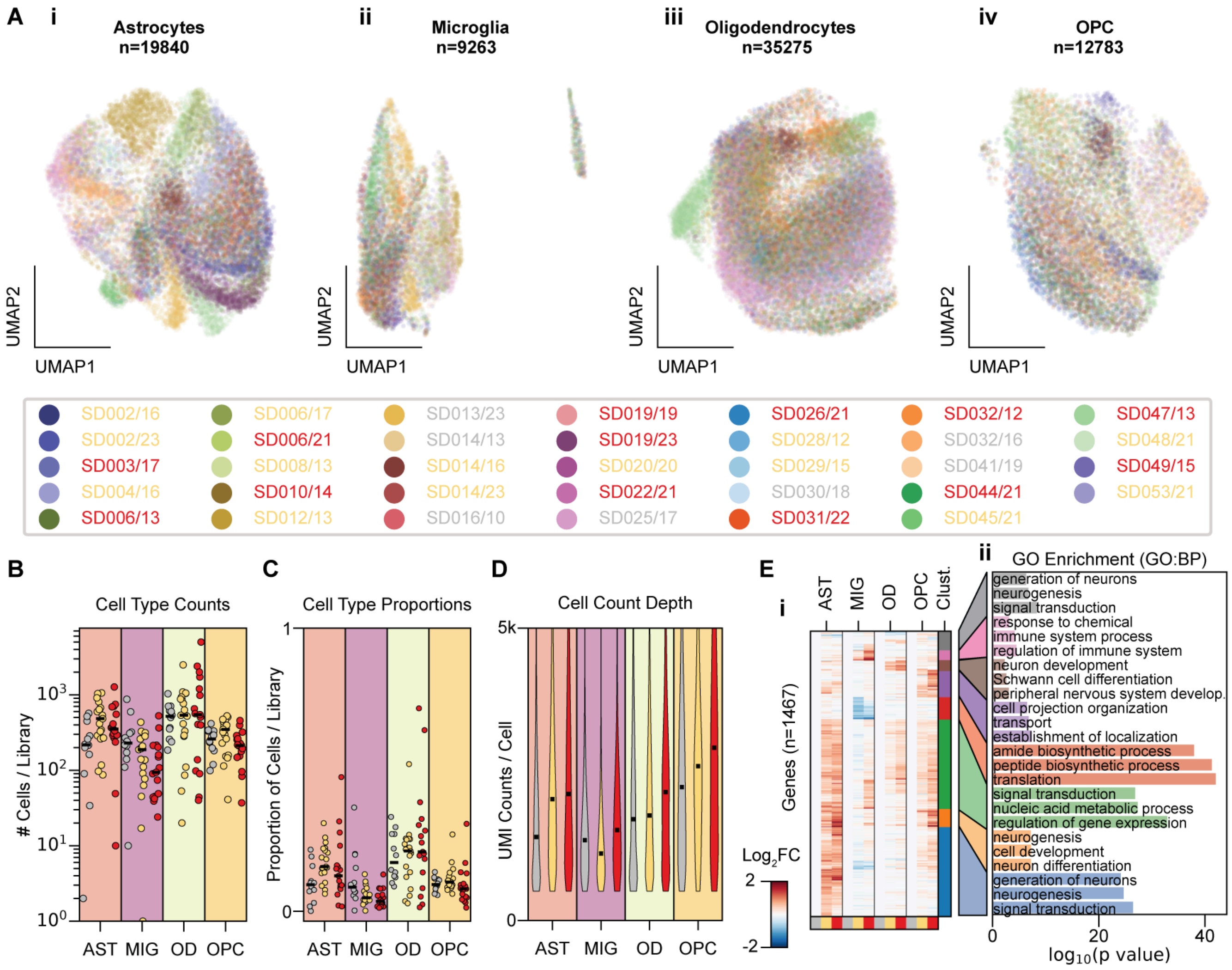
A
i
Astrocytes
n=19840
ii
Microglia
n=9263
iii
Oligodendrocytes
n=35275
iv
OPC
n=12783
UMAP2
UMAP1
SD002/16
SD002/23
SD003/17
SD004/16
SD006/13
SD006/17
SD006/21
SD008/13
SD010/14
SD012/13
SD013/23
SD014/13
SD014/16
SD014/23
SD016/10
SD019/19
SD019/23
SD020/20
SD022/21
SD025/17
SD026/21
SD028/12
SD029/15
SD030/18
SD031/22
SD032/12
SD032/16
SD041/19
SD044/21
SD045/21
SD047/13
SD048/21
SD049/15
SD053/21
B
Cell Type Counts
### Cells / Library
AST MIG OD OPC
C
Cell Type Proportions
Proportion of Cells / Library
AST MIG OD OPC
D
Cell Count Depth
UMI Counts / Cell
5k
0
AST MIG OD OPC
E
i
Genes (n=1467)
AST
MIG
OD
OPC
Clust.
Log2FC
2
-2
ii
GO Enrichment (GO:BP)
generation of neurons
neurogenesis
signal transduction
response to chemical
immune system process
regulation of immune system
neuron development
Schwann cell differentiation
peripheral nervous system develop.
cell projection organization
transport
establishment of localization
amide biosynthetic process
peptide biosynthetic process
translation
signal transduction
nucleic acid metabolic process
regulation of gene expression
neurogenesis
cell development
neuron differentiation
generation of neurons
neurogenesis
signal transduction
0
20
40
log10(p value)

**Supplemental Figure S7: Glial cells exhibit ALS-specific transcriptional responses but not ALS-specific changes in cell abundance.**

**A**) UMAP plots for each glial cell type, colored by individual donors. Donor IDs are colored by clinical grouping (grey, controls; yellow, ALS; red, ALSci).

**B**) Number of glial cells per library, separated by ALS phenotype (dot color). Medians are indicated by black bars.

**C**) Proportion of glial cells in each library, separated by ALS phenotype. Medians are indicated by black bars.

**D**) Distribution of UMI counts per cell for each glial cell type, separated by ALS phenotype.

**E**) Differentially expressed genes for each glial cell type. Clusters were assigned based on hierarchical clustering (**i**). The top three GO biological process terms are shown for each cluster in (**ii**).

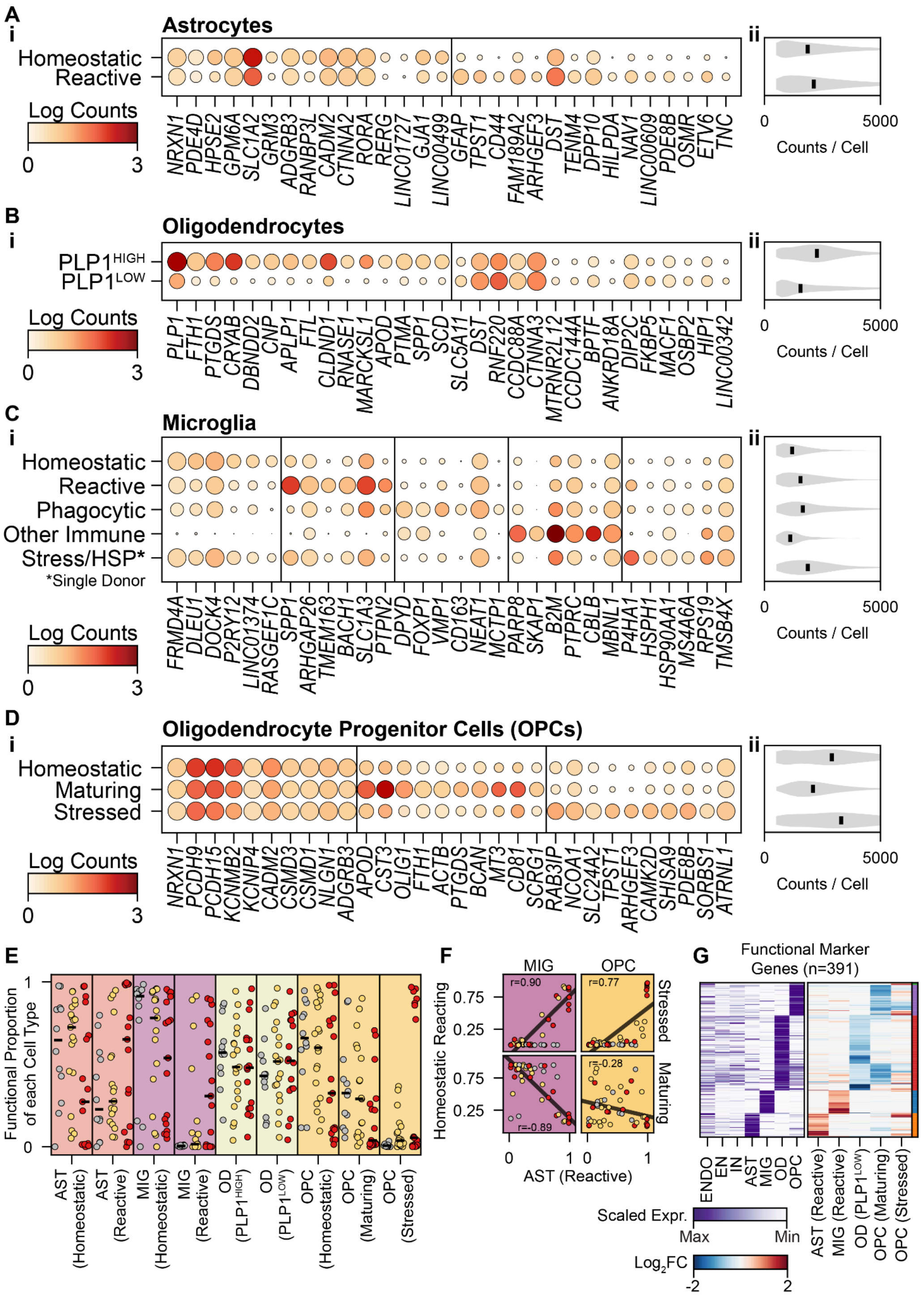
A
i
Astrocytes
Homeostatic
Reactive
Log Counts
0
3
NRXN1 PDE4D HPSE2 GPM6A SLC1A2 GRM3 ADGRB3 RANBP3L CADM2 CTNNA2 RORA RERG LINC01727 GJA1 LINC00499 GFAP TPST1 CD44 FAM189A2 ARHGEF3 DST TENM4 DPP10 HILPDA NAV1 LINC00609 PDE8B OSMR ETV6 TNC
ii
0
5000
Counts / Cell
B
i
Oligodendrocytes
PLP1HIGH
PLP1LOW
Log Counts
0
3
PLP1 FTH1 PTGDS CRYAB DBNDD2 CNP APLP1 FTL CLDND1 RNASE1 MARCKSL1 APOD PTMA SPP1 SCD SLC5A11 DST RNF220 CCDC88A CTNNA3 MTRNR2L12 CCDC144A BPTF ANKRD18A DIP2C FKBP5 MACF1 OSBP2 HIP1 LINC00342
ii
0
5000
Counts / Cell
C
i
Microglia
Homeostatic
Reactive
Phagocytic
Other Immune
Stress/HSP*
*Single Donor
Log Counts
0
3
FRMD4A DLEU1 DOCK4 P2RY12 LINC01374 RASGEF1C SPP1 ARHGAP26 TMEM163 BACH1 SLC1A3 PTPN2 DPYD FOXP1 VMP1 CD163 NEAT1 MCTP1 PARP8 SKAP1 B2M PTPRC CBLB MBNL1 P4HA1 HSPH1 HSP90AA1 MS4A6A RPS19 TMSB4X
ii
0
5000
Counts / Cell
D
i
Oligodendrocyte Progenitor Cells (OPCs)
Homeostatic
Maturing
Stressed
Log Counts
0
3
NRXN1 PCDH9 PCDH15 KCNMB2 KCNIP4 CADM2 CSMD3 CSMD1 NLGN1 ADGRB3 APOD CST3 OLIG1 FTH1 ACTB PTGDS BCAN MT3 CD81 SCRG1 RAB3IP NCOA1 SLC24A2 TPST1 ARHGEF3 CAMK2D SHISA9 PDE8B SORBS1 ATRNL1
ii
0
5000
Counts / Cell
E
Functional Proportion of each Cell Type
1
0
AST (Homeostatic)
AST (Reactive)
MIG (Homeostatic)
MIG (Reactive)
OD (PLP1HIGH)
OD (PLP1LOW)
OPC (Homeostatic)
OPC (Maturing)
OPC (Stressed)
F
MIG
OPC
r=0.90
r=0.77
r=-0.89
r=-0.28
Stressed
Maturing
Homeostatic Reacting
0.75
0.25
0.75
0.25
0
1
0
1
AST (Reactive)
G
Functional Marker Genes (n=391)
ENDO
EN
IN
AST
MIG
OD
OPC
AST (Reactive)
MIG (Reactive)
OD (PLP1LOW)
OPC (Maturing)
OPC (Stressed)
Scaled Expr.
Max
Min
Log2FC
-2
2

**Supplemental Figure S8: Glial cells are functionally separated based on transcriptional signatures.**

**A**) Expression of astrocyte-associated transcripts most significantly changed between functional categories (defined by scanpy as highest ranked by *t*-test *p*-values). Dot size indicates the percentage of cells in each functional category with non-zero counts (ranging from 0% to 100%), and color denotes log-transformed, standardized counts (**i**). Distribution of UMI counts per cell within each functional category (**ii**).

**B**) Expression of oligodendrocyte-associated transcripts, plotted as in panel Supplemental Figure S8A.

**C**) Expression of microglia-associated transcripts, plotted as in Supplemental Figure S8A.

**D**) Expression of oligodendrocyte progenitor cells-associated transcripts, plotted as in Supplemental Figure S8A.

**E**) Proportion of glial cells from each donor sample library annotated to specific functional categories. Black lines indicate the median across donor subgroups.

**F**) Proportion of homeostatic and reacting microglia plotted against the proportion of reactive astrocytes per library (left). Proportion of stressed and maturing OPCs plotted against the proportion of reactive astrocytes, using the same approach (right). All points are colored by ALS phenotype. Pearson correlation coefficients and least-squares regression lines are plotted.

**G**) Marker genes, which are differentially expressed based on glial cell functional category, were used to define functional scores in spatial data. Plots show min-max scaled expression per cell type (left), demonstrating cell type specificity, and $\log_2$ fold change values, denoting differential expression across (right).

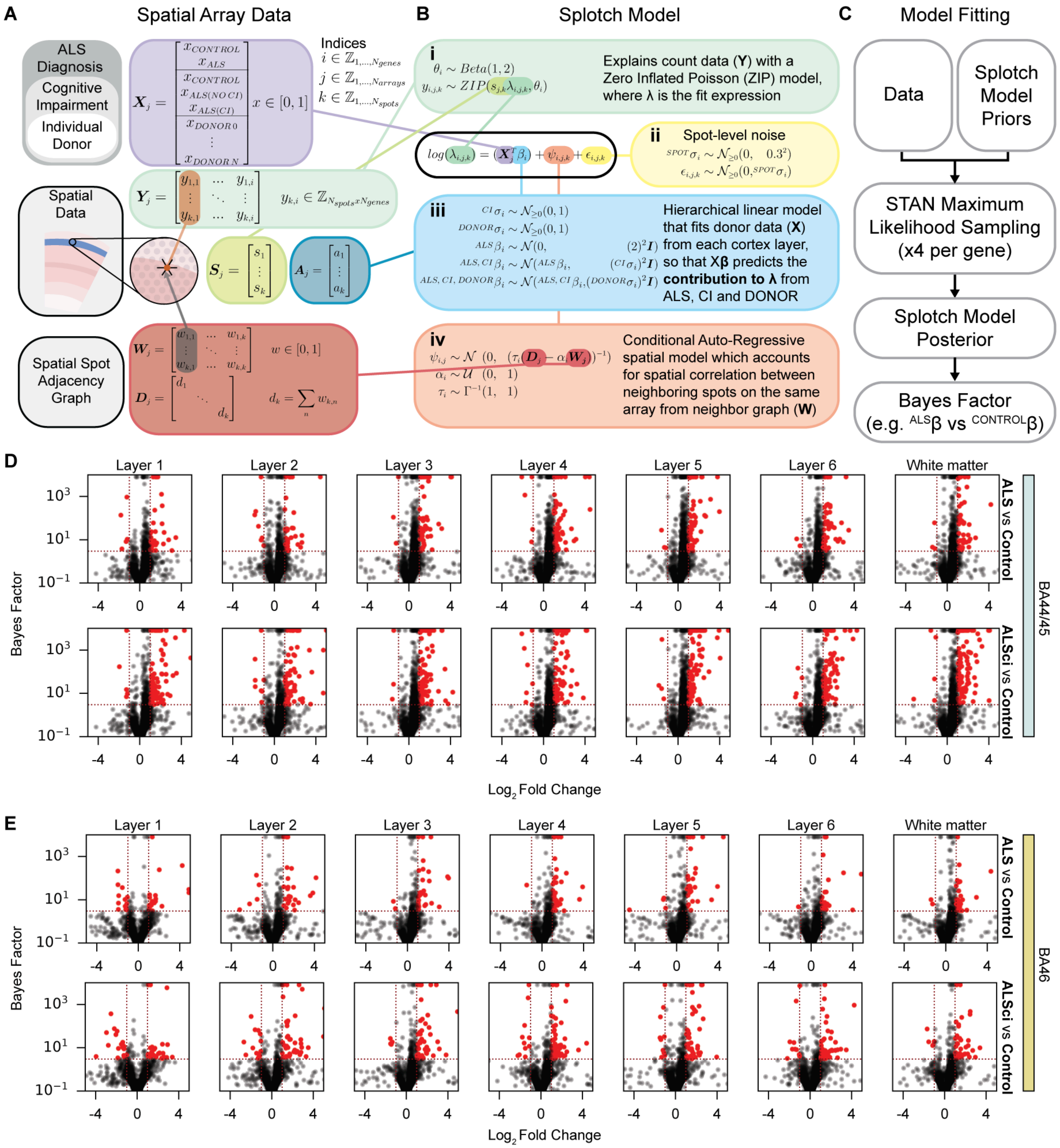
A
Spatial Array Data
ALS Diagnosis
Cognitive Impairment
Individual Donor
Indices
Spatial Data
Spatial Spot Adjacency Graph
B
Splotch Model
i
Explains count data (Y) with a Zero Inflated Poisson (ZIP) model, where λ is the fit expression
ii
Spot-level noise
iii
Hierarchical linear model that fits donor data (X) from each cortex layer, so that Xβ predicts the contribution to λ from ALS, CI and DONOR
iv
Conditional Auto-Regressive spatial model which accounts for spatial correlation between neighboring spots on the same array from neighbor graph (W)
C
Model Fitting
Data
Splotch Model Priors
STAN Maximum Likelihood Sampling (x4 per gene)
Splotch Model Posterior
Bayes Factor (e.g. ALSβ vs CONTROLβ)
D
Layer 1
Layer 2
Layer 3
Layer 4
Layer 5
Layer 6
White matter
Bayes Factor
ALS vs Control
ALSci vs Control
BA44/45
Log2 Fold Change
E
Layer 1
Layer 2
Layer 3
Layer 4
Layer 5
Layer 6
White matter
Bayes Factor
ALS vs Control
ALSci vs Control
BA46
Log2 Fold Change

**Supplemental Figure S9: Bayesian modeling of ST arrays using Splotch to generate posterior probabilities of gene expression and Bayes factors for differential gene expression testing.**

**A**) Schematic illustration of donor metadata ($\mathbf{X}_j$), spatial array count data ($\mathbf{Y}_j$), spatial array size factors ($\mathbf{S}_j$), cortex layer annotated anatomical region ($\mathbf{A}_j$), and spatial array adjacency matrix ($\mathbf{W}_j$) for a Visium transcriptomic array *j*.

**B**) Schematic illustration of the splotch model, separated into (**i**) modeling observed counts from (**ii**) individual-spot noise, (**iii**) a linear model that takes ALS disease state and donor identity as predictors, and (**iv**) a conditional auto-regressive spatial model that incorporates array spot adjacencies.

**C**) Schematic illustration of model fitting using STAN to obtain posterior probabilities for model parameters conditioned on priors and observed data.

**D**) Bayes factor volcano plots comparing ALS to Control and ALSci to Control for BA44/45 samples, with significantly changing defined as BF ≥ 3 and $|\log_2(FC)| \geq 1$, and with significantly changing genes highlighted in red.

**E**) Bayes factor volcano plots for BA46 samples, using the same criteria as in Supplemental Figure S9D.

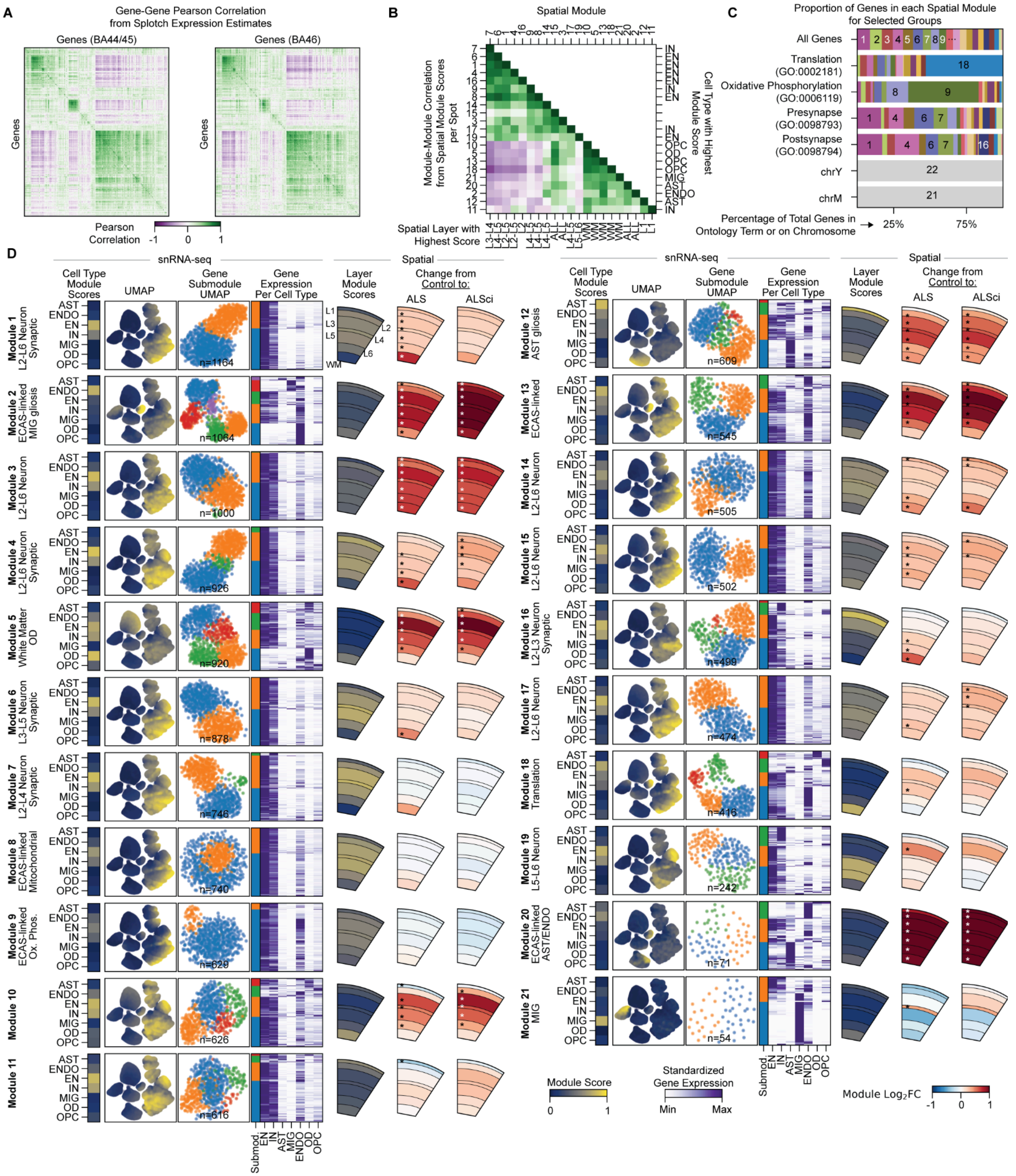
A
Gene-Gene Pearson Correlation from Splotch Expression Estimates
Genes (BA44/45)
Genes (BA46)
Genes
Pearson Correlation
-1 0 1
B
Spatial Module
Module-Module Correlation from Spatial Module Scores per Spot
Cell Type with Highest Module Score
Spatial Layer with Highest Score
C
Proportion of Genes in each Spatial Module for Selected Groups
All Genes
Translation (GO:0002181)
Oxidative Phosphorylation (GO:0006119)
Presynapse (GO:0098793)
Postsynapse (GO:0098794)
chrY
chrM
Percentage of Total Genes in Ontology Term or on Chromosome
25%
75%
D
snRNA-seq
Spatial
Cell Type Module Scores
UMAP
Gene Submodule UMAP
Gene Expression Per Cell Type
Layer Module Scores
Change from Control to:
ALS
ALSci
Module 1 L2-L6 Neuron Synaptic
Module 2 ECAS-linked MIG gliosis
Module 3 L2-L6 Neuron
Module 4 L2-L6 Neuron Synaptic
Module 5 White Matter OD
Module 6 L3-L5 Neuron Synaptic
Module 7 L2-L4 Neuron Synaptic
Module 8 ECAS-linked Mitochondrial
Module 9 ECAS-linked Ox. Phos.
Module 10
Module 11
Module 12 AST gliosis
Module 13 ECAS-linked
Module 14 L2-L6 Neuron
Module 15 L2-L6 Neuron
Module 16 L2-L3 Neuron Synaptic
Module 17 L2-L6 Neuron
Module 18 Translation
Module 19 L5-L6 Neuron
Module 20 ECAS-linked AST/ENDO
Module 21 MIG
Module Score
Standardized Gene Expression
Min
Max
Module $Log_2FC$

**Supplemental Figure S10: Spatial modules are associated with cell types, spatial location, and ALS phenotype.**

**A**) Pearson correlation between Splotch model estimates of spatial expression in BA44/45 (left) and BA46 (right).

**B**) Pearson correlation between spatial module scores across all ST samples. The right axis is annotated with the cell type that has the highest relative expression for each module. The bottom axis is annotated with the spatial layer with the highest relative expression of the module.

**C**) Top to bottom, the proportion of genes in each spatial module is plotted: 1) for all genes, 2) genes annotated with four selected GO terms, 3) genes on chromosome Y (chrY), and 4) genes on the mitochondrial chromosome (chrM).

**D**) Overview of each spatial module, presented from left to right: 1) Module number and brief summary; 2) average module score for each snRNA-seq-defined cell type; 3) snRNA-seq UMAP plot, where each point represents an individual cell colored by module score; 4) gene UMAP plot generated for all genes of the module from a *k*-NN graph (10 neighbors, snRNA-seq correlation distance metric), where each point is an individual gene colored by submodule assignment; 5) average standardized gene expression for each snRNA-seq cell type, with submodule assignment annotated; 6) Average ST module scores stratified by cortical layer; **7**) $Log_2$ fold change in module scores comparing ALS or ALSci donors to controls.

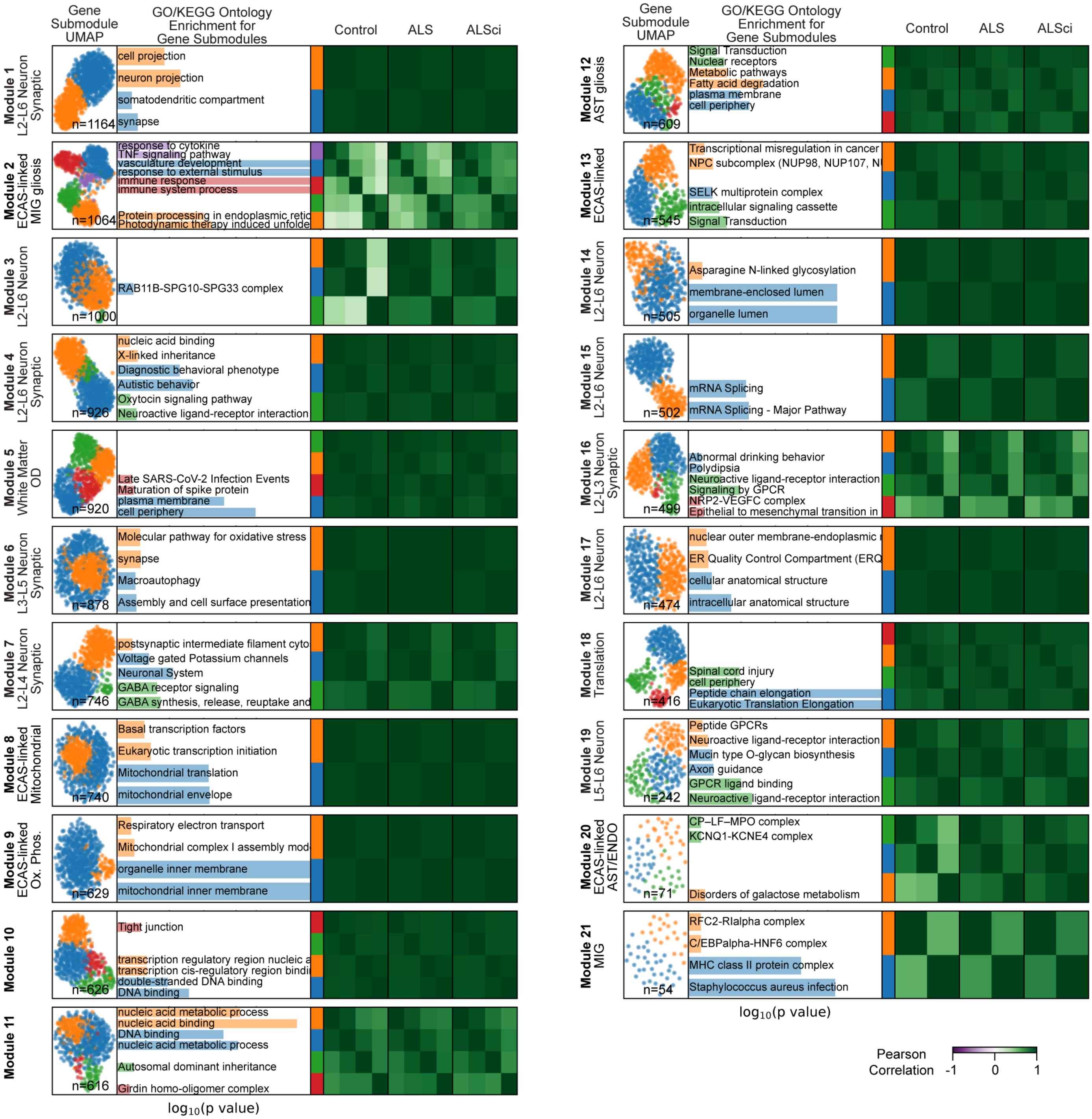
Gene Submodule UMAP
GO/KEGG Ontology Enrichment for Gene Submodules
Control
ALS
ALSci
Module 1 L2-L6 Neuron Synaptic
n=1164
cell projection
neuron projection
somatodendritic compartment
synapse
Module 2 ECAS-linked MIG gliosis
n=1064
response to cytokine
TNF signaling pathway
vasculature development
response to external stimulus
immune response
immune system process
Protein processing in endoplasmic retic
Photodynamic therapy induced unfolde
Module 3 L2-L6 Neuron
n=1000
RAB11B-SPG10-SPG33 complex
Module 4 L2-L6 Neuron Synaptic
n=926
nucleic acid binding
X-linked inheritance
Diagnostic behavioral phenotype
Autistic behavior
Oxytocin signaling pathway
Neuroactive ligand-receptor interaction
Module 5 White Matter OD
n=920
Late SARS-CoV-2 Infection Events
Maturation of spike protein
plasma membrane
cell periphery
Module 6 L3-L5 Neuron Synaptic
n=878
Molecular pathway for oxidative stress
synapse
Macroautophagy
Assembly and cell surface presentation
Module 7 L2-L4 Neuron Synaptic
n=746
postsynaptic intermediate filament cyto
Voltage gated Potassium channels
Neuronal System
GABA receptor signaling
GABA synthesis, release, reuptake and
Module 8 ECAS-linked Mitochondrial
n=740
Basal transcription factors
Eukaryotic transcription initiation
Mitochondrial translation
mitochondrial envelope
Module 9 ECAS-linked Ox. Phos.
n=629
Respiratory electron transport
Mitochondrial complex I assembly mod
organelle inner membrane
mitochondrial inner membrane
Module 10
n=626
Tight junction
transcription regulatory region nucleic a
transcription cis-regulatory region bindi
double-stranded DNA binding
DNA binding
Module 11
n=616
nucleic acid metabolic process
nucleic acid binding
DNA binding
nucleic acid metabolic process
Autosomal dominant inheritance
Girdin homo-oligomer complex
log10(p value)
Module 12 AST gliosis
n=609
Signal Transduction
Nuclear receptors
Metabolic pathways
Fatty acid degradation
plasma membrane
cell periphery
Module 13 ECAS-linked
n=545
Transcriptional misregulation in cancer
NPC subcomplex (NUP98, NUP107, N
SELK multiprotein complex
intracellular signaling cassette
Signal Transduction
Module 14 L2-L6 Neuron
n=505
Asparagine N-linked glycosylation
membrane-enclosed lumen
organelle lumen
Module 15 L2-L6 Neuron
n=502
mRNA Splicing
mRNA Splicing - Major Pathway
Module 16 L2-L3 Neuron Synaptic
n=499
Abnormal drinking behavior
Polydipsia
Neuroactive ligand-receptor interaction
Signaling by GPCR
NRP2-VEGFC complex
Epithelial to mesenchymal transition in
Module 17 L2-L6 Neuron
n=474
nuclear outer membrane-endoplasmic r
ER Quality Control Compartment (ERQ
cellular anatomical structure
intracellular anatomical structure
Module 18 Translation
n=416
Spinal cord injury
cell periphery
Peptide chain elongation
Eukaryotic Translation Elongation
Module 19 L5-L6 Neuron
n=242
Peptide GPCRs
Neuroactive ligand-receptor interaction
Mucin type O-glycan biosynthesis
Axon guidance
GPCR ligand binding
Neuroactive ligand-receptor interaction
Module 20 ECAS-linked AST/ENDO
n=71
CP–LF–MPO complex
KCNQ1-KCNE4 complex
Disorders of galactose metabolism
Module 21 MIG
n=54
RFC2-RIalpha complex
C/EBPalpha-HNF6 complex
MHC class II protein complex
Staphylococcus aureus infection
log10(p value)
Pearson Correlation
-1
0
1

**Supplemental Figure S11: Separating spatial gene modules into cell type submodules using snRNA-seq transcriptome correlations.**

For each module, panels are displayed from left to right using this logic: 1) Module number and brief summary; 2) snRNA-seq UMAP plot, where each point represents an individual cell colored by module score; 3) GO or KEGG term enriched in the submodule, accompanied by a bar plot colored by submodule identity and where bar length denotes $-\log_{10}$($p$-value); 4) Pearson correlation coefficients between submodule scores in ST data, stratified by donor group (control, ALS, and ALSci).

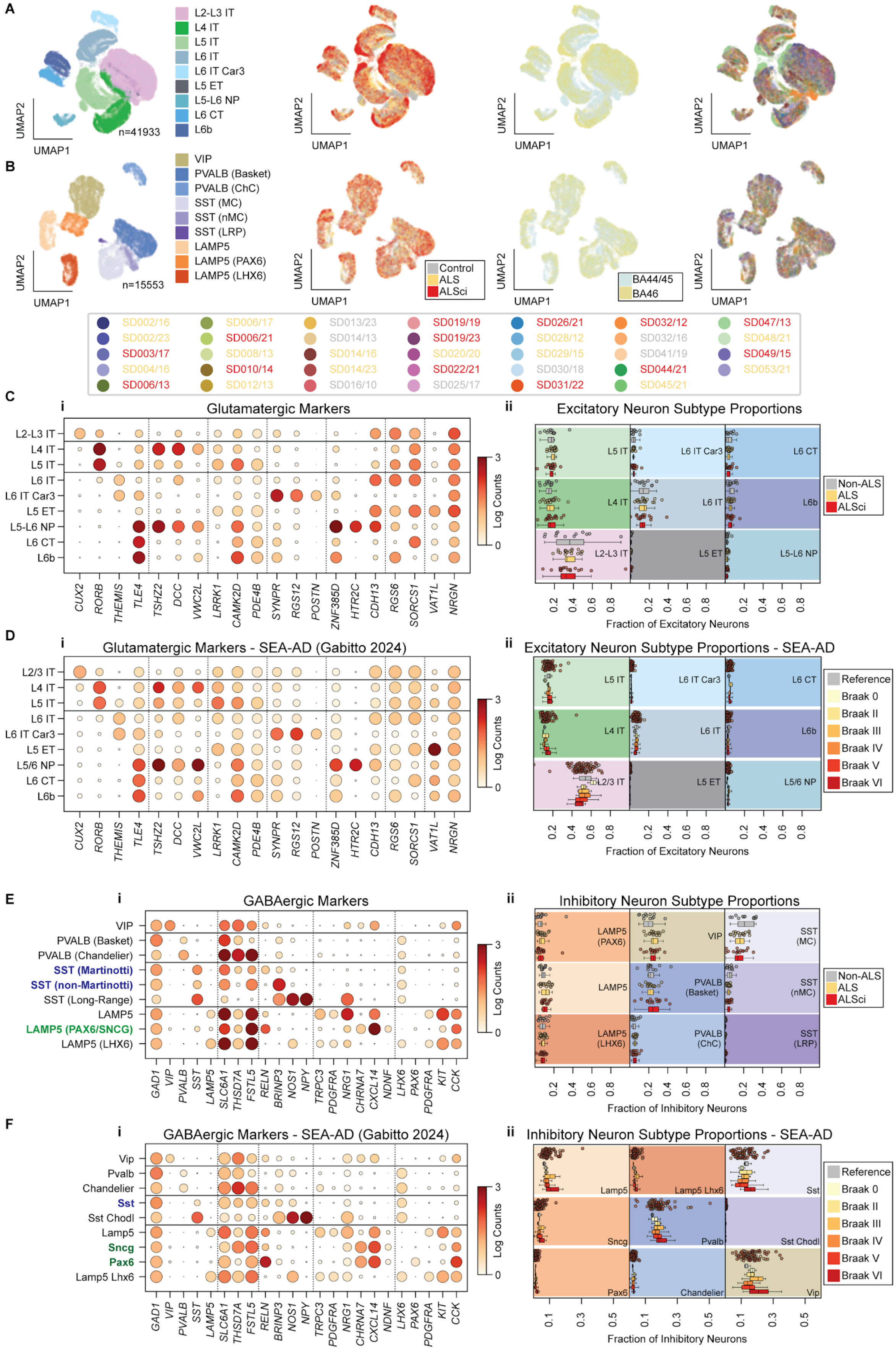
A
L2-L3 IT
L4 IT
L5 IT
L6 IT
L6 IT Car3
L5 ET
L5-L6 NP
L6 CT
L6b
n=41933
UMAP1
UMAP2
B
VIP
PVALB (Basket)
PVALB (ChC)
SST (MC)
SST (nMC)
SST (LRP)
LAMP5
LAMP5 (PAX6)
LAMP5 (LHX6)
n=15553
Control
ALS
ALSci
BA44/45
BA46
SD002/16 SD006/17 SD013/23 SD019/19 SD026/21 SD032/12 SD047/13
SD002/23 SD006/21 SD014/13 SD019/23 SD028/12 SD032/16 SD048/21
SD003/17 SD008/13 SD014/16 SD020/20 SD029/15 SD041/19 SD049/15
SD004/16 SD010/14 SD014/23 SD022/21 SD030/18 SD044/21 SD053/21
SD006/13 SD012/13 SD016/10 SD025/17 SD031/22 SD045/21
C
i Glutamatergic Markers
ii Excitatory Neuron Subtype Proportions
Log Counts
Non-ALS
ALS
ALSci
Fraction of Excitatory Neurons
D
i Glutamatergic Markers - SEA-AD (Gabitto 2024)
ii Excitatory Neuron Subtype Proportions - SEA-AD
Reference
Braak 0
Braak II
Braak III
Braak IV
Braak V
Braak VI
Fraction of Excitatory Neurons
CUX2 RORB THEMIS TLE4 TSHZ2 DCC VWC2L LRRK1 CAMK2D PDE4B SYNPR RGS12 POSTN ZNF385D HTR2C CDH13 RGS6 SORCS1 VAT1L NRGN
E
i GABAergic Markers
ii Inhibitory Neuron Subtype Proportions
VIP
PVALB (Basket)
PVALB (Chandelier)
SST (Martinotti)
SST (non-Martinotti)
SST (Long-Range)
LAMP5
LAMP5 (PAX6/SNCG)
LAMP5 (LHX6)
Fraction of Inhibitory Neurons
F
i GABAergic Markers - SEA-AD (Gabitto 2024)
ii Inhibitory Neuron Subtype Proportions - SEA-AD
Vip
Pvalb
Chandelier
Sst
Sst Chodl
Lamp5
Sncg
Pax6
Lamp5 Lhx6
GAD1 VIP PVALB SST LAMP5 SLC6A1 THSD7A FSTL5 RELN BRINP3 NOS1 NPY TRPC3 PDGFRA NRG1 CHRNA7 CXCL14 NDNF LHX6 PAX6 PDGFRA KIT CCK
Fraction of Inhibitory Neurons

**Supplemental Figure S12: Neuronal subtype classification and comparison with published data.**

**A**) UMAP plot of excitatory neurons, colored by neuronal subtype, ALS disease phenotype, brain region, and individual donor. Donor legend labels are colored by ALS disease phenotype**.**

**B**) UMAP plot of inhibitory neurons colored as in Supplemental Figure S13A.

**C**) Glutamatergic (EN) neuronal subtype marker genes, colored by log normalized expression and dot size indicating the proportion of cells with nonzero counts (i). The proportion of each subtype relative to the total number of glutamatergic neurons per donor, separated by ALS disease phenotype, is shown in panel (ii).

**D**) Glutamatergic (EN) subtype annotations from an Alzheimer's disease cohort (Gabitto et al., Reference 40). It includes marker genes (i) and the proportion of each subtype relative to total number of glutamatergic neurons per donor, stratified by Braak score (ii).

**E**) GABAergic (IN) neuronal subtypes in ALS cohort, plotted as in Supplemental Figure S13C.

**F)** GABAergic (IN) neuronal subtypes plotted as in Supplemental Figure S13B for Alzheimer's cohort. SST (Martinotti) and SST (non-Martinotti) in this cohort are a single SST annotation in Gabitto et al (Reference 40), and *LAMP5*$^{+}$ (*PAX6*$^{+}$) is a single annotation in this cohort but separate Sncg and Pax6 annotations in Gabitto et al.

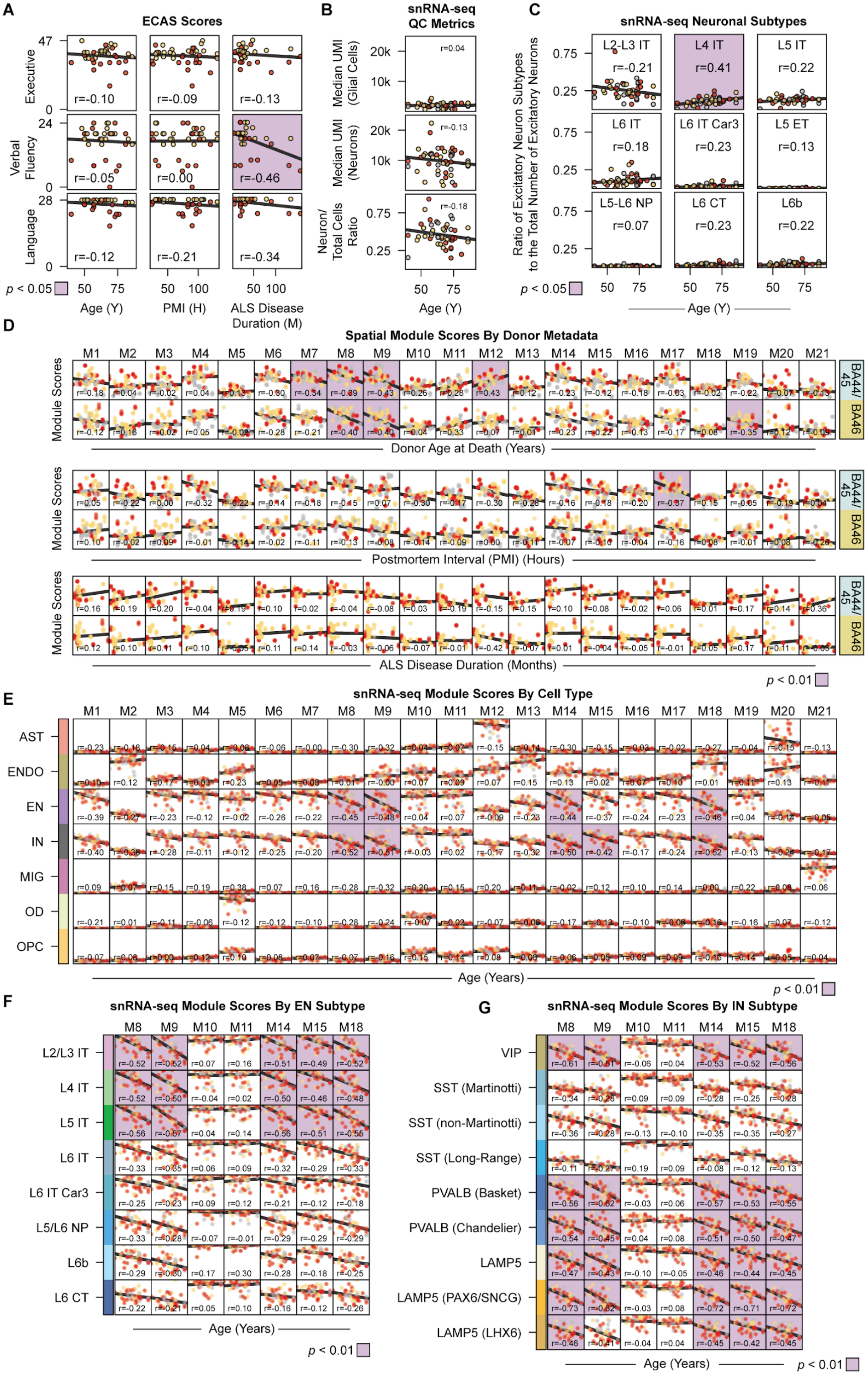
A
ECAS Scores
Executive
Verbal Fluency
Language
r=-0.10
r=-0.09
r=-0.13
r=-0.05
r=0.00
r=-0.46
r=-0.12
r=-0.21
r=-0.34
p < 0.05
Age (Y)
PMI (H)
ALS Disease Duration (M)
B
snRNA-seq QC Metrics
Median UMI (Glial Cells)
Median UMI (Neurons)
Neuron/ Total Cells Ratio
r=0.04
r=-0.13
r=-0.18
Age (Y)
C
snRNA-seq Neuronal Subtypes
Ratio of Excitatory Neuron Subtypes to the Total Number of Excitatory Neurons
L2-L3 IT
r=-0.21
L4 IT
r=0.41
L5 IT
r=0.22
L6 IT
r=0.18
L6 IT Car3
r=0.23
L5 ET
r=0.13
L5-L6 NP
r=0.07
L6 CT
r=0.23
L6b
r=0.22
p < 0.05
Age (Y)
D
Spatial Module Scores By Donor Metadata
Module Scores
BA44/45
BA46
Donor Age at Death (Years)
Postmortem Interval (PMI) (Hours)
ALS Disease Duration (Months)
p < 0.01
E
snRNA-seq Module Scores By Cell Type
AST
ENDO
EN
IN
MIG
OD
OPC
Age (Years)
p < 0.01
F
snRNA-seq Module Scores By EN Subtype
L2/L3 IT
L4 IT
L5 IT
L6 IT
L6 IT Car3
L5/L6 NP
L6b
L6 CT
Age (Years)
p < 0.01
G
snRNA-seq Module Scores By IN Subtype
VIP
SST (Martinotti)
SST (non-Martinotti)
SST (Long-Range)
PVALB (Basket)
PVALB (Chandelier)
LAMP5
LAMP5 (PAX6/SNCG)
LAMP5 (LHX6)
Age (Years)
p < 0.01

**Supplemental Figure S13: Neuronal Transcription is Affected by Age.**

**A**) ECAS Executive, Verbal Fluency, and Language scores plotted against donor age at death, postmortem interval (PMI), and ALS disease duration. Non-neuropathological controls are plotted at the maximum ECAS score, although they did not undergo cognitive screening. Least-squares regression lines and Pearson correlation coefficients are shown for ALS and ALSci donors. Plots with correlations significantly different from zero (Wald test, $p < 0.05$) are shaded in lilac.

**B**) Median UMI counts per donor sample for glia, neurons, and neuron-to-glia ratios plotted against donor age. Plots with correlations significantly different from zero (Wald test, $p < 0.05$) would be shaded in lilac, but none are significant.

**C**) Proportion of each glutaminergic neuronal subtype relative to the total number of glutamatergic neurons per donor, plotted against donor age. Plots with correlations significantly different from zero (Wald test, $p < 0.05$) are shaded in lilac.

**D**) Spatial module scores from ST data, averaged across all cortex layers for each donor sample, plotted on the y-axis, compared to donor age, PMI, and ALS disease duration plotted on the x-axis. Panels highlighted in lilac have a statistically significant non-zero linear relationship by Benjamini-Hochberg corrected Wald test ($p < 0.01$).

**E**) snRNA-seq module scores stratified by cell type, averaged for each donor sample, and plotted against donor age on the x-axis. Panels highlighted in lilac have a statistically significant non-zero linear relationship by Benjamini-Hochberg corrected Wald test ($p < 0.01$).

**F**) snRNA-seq module scores stratified by EN neuronal subtype, averaged for each donor sample, and plotted against donor age on the x-axis. The five significant modules highlighted in E are shown, along with M10 and M11 for comparison. Panels highlighted in lilac have a statistically significant non-zero linear relationship by Benjamini-Hochberg corrected Wald test ($p < 0.01$).

**G**)  snRNA-seq module scores stratified by IN neuronal subtype, plotted and tested as in plotted as in Supplemental Figure S14F.

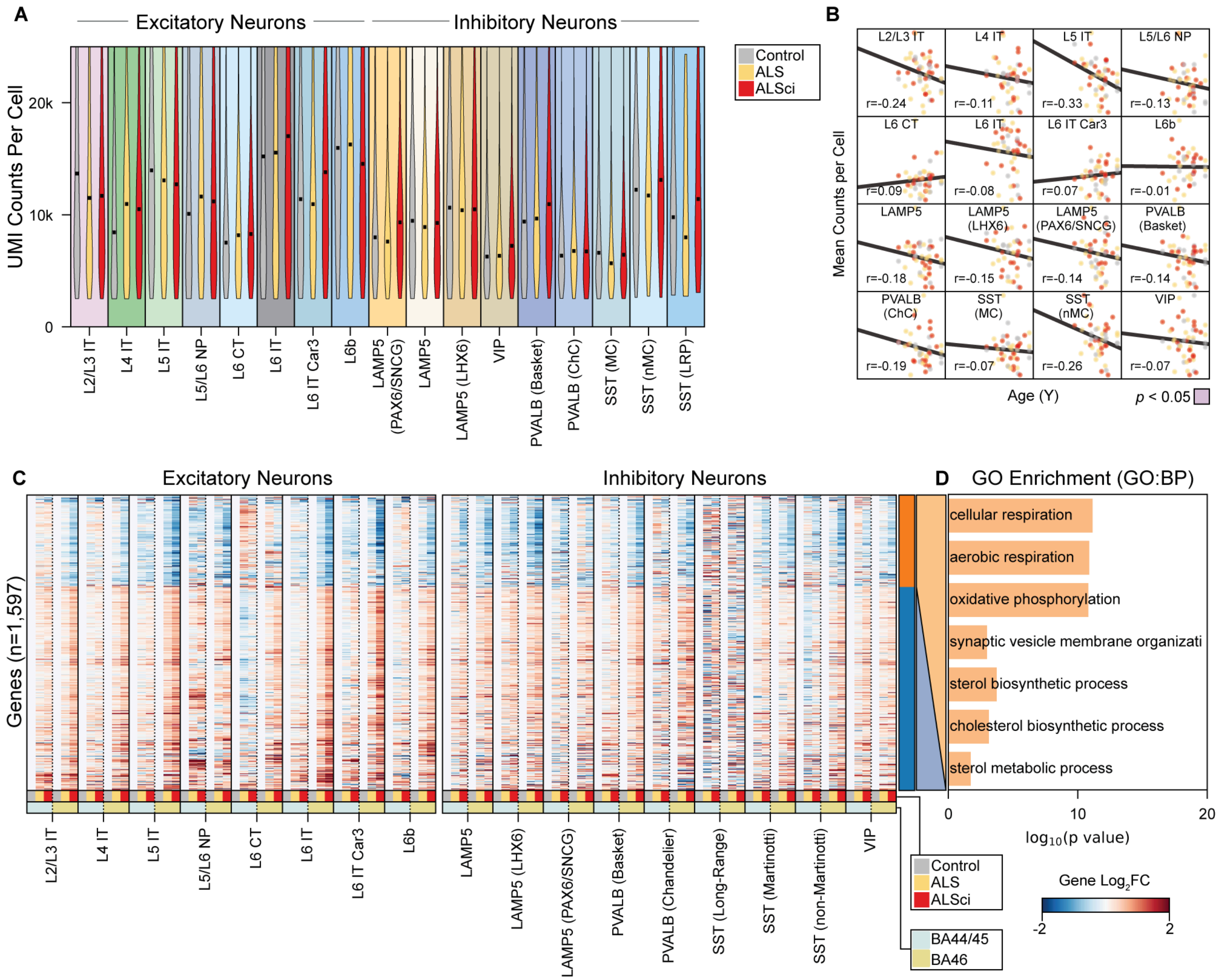
A
Excitatory Neurons
Inhibitory Neurons
UMI Counts Per Cell
Control
ALS
ALSci
L2/L3 IT
L4 IT
L5 IT
L5/L6 NP
L6 CT
L6 IT
L6 IT Car3
L6b
LAMP5 (PAX6/SNCG)
LAMP5
LAMP5 (LHX6)
VIP
PVALB (Basket)
PVALB (ChC)
SST (MC)
SST (nMC)
SST (LRP)
B
Mean Counts per Cell
Age (Y)
p < 0.05
C
Excitatory Neurons
Inhibitory Neurons
Genes (n=1,597)
BA44/45
BA46
Gene Log2FC
D
GO Enrichment (GO:BP)
cellular respiration
aerobic respiration
oxidative phosphorylation
synaptic vesicle membrane organizati
sterol biosynthetic process
cholesterol biosynthetic process
sterol metabolic process
log10(p value)

**Supplemental Figure S14: Transcriptional differences in neuron subtypes between control donors and ALS or ALSci donors**

**A**) Violin plot of counts per cell in each neuronal subtype, stratified by Control, ALS, and ALSci. Black lines indicate median counts per cell.

**B**) Mean counts per cell for neuronal subtype in each donor plotted against donor age. Ordinary least squares regression line is plotted and tested for slope not equal to zero by Wald test. Statistically significant relationships are highlighted in lilac (all are non-significant).

**C**) $\mathrm{Log}_2$ Fold Change heatmap for 1,597 genes which are significantly different (DESeq2; Multiple hypothesis test corrected Wald test; $p < 0.05$) in one cell type when comparing to control.

**D**) $-\mathrm{Log}_{10}$($p$-value) of selected GO:BP terms significantly enriched in the orange cluster. The blue cluster does not have significantly enriched GO:BP terms.

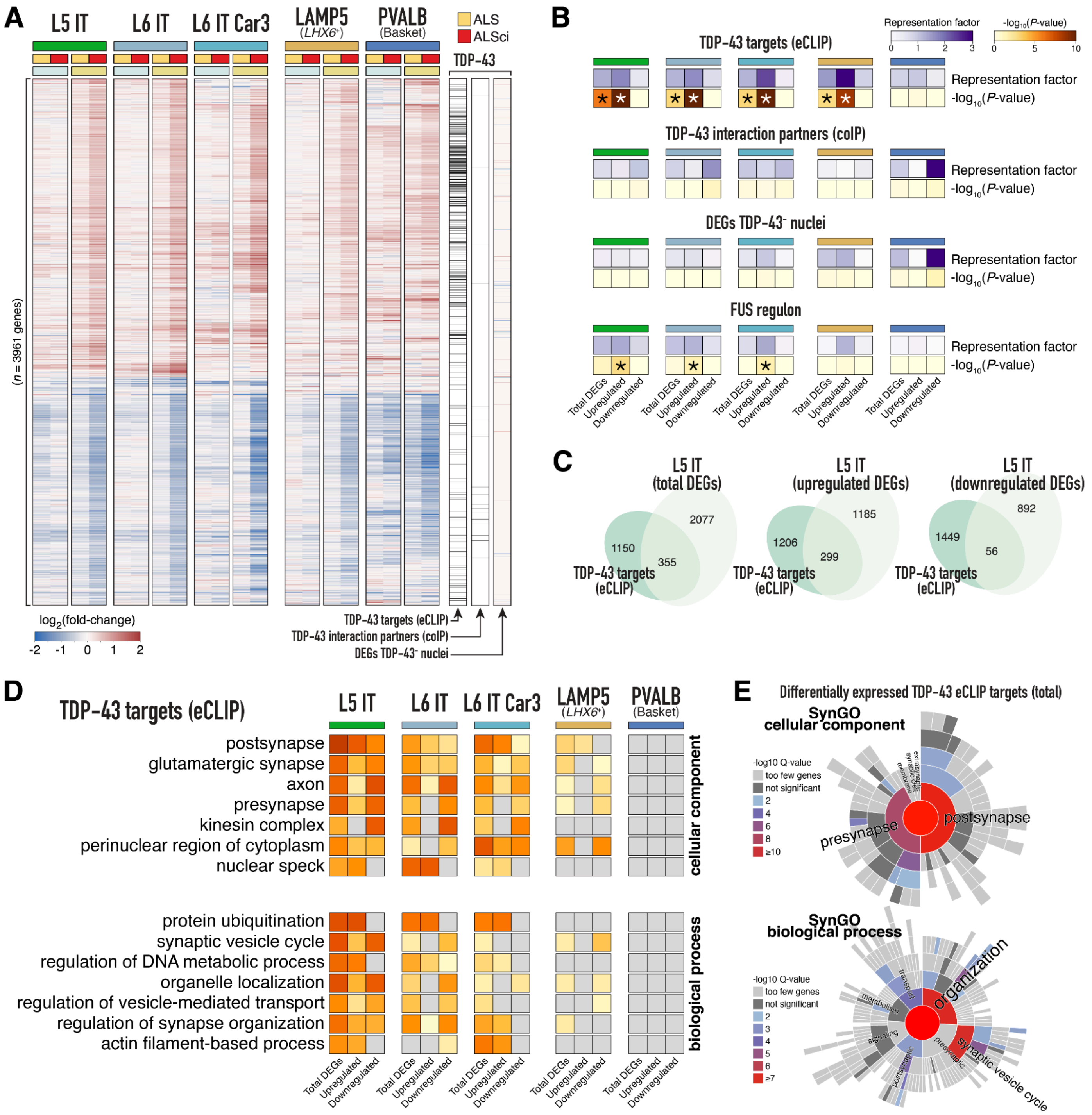
A
L5 IT
L6 IT
L6 IT Car3
LAMP5
(LHX6+)
PVALB
(Basket)
ALS
ALSci
TDP-43
(n = 3961 genes)
log2(fold-change)
-2 -1 0 1 2
TDP-43 targets (eCLIP)
TDP-43 interaction partners (coIP)
DEGs TDP-43- nuclei
B
Representation factor
0 1 2 3
-log10(P-value)
0 5 10
TDP-43 targets (eCLIP)
Representation factor
-log10(P-value)
TDP-43 interaction partners (coIP)
DEGs TDP-43- nuclei
FUS regulon
Total DEGs
Upregulated
Downregulated
C
L5 IT
(total DEGs)
2077
1150
355
TDP-43 targets
(eCLIP)
L5 IT
(upregulated DEGs)
1185
1206
299
L5 IT
(downregulated DEGs)
892
1449
56
D
TDP-43 targets (eCLIP)
L5 IT
L6 IT
L6 IT Car3
LAMP5
(LHX6+)
PVALB
(Basket)
postsynapse
glutamatergic synapse
axon
presynapse
kinesin complex
perinuclear region of cytoplasm
nuclear speck
cellular component
protein ubiquitination
synaptic vesicle cycle
regulation of DNA metabolic process
organelle localization
regulation of vesicle-mediated transport
regulation of synapse organization
actin filament-based process
biological process
Total DEGs
Upregulated
Downregulated
E
Differentially expressed TDP-43 eCLIP targets (total)
SynGO
cellular component
-log10 Q-value
too few genes
not significant
2
4
6
8
≥10
presynapse
postsynapse
SynGO
biological process
-log10 Q-value
too few genes
not significant
2
3
4
5
6
≥7
organization
transport
metabolism
signaling
synaptic vesicle cycle

**Supplemental Figure 15: Synaptic enrichment of eCLIP-defined TDP-43 targets in ALSci.**

**A**) Heatmap showing $log_2$-transformed gene expression levels (fold-change versus control) for genes differentially expressed in at least one of five neuronal subtypes. Additional annotations indicate TDP-43 binding transcripts defined by eCLIP-seq analyses, transcripts encoding for proteins that physically interact with TDP-43, and differentially expressed transcripts upon TDP-43 nuclear depletion in postmortem FTD-ALS brains.

**B**) Representation factor analysis of TDP-43-associated gene sets in neuronal subtypes, performed using ALSci nuclei from BA46. Asterisks denote significant representation. Results are shown for total, upregulated, and downregulated differentially expressed genes.

**C**) Venn diagrams showing the overlap between eCLIP-defined TDP-43 targets and differentially expressed genes in L5 IT neurons from ALSci BA46 samples. Results are shown for total, upregulated, and downregulated differentially expressed genes.

**D**) Gene ontology (GO) meta-analysis using Metascape of differentially expressed genes across neuronal subtypes from ALSci BA46 samples. Results are shown for total, upregulated, and downregulated differentially expressed genes.

**E**) GO analysis using SynGO of TDP-43 target genes (defined by eCLIP-seq) differentially expressed in L5 IT neurons from ALSci BA46 samples.

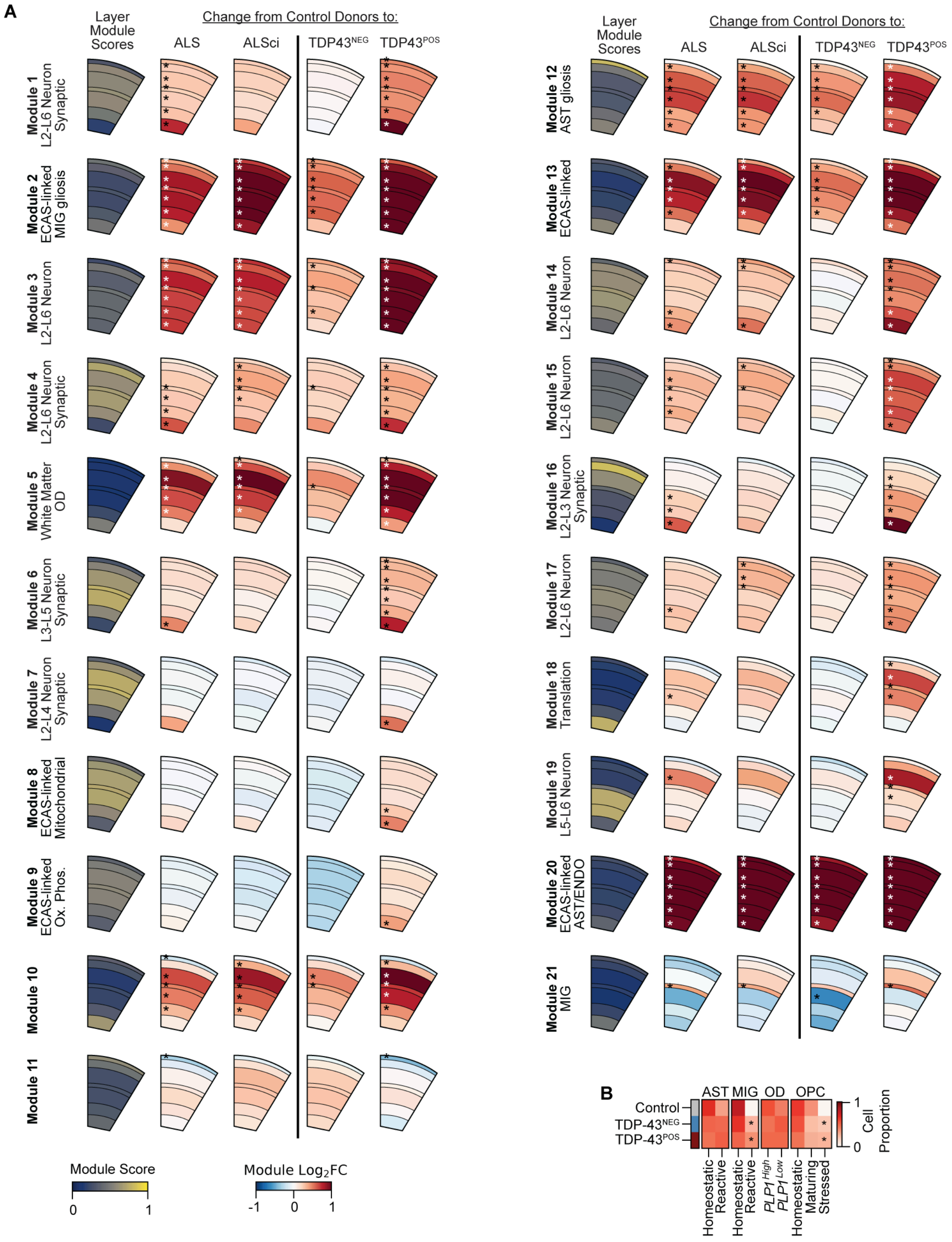
A
Layer Module Scores
Change from Control Donors to:
ALS
ALSci
TDP43NEG
TDP43POS
Module 1 L2-L6 Neuron Synaptic
Module 2 ECAS-linked MIG gliosis
Module 3 L2-L6 Neuron
Module 4 L2-L6 Neuron Synaptic
Module 5 White Matter OD
Module 6 L3-L5 Neuron Synaptic
Module 7 L2-L4 Neuron Synaptic
Module 8 ECAS-linked Mitochondrial
Module 9 ECAS-linked Ox. Phos.
Module 10
Module 11
Module 12 AST gliosis
Module 13 ECAS-linked
Module 14 L2-L6 Neuron
Module 15 L2-L6 Neuron
Module 16 L2-L3 Neuron Synaptic
Module 17 L2-L6 Neuron
Module 18 Translation
Module 19 L5-L6 Neuron
Module 20 ECAS-linked AST/ENDO
Module 21 MIG
Module Score
0
1
Module Log2FC
-1
0
1
B
AST MIG OD OPC
Control
TDP-43NEG
TDP-43POS
Homeostatic
Reactive
Homeostatic
Reactive
PLP1High
PLP1Low
Homeostatic
Maturing
Stressed
1
0
Cell Proportion

**Supplemental Figure S16: Neuronal spatial gene module upregulation in ALS/ALSci donors is linked to TDP-43 aggregation.**

**A**) For each module, the average score of the spatial gene module from spatial expression data is plotted for each cortex layer. Differential module expression is plotted from spatial expression data, comparing control donors to ALS and ALSci donors, and comparing control donors to donors negative for TDP-43 aggregation (TDP-43$^{NEG}$) and donors positive for TDP-43 aggregation (TDP-43$^{POS}$). Asterisks indicate significant differences for a cortex layer, tested by Benjaminini-Yekutieli controlled Welch's *t*-test ($p < 0.01$).

**B**) Averaged proportions of glial cell functional states across controls, TDP-43$^{NEG}$, and TDP-43$^{POS}$ donors. Asterisks denote significant differences compared to controls (Welch's *t*-test, $p < 0.05$).

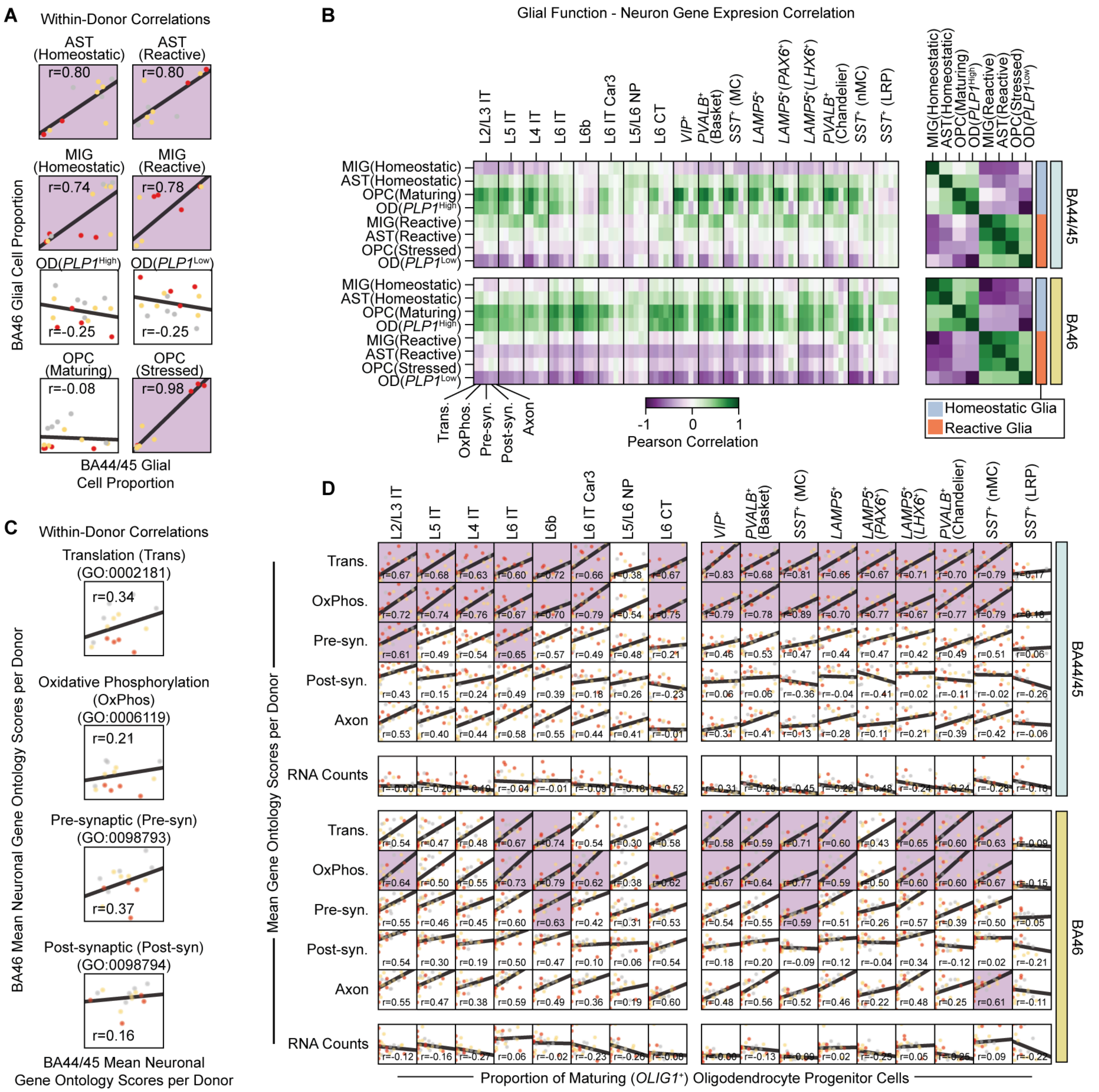
A
Within-Donor Correlations
AST (Homeostatic)
r=0.80
AST (Reactive)
r=0.80
MIG (Homeostatic)
r=0.74
MIG (Reactive)
r=0.78
OD(PLP1High)
r=-0.25
OD(PLP1Low)
r=-0.25
OPC (Maturing)
r=-0.08
OPC (Stressed)
r=0.98
BA46 Glial Cell Proportion
BA44/45 Glial Cell Proportion
B
Glial Function - Neuron Gene Expresion Correlation
L2/L3 IT
L5 IT
L4 IT
L6 IT
L6b
L6 IT Car3
L5/L6 NP
L6 CT
VIP+
PVALB+ (Basket)
SST+ (MC)
LAMP5+
LAMP5+(PAX6+)
LAMP5+(LHX6+)
PVALB+ (Chandelier)
SST+ (nMC)
SST+ (LRP)
MIG(Homeostatic)
AST(Homeostatic)
OPC(Maturing)
OD(PLP1High)
MIG(Reactive)
AST(Reactive)
OPC(Stressed)
OD(PLP1Low)
BA44/45
BA46
Trans.
OxPhos.
Pre-syn.
Post-syn.
Axon
-1
0
1
Pearson Correlation
Homeostatic Glia
Reactive Glia
C
Within-Donor Correlations
Translation (Trans) (GO:0002181)
r=0.34
Oxidative Phosphorylation (OxPhos) (GO:0006119)
r=0.21
Pre-synaptic (Pre-syn) (GO:0098793)
r=0.37
Post-synaptic (Post-syn) (GO:0098794)
r=0.16
BA46 Mean Neuronal Gene Ontology Scores per Donor
BA44/45 Mean Neuronal Gene Ontology Scores per Donor
D
Mean Gene Ontology Scores per Donor
Trans.
OxPhos.
Pre-syn.
Post-syn.
Axon
RNA Counts
BA44/45
BA46
Proportion of Maturing (OLIG1+) Oligodendrocyte Progenitor Cells

**Supplemental Figure 17: Gliotic response mediated by astrocytes and microglia occurs in both BA44/45 and BA46 from the same donors, but neuronal dysfunction does not, and correlates with oligodendrocyte dysfunction.**

**A**) Within-donor correlation between BA44/45 and BA46 for the proportion of homeostatic and reactive astrocytes, homeostatic and reactive microglia, high- and low-myelinating oligodendrocytes, and maturing and stressed OPCs. Black line shows best-fit from ordinary least squares regression. Plots highlighted in lilac have best-fit lines with slopes significantly different from 0 (Wald test, $p < 0.05$).

**B**) Correlation between glial cell functional proportions and the aggregate snRNA-seq expression of neuronal transcripts annotated with specific ontologies, plotted for in BA44/45 (top) and BA46 (bottom). Correlation between glial cell functional proportions is shown on the right (right).

**C**) Within-donor correlation of neuronal expression profiles of genes annotated with specific ontologies between BA44/45 and BA46. Best-fit regression lines and statistical testing as in Supplemental Figure 17A; all comparisons are non-significant.

**D**) Correlation between the proportion  of maturing OPCs and aggregate gene expression in each neuronal subtype, plotted separately for BA44/45 and BA46. Best-fit lines and statistical testing as in Supplemental Figure 17A. Panels highlighted in lilac are significant by Benjamini-Hochberg corrected Wald test ($p < 0.01$). Average RNA counts per donor are also plotted to rule out count depth as a potential confounding factor.
